## Supplementary Info for "Enhancing Intracellular Accumulation and Target Engagement of PROTACs with Reversible Covalent Chemistry"

Guo *et al.*

### Supplementary Methods

### Compound Synthesis and Characterization

**Materials:** All chemicals were purchased from Sigma-Aldrich, Combi-blocks or Alfa Aesar, unless otherwise specified. All solvents and reagents were used as obtained without further purification.

**Instrumentation:** $^{1}$H NMR and $^{13}$C NMR spectra were on a Varian (Palo Alto, CA) 400-MR spectrometer. Chemical shifts (δ) are reported in ppm, and coupling constants (J) are in Hertz (Hz). The following abbreviations were used to explain the multiplicities: s = singlet, d = doublet, t = triplet, q = quartet, m = multiplet, br = broad. Flash chromatography was performed on a Teledyne ISCO CombiFlash Rf 200. ESI mass spectrometry was measured on an Agilent Mass Spectrometer.

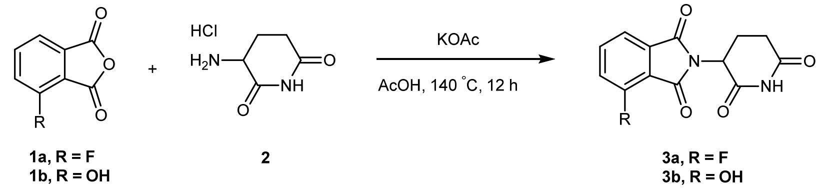


Compound **1a or 1b** (1 g, 6.1 mmol) and compound **2** (1g, 6.1 mmol) were dissolved in AcOH (20 mL) and heated to 140°C for overnight, the mixture was cooled to room temperature and concentrated under reduced pressure. The residue was purified by flash column chromatography (0-10% MeOH in DCM) to give compound **3a or 3b** as a white solid (1.3 g, 79%). Compound **3a**. $^{1}$H NMR (400 MHz, *D$_6$*-DMSO) δ 11.16 (s, 1H), 7.95 (m, 1H), 7.79 (d, *J* = 7.3 Hz, 1H), 7.74 (t, *J* = 8.9 Hz, 1H), 5.16 (dd, *J* = 12.8, 5.4 Hz, 1H), 2.89 (m, 1H), 2.57 (m, 2H), 2.07 (M, 1H). Compound **3b.** $^{1}$H NMR (400 MHz, *D$_6$*-DMSO) δ 11.15 (s, 1H), 11.06 (s, 1H), 7.63 (t, *J* = 7.8 Hz, 1H), 7.29 (d, *J* = 7.0 Hz, 1H), 7.22 (d, *J* = 8.3 Hz, 1H), 5.04 (dd, *J* = 12.9, 4.8 Hz, 1H), 2.86 (m, 1H), 2.54 (m, 2H), 1.99 (m, 1H).

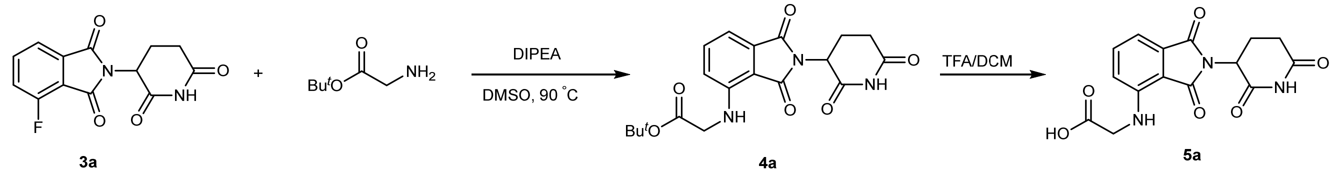


***tert*-butyl (2-(2,6-dioxopiperidin-3-yl)-1,3-dioxoisoindolin-4-yl)glycinate (4a)**. Compound **3a** (552 mg, 2 mmol), *tert*-butyl glycinate (524 mg, 4 mmol) and DIPEA (516 mg, 4 mmol) were dissolved in DMSO (10 mL) and heated to 90°C. After 4 hours, the mixture was cooled to room temperature and concentrated under reduced pressure. The residue was purified by flash column chromatography (0-10% MeOH in DCM) to give compound **4a** as a yellow solid (542 mg, 70%).$^{1}$H NMR (400 MHz, *D$_6$*-DMSO) δ 11.11 (s, 1H), 7.58 (t, *J* = 8.0 Hz, 1H), 7.08 (d, *J* = 7.1 Hz, 1H), 6.97 (d, *J* = 8.5 Hz, 1H), 6.85 (t, *J* = 6.0 Hz, 1H), 5.07 (dd, *J* = 13.0, 5.4 Hz, 1H), 4.09 (d, *J* = 6.0 Hz, 2H), 2.89 (m, 1H), 2.56 (m, 2H), 2.05 (m, 1H), 1.43 (s, 9H).

**(2-(2,6-Dioxopiperidin-3-yl)-1,3-dioxoisoindolin-4-yl)glycine (5a)**. In a 100 mL flask was added compound **4a** (542 mg, 1.4 mmol) in TFA/DCM (10 mL, 1/1). The mixture was stirred for 30 min at room temperature. LC-

MS showed **4a** converted into **5a** completely. Then remove the solvent *in vacuo* to give product **5a** (440 mg, 95%), which was used for for next step without further purification.

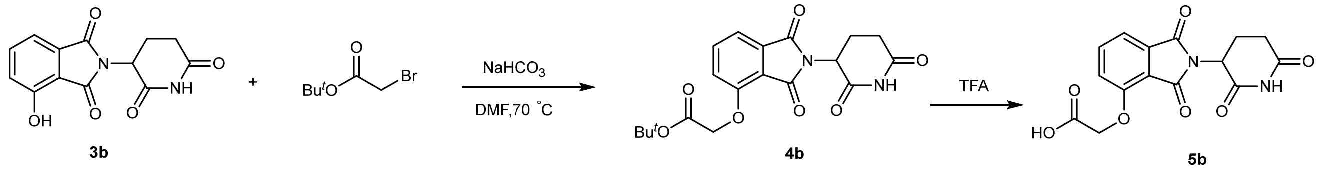


***tert*-Butyl 2-((2-(2,6-dioxopiperidin-3-yl)-1,3-dioxoisoindolin-4-yl)oxy)acetate (4b).** To a solution of compound **3b** (552 mg, 2 mmol) and *tert*-butyl 2-bromoacetate (467 mg, 2.4 mmol) in DMF (10 mL), add $NaHCO_3$ (252 mg, 3 mmol) and heated to 70°C for 4 hours, then the mixture was cooled to room temperature and concentrated under reduced pressure. The residue was purified by flash column chromatography (0-10% MeOH in DCM) to give compound **4b** as a white solid (582 mg, 75%). $^1$H NMR (400 MHz, $D_6$-DMSO) δ 8.31 (s, 1H), 7.81 (t, *J* = 7.9 Hz, 1H), 7.49 (d, *J* = 7.3 Hz, 1H), 7.38 (d, *J* = 8.6 Hz, 1H), 5.23 (dd, *J* = 13.1, 5.2 Hz, 1H), 4.97 (s, 2H), 3.09 (m, 1H), 2.84 (m, 1H), 2.65 (m, 1H), 2.11 (m, 1H), 1.43 (s, 9H), 1.40 (s, 9H).

**2-((2-(2,6-Dioxopiperidin-3-yl)-1,3-dioxoisoindolin-4-yl)oxy)acetic acid (5b)**. In a 100 mL flask was added **4b** (78 mg, 0.2 mmol) in TFA/DCM (10 mL, 1/1). The mixture was stirred for 30 min at room temperature. LC-MS showed **4b** converted into **5b** completely. Then remove the solvent *in vacuo* to give product **5b** (66 mg, 95%), which was used for for next step without further purification.

**General Procedure of RC PROTACs Synthesis (RC-1).**

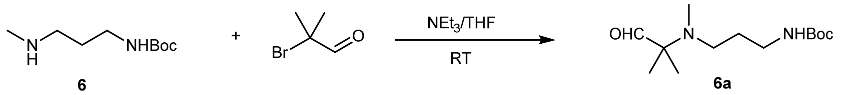


***tert*-Butyl (3-(methyl(2-methyl-1-oxopropan-2-yl)amino)propyl)carbamate (6a).** To a solution of compound **6** (658 mg, 3.5 mmol) in 20 mL THF, add $NEt_3$ (707 mg, 7 mmol) and 2-bromo-2-methylpropanal (1.05 g, 7 mmol). The solution was stirred for overnight. Then the reaction mixture was concentrated *in vacuo* and the residue was purified by flash column chromatography (5% MeOH in DCM) to give product **6a** (542 mg, 60%) as a yellow oil. $^1$H NMR (400 MHz, $CDCl_3$) δ 9.41 (s, 1H), 5.05 (s, 1H), 3.18 (q, *J* = 6.1 Hz, 2H), 2.35 (t, *J* = 6.6 Hz, 2H), 2.20 (s, 3H), 1.65 (m, 2H), 1.43 (s, 9H), 1.07 (s, 6H).

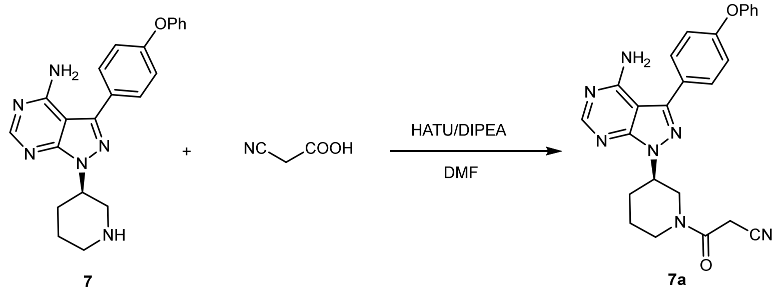


**(*R*)-3-(3-(4-Amino-3-(4-phenoxyphenyl)-1H-pyrazolo[3,4-d]pyrimidin-1-yl)piperidin-1-yl)-3-oxopropanenitrile (7a).** To a 100 mL flask was added compound **7** (1 g, 2.6 mmol), 2-cyanoacetic acid (330

mg, 3.9 mmol), HATU (1.48 g, 3.9 mmol) and DIPEA (671 mg, 5.2 mmol) in DMF (20 mL). Then the reaction mixture was concentrated *in vacuo* and the residue was purified by flash column chromatography (0~10% MeOH in Ethyl Acetate) to give product **7a** as a white solid (1.0 g, 90%). $^{1}$H NMR (400 MHz, *D₆*-DMSO) δ 8.26 (s, 1H), 7.67 (d, *J* = 8.0 Hz, 2H), 7.43 (m, 2H), 7.15 (m, 5H), 6.77 (s, 1H), 4.78 (m, 1H), 4.44 (m, 1H), 3.89 (m, 4H), 3.27 (m, 2H), 2.23 (m, 1H), 1.72 (m, 2H).

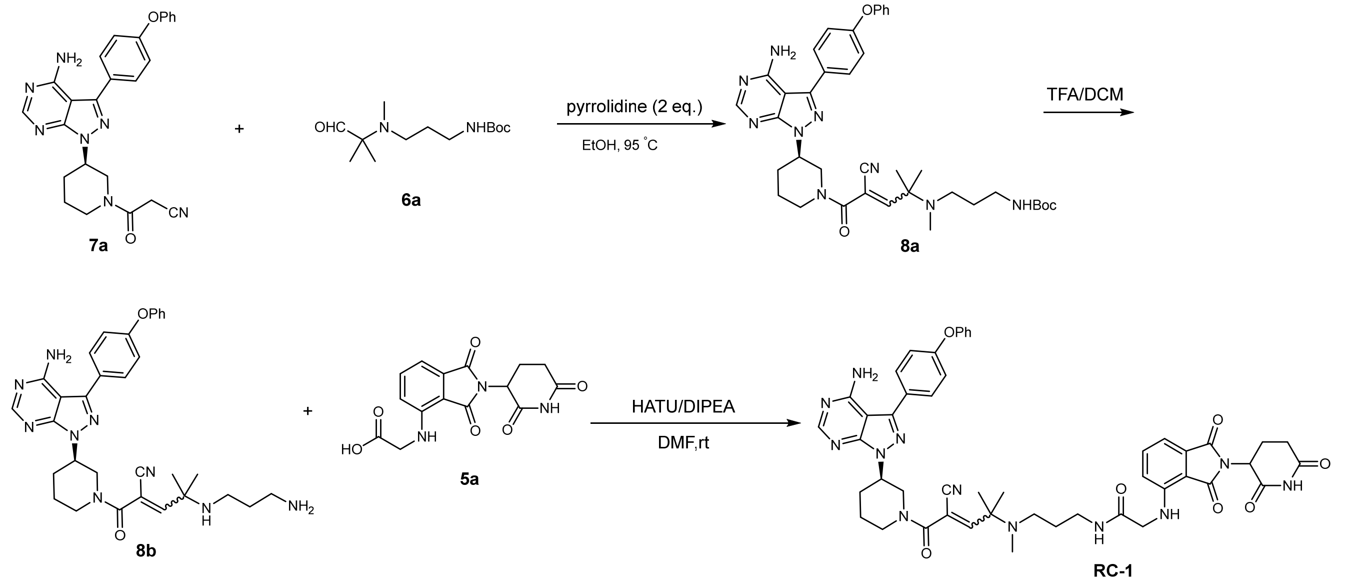


**tert-Butyl (*R*)-(3-((5-(3-(4-amino-3-(4-phenoxyphenyl)-1H-pyrazolo[3,4-d]pyr-imidin-1-yl)piperidin-1-yl)-4-cyano-2-methyl-5-oxopent-3-en-2-yl)(methyl)-amino)propyl)carbamate (8a).** To a round bottom flask was added compound **7a** (91 mg, 0.2 mmol), pyrrolidine (28 mg, 0.4 mmol), compound **6a** (103 mg, 0.4 mmol) in EtOH (3 mL) and heated to 95°C for 4 hours. TLC showed compound **8a** was generated as a major product. Then the reaction mixture was extracted with ethyl acetate, and the organic layer was washed with water, dried over $Na_2SO_4$, and filtered. The filtrate was concentrated *in vacuo* and the residue was purified by flash column chromatography (0~10% MeOH in Ethyl Acetate) to give product **8a** as a white solid (69 mg, 50%). $^{1}$H NMR (400 MHz, *D₆*-DMSO) δ 8.24 (s, 1H), 7.66 (d, *J* = 8.3 Hz, 2H), 7.43 (t, *J* = 7.9 Hz, 2H), 7.15 (m, 5H), 6.78 (br, 2H), 4.87 (s, 1H), 4.29 (s, 1H), 3.97 (s, 1H), 3.77 (s, 1H), 3.44 (m, 1H), 2.93 (m, 2H), 2.30 (s, 3H), 2.13 (m, 3H), 2.02 (m, 2H), 1.71 (m, 1H), 1.48 (m, 2H), 1.35 (s, 9H), 1.17 (d, *s*, 6H).

**N-(3-((5-((*R*)-3-(4-Amino-3-(4-phenoxyphenyl)-1H-pyrazolo[3,4-d]pyrimidin-1-yl)piperidin-1-yl)-4-cyano-2-methyl-5-oxopent-3-en-2-yl)(methyl)amino)propyl)-2-((2-(2,6-dioxopiperidin-3-yl)-1,3-dioxoisoindolin-4-yl)amino)acetamide (RC-1)**. In a 25 mL flask was added **8a** (35 mg, 0.05 mmol) in TFA/DCM (5 mL, 1/1). The mixture was stirred for 30 min at room temperature. LC-MS showed **8a** converted into **8b** completely. Then remove the solvent *in vacuo* to give product **8b** (28 mg, 95%), which was used for next step without further purification. To above **8b** was added compound **5a** (33 mg, 0.1 mol), HATU (38 mg, 0.1 mmol) and DIPEA (32 mg, 0.25 mmol) in DMF (3 mL). The mixture was stirred at room temperature for 30 min. Then the reaction mixture was concentrated *in vacuo* and the residue was purified by PrepHPLC with a reverse phase C18 column to afford the product as a yellow solid **RC-1** (18 mg, 40%). $^{1}$H NMR (400 MHz, *D₆*-DMSO) δ 8.24 (s, 1H), 7.92 (s, 1H), 7.67 (d, *J* = 8.4 Hz, 2H), 7.56 (t, *J* = 7.8 Hz, 1H), 7.43 (t, *J* = 7.8 Hz, 2H), 7.15 (m, 5H), 7.05 (d, *J* = 7.1

Hz, 1H), 6.89 (m, 2H), 6.76 (br, 2H), 5.04 (dd, *J* = 12.7, 5.3 Hz, 1H), 4.86 (m, 1H), 4.17 (m, 1H), 3.91 (d, *J* = 5.4 Hz, 2H), 3.80 (m, 1H), 3.57 (m, 1H), 3.10 (m, 2H), 2.87 (m, 1H), 2.59 (m, 3H), 2.29 (m, 3H), 2.16 (m, 1H), 2.10 (s, 3H), 2.02 (m, 2H), 1.72 (m, 1H), 1.55 (m, *J* = 6.1 Hz, 2H), 1.18 (m, 6H). $^{13}$C NMR (100 MHz, *D₆*-DMSO) δ 173.0, 170.2, 169.1, 168.7, 167.7, 163.0, 162.5, 158.7, 157.7, 156.8, 156.0, 154.6, 146.4, 143.8, 136.5, 132.6, 130.5, 128.4, 124.2, 119.44, 119.41, 119.4, 117.8, 115.1, 111.4, 110.5, 109.1, 98.0, 62.5, 59.9, 56.1, 52.3, 49.3, 49.1, 45.8, 37.4, 35.3, 34.7, 31.5, 28.5, 25.9, 23.1, 23.0, 22.7. HRMS (m/z): $[M+H]^+$ calcd. for $C_{48}H_{51}N_{12}O_7$, 907.4004; found: 907.4033

Follow **General Procedure of RC PROTACs Synthesis**, other RC PROTACs were synthesized.

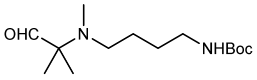


***tert*-Butyl (4-(methyl(2-methyl-1-oxopropan-2-yl)amino)butyl)carbamate (6b).** $^1$H NMR (400 MHz, $CDCl_3$) δ 9.44 (s, 1H), 4.98 (s, 1H), 3.11 (m, 2H), 2.28 (t, *J* = 6.0 Hz, 2H), 2.20 (s, 3H), 1.50 (m, 4H), 1.44 (s, 9H).

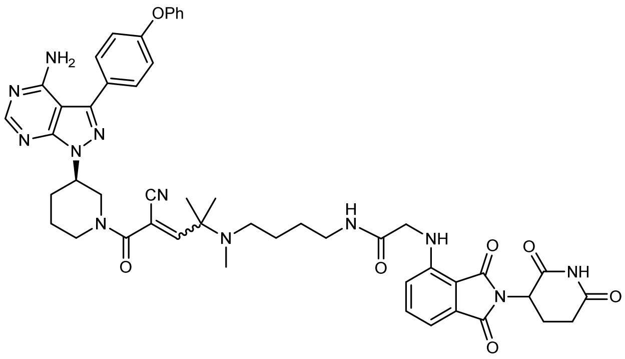


***N*-(4-((5-((*R*)-3-(4-Amino-3-(4-phenoxyphenyl)-1*H*-pyrazolo[3,4-*d*]pyrimidin-1-yl)piperidin-1-yl)-4-cyano-2-methyl-5-oxopent-3-en-2-yl)(methyl)amino)butyl)-2-((2-(2,6-dioxopiperidin-3-yl)-1,3-dioxoisoindolin-4-yl)amino)acetamide (RC-2).** $^1$H NMR (400 MHz, *D₆*-DMSO) δ 8.22 (s, 1H), 7.84 (s, 1H), 7.64 (d, *J* = 8.4 Hz, 2H), 7.54 (t, *J* = 7.8 Hz, 1H), 7.40 (t, *J* = 7.9 Hz, 2H), 7.13 (m, 5H), 7.04 (d, *J* = 6.8 Hz, 1H), 6.85 (m, 2H), 6.71 (br, 2H), 5.01 (dd, *J* = 12.5, 5.3 Hz, 1H), 4.83 (m, 1H), 4.14 (m, 1H), 3.88 (d, *J* = 5.5 Hz, 2H), 3.80 (m, 1H), 3.61 (m, 1H), 3.31 (m, 3H), 2.86 (m, 1H), 2.56 (m, 2H), 2.22 (m, 3H), 2.07 m, 3H), 2.013 (m, 2H), 1.68 (m, 1H), 1.35 (m, 4H), 1.18 (m, 6H). $^{13}$C NMR (100 MHz, *D₆*-DMSO) δ 172.9, 170.2, 169.1, 168.6, 167.7, 163.0, 162.5, 158.7, 157.8, 156.8, 156.1, 154.6, 146.4, 143.8, 136.6, 132.6, 130.5, 130.4, 128.4, 124.1, 119.4, 119.3, 117.8, 115.1, 111.4, 110.6, 109.2, 98.1, 59.9 , 52.3, 51.3, 49.3, 49.1, 45.9, 39.0, 35.4, 31.5, 29.4, 27.3, 26.0, 23.3, 22.7. HRMS (m/z): $[M+H]^+$ calcd. for $C_{49}H_{53}N_{12}O_7$, 921.4160; found: 921.4199.

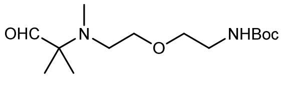

***tert*-Butyl (2-(2-(methyl(2-methyl-1-oxopropan-2-yl)amino)ethoxy)ethyl)car- bamate (6c).** $^{1}$H NMR (400 MHz, $CDCl_3$) δ 9.48 (s, 1H), 5.25 (s, 1H), 3.71 (q, *J* = 7.0 Hz, 2H), 3.52 (dd, *J* = 11.0, 5.4 Hz, 4H), 3.31 (m, 2H), 2.51 (t, *J* = 5.7 Hz, 2H), 2.29 (s, 3H), 1.44 (s, 9H), 1.10 (s, 6H).

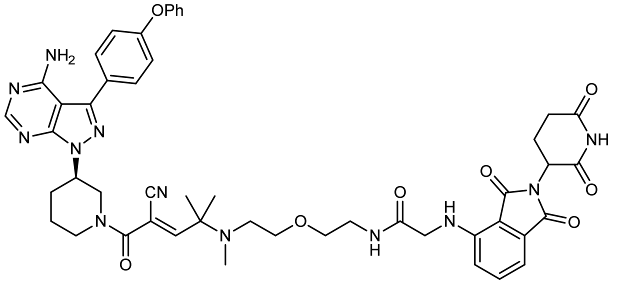


***N*-(2-(2-((5-((*R*)-3-(4-amino-3-(4-phenoxyphenyl)-1*H*-pyrazolo[3,4-*d*]pyrimidin-1-yl)piperidin-1-yl)-4-cyano-2-methyl-5-oxopent-3-en-2-yl)(methyl)amino)ethoxy)ethyl)-2-((2-(2,6-dioxopiperidin-3-yl)-1,3-dioxoisoindolin-4-yl)amino)acetamide (RC-3).** $^{1}$H NMR (400 MHz, *D$_6$*-DMSO) δ 8.24 (s, 1H), 7.93 (s, 1H), 7.67 (d, *J* = 8.5 Hz, 2H), 7.57 (t, *J* = 7.9 Hz, 1H), 7.43 (t, *J* = 7.9 Hz, 2H), 7.16 (m, 5H), 7.06 (d, *J* = 7.1 Hz, 1H), 6.78 (s, 2H), 6.72 (br, 1H), 5.04 (dd, *J* = 12.6, 5.4 Hz, 1H), 4.86 (m, 1H), 4.18 (m, 1H), 3.90 (d, *J* = 18.4 Hz, 2H), 3.63 (m, 1H), 3.40 (m, 6H), 3.25 (m, 4H), 2.87 (m, 1H), 2.65 (m, 3H), 2.31 (m, 1H), 2.18 (s, 3H), 2.04 (m, 2H), 1.72 (m, 1H), 1.20 (m, 6H). HRMS (m/z): $[M+H]^+$ calcd. for $C_{49}H_{53}N_{12}O_8$, 937.4109; found: 937.4137.

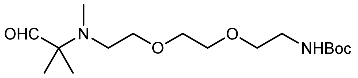


***tert*-Butyl (2-(2-(2-(methyl(2-methyl-1-oxopropan-2 yl)amino)ethoxy)ethoxy) ethyl)carbamate (6d).** $^{1}$H NMR (400 MHz, $CDCl_3$) δ 9.47 (s, 1H), 5.09 (s, 1H), 3.60 (m, 4H), 3.55 (m, 4H), 3.31 (dd, *J* = 5.0 Hz, 2H), 2.54 (t, *J* = 6.1 Hz, 2H), 2.29 (s, 3H), 1.44 (s, 9H), 1.09 (s, 6H).

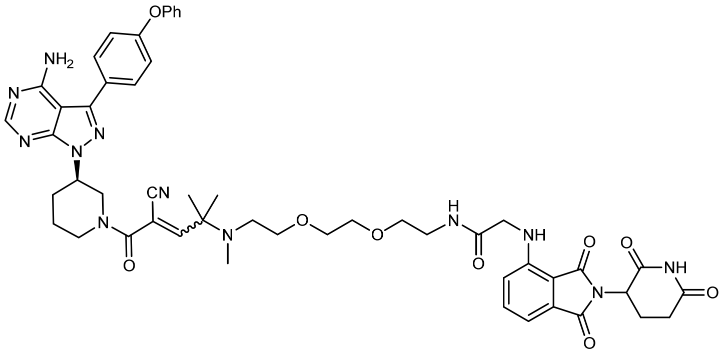


***N*-(2-(2-(2-((5-((*R*)-3-(4-amino-3-(4-phenoxyphenyl)-1*H*-pyrazolo[3,4-*d*]pyrimidin-1-yl)piperidin-1-yl)-4-cyano-2-methyl-5-oxopent-3-en-2-yl)(methyl)amino)ethoxy)ethoxy)ethyl)-2-((2-(2,6-dioxopiperidin-3-yl)-1,3-dioxoisoindolin-4-yl)amino)acetamide (RC-4).** $^{1}$H NMR (400 MHz, *D$_6$*-DMSO) δ 10.89 (s, 1H), 8.22 (s, 1H), 7.96 (s, 1H), 7.64 (d, *J* = 8.0 Hz, 2H), 7.55 (t, *J* = 7.7 Hz, 1H), 7.41 (t, *J* = 7.6 Hz, 2H), 7.13 (m, 5H), 7.03 (d, *J* = 7.0 Hz, 1H), 6.86 (m, 2H), 6.75 (s, 1H), 6.70 (br, 1H), 5.01 (dd, *J* = 12.6, 5.2 Hz, 1H), 4.83 (m, 1H), 4.13 (m, 1H), 3.91 (d, *J* = 5.2 Hz, 2H), 3.82 (m, 1H), 3.60 (m, 1H), 3.43 (m, 9H), 3.23 (m, 3H), 2.83 (m, 1H), 2.58 (m, 3H), 2.28 (m, 2H), 2.16 (s, 4H), 2.01 (m, 2H), 1.69 (s, 1H), 1.17 (m, 6H). $^{13}$C NMR (100 MHz, *D$_6$*-DMSO) δ 173.1, 170.3, 169.2, 169.0, 167.7, 163.0, 162.4, 158.7, 157.7, 156.8, 156.1, 154.6, 146.4, 143.8, 136.5, 132.6, 130.5,

128.4, 124.2, 119.4, 117.9, 115.0, 111.4, 110.5, 109.3, 98.1, 70.20, 70.17, 70.1, 69.4, 59.8, 56.0, 51.6, 49.2, 45.8, 39.2, 36.3, 31.5, 29.4, 23.3, 23.2, 22.7. HRMS (m/z): $[M+H]^+$ calcd. for $C_{51}H_{57}N_{12}O_9$, 981.4371; found: 981.4405.

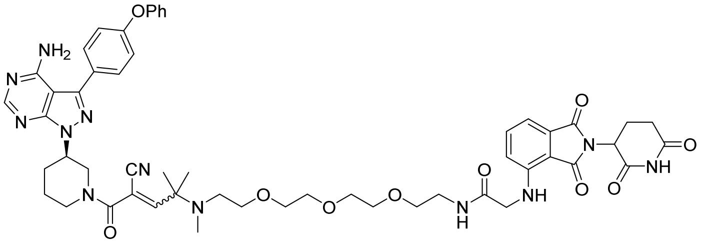

***N*-(16-((*R*)-3-(4-amino-3-(4-phenoxyphenyl)-1*H*-pyrazolo[3,4-*d*]pyrimidin-1-yl)piperidin-1-yl)-15-cyano-12,13,13-trimethyl-16-oxo-3,6,9-trioxa-12-azahexadec-14-en-1-yl)-2-((2-(2,6-dioxopiperidin-3-yl)-1,3-dioxoisoindolin-4-yl)amino)acetamide (RC-5)** $^1$H NMR (400 MHz, *D$_6$*-DMSO) δ 10.92 (s, 1H), 8.24 (s, *J* = 8.0 Hz, 1H), 7.99 (s, 1H), 7.67 (d, *J* = 8.5 Hz, 2H), 7.58 (t, *J* = 8.o Hz, 1H), 7.43 (t, *J* = 7.9 Hz, 2H), 7.16 (m, 5H), 7.06 (d, *J* = 7.1 Hz, 1H), 6.88 (m, 2H), 6.78 (s, 1H), 6.73 (br, 1H), 5.04 (dd, *J* = 12.7, 5.4 Hz, 1H), 4.86 (m, 1H), 4.17 (m, 1H), 3.91 (t, *J* = 4.0 Hz, 2H), 3.85 (m, 1H), 3.63 (m, 1H), 3.45 (m, 13H), 3.28 (m, 4H), 2.89 (m, 1H), 2.57 (m, 2H), 2.30 (m, 1H), 2.18 (s, 4H), 2.04 (m, 2H), 1.72 (m, 1H), 1.21 (m, 6H). HRMS (m/z): $[M+H]^+$ calcd. for $C_{53}H_{61}N_{12}O_{10}$, 1025.4634; found: 1025.4668.

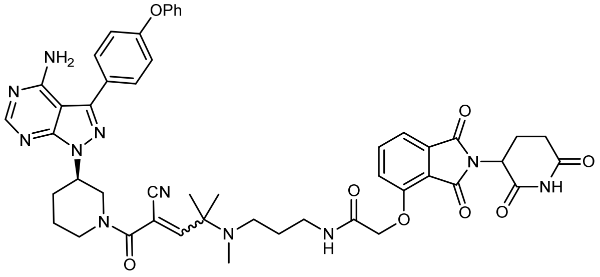

***N*-(3-((5-((*R*)-3-(4-amino-3-(4-phenoxyphenyl)-1*H*-pyrazolo[3,4-*d*]pyrimidin-1-yl)piperidin-1-yl)-4-cyano-2-methyl-5-oxopent-3-en-2-yl)(methyl)amino)propyl)-2-((2-(2,6-dioxopiperidin-3-yl)-1,3-dioxoisoindolin-4-yl)oxy)acetamide (RC-9).** $^1$H NMR (400 MHz, *D$_6$*-DMSO) δ 10.93 (s, 1H), 8.21 (s, 1H), 7.76 (t, *J* = 7.7 Hz, 2H), 7.64 (d, *J* = 8.0 Hz, 2H), 7.41 (m, 4H), 7.13 (m, 5H), 6.68 (br, 3H), 5.06 (dd, *J* = 12.6, 5.3 Hz, 1H), 4.82 (m, 1H), 4.71 (s, 2H), 4.13 (m, 1H), 3.75 (m, 2H), 2.83 (m, 1H), 2.57 (m, 3H), 2.33 (m, 2H), 2.26 (m, *J* = 9.6 Hz, 1H), 2.15 (m, 1H), 2.09 (s, 3H), 2.02 (m, 2H), 1.67 (m, 1H), 1.57 (m, 2H), 1.20 (s, 6H). $^{13}$C NMR (100 MHz, *D$_6$*-DMSO) δ 172.9, 170.0, 167.1, 167.0, 165.9, 163.0, 162.5, 158.7, 157.7, 156.8, 156.0, 155.5, 154.6, 143.8, 137.3, 133.6, 130.5, 128.4, 124.2, 121.1, 119.4, 117.5, 116.6, 115.1, 109.1, 98.1, 68.5, 60.0, 52.4, 49.5, 49.4, 49.2, 41.1, 37.2, 35.4, 31.4, 29.3, 28.4, 25.9, 23.2, 23.0, 22.5. HRMS (m/z): $[M+H]^+$ calcd. for $C_{48}H_{50}N_{11}O_8$, 908.3844; found: 908.3870.

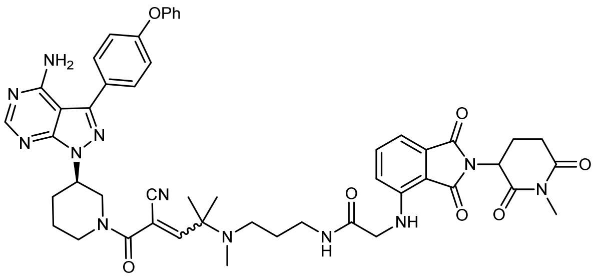

***N*-(3-((5-((*R*)-3-(4-amino-3-(4-phenoxyphenyl)-1*H*-pyrazolo[3,4-*d*]pyrimidin-1-yl)piperidin-1-yl)-4-cyano-2-methyl-5-oxopent-3-en-2-yl)(methyl)amino)propyl)-2-((2-(1-methyl-2,6-dioxopiperidin-3-yl)-1,3-dioxoisoindolin-4-yl)amino)acetamide (RC-1-Me).** $^{1}$H NMR (400 MHz, *D$_6$*-DMSO) δ 8.24 (s, 1H), 7.88 (s, 1H), 7.67 (d, *J* = 7.1 Hz, 2H), 7.56 (t, *J* = 7.2 Hz, 1H), 7.43 (t, *J* = 7.1 Hz, 2H), 7.15 (m, 5H), 7.06 (d, *J* = 7.0 Hz, 1H), 6.88 (m, 2H), 6.76 (s, 1H), 6.71 (br, 2H), 5.11 (dd, *J* = 12.6, 3.8 Hz, 1H), 4.86 (m, 1H), 4.17 (m, 1H), 3.91 (d, *J* = 4.2 Hz, 3H), 3.63 (m, 1H), 3.32 (m, 1H), 3.11 (m, 2H), 3.04 (s, 3H), 2.92 (m, 1H), 2.77 (m, 1H), 2.57 (m, 1H), 2.31 (m, 3H), 2.19 (m, 1H), 2.11 (s, 3H), 2.04 (m, 2H), 1.71 (m, 1H), 1.57 (m, 2H), 1.20 (s, 6H). $^{13}$C NMR (100 MHz, *D$_6$*-DMSO) δ 172.1, 170.0, 169.1, 168.7, 167.7, 163.0, 162.5, 158.7, 157.8, 157.7, 156.8, 156.1, 156.0, 154.6, 146.4, 143.8, 136.6, 136.5, 132.6, 130.5, 130.4, 128.4, 124.2, 119.4, 119.3, 117.9, 115.1, 111.5, 111.4, 110.5, 109.2, 98.1, 59.9, 56.2, 52.4, 52.3, 49.8, 49.7, 49.2, 47.8, 45.9, 37.5, 35.4, 34.8, 31.6, 29.4, 28.5, 26.9, 23.6, 23.1, 21.9. HRMS (m/z): [M+H]$^+$ calcd. for $C_{49}H_{53}N_{12}O_7$, 921.4160; found: 921.4192.

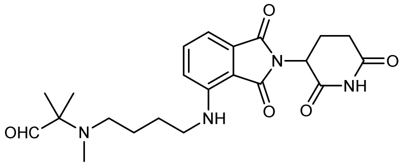

**2-((4-((2-(2,6-dioxopiperidin-3-yl)-1,3-dioxoisoindolin-4-yl)amino)butyl)(methyl)amino)-2-methylpropanal (6e).** $^{1}$H NMR (400 MHz, $CDCl_3$) δ 9.43 (s, 1H), 7.50 (t, *J* = 7.4 Hz, 1H), 7.09 (d, *J* = 7.3 Hz, 1H), 6.88 (d, *J* = 8.3 Hz, 1H), 6.24 (s, 1H), 4.91 (m, 1H), 3.29 (q, *J* = 6.6 Hz, 2H), 2.81 (m, 3H), 2.32 (t, *J* = 6.9 Hz, 2H), 2.22 (s, 3H), 2.11 (m, 1H), 1.69 (m, 2H), 1.60 (m, 2H), 1.08 (s, 6H).

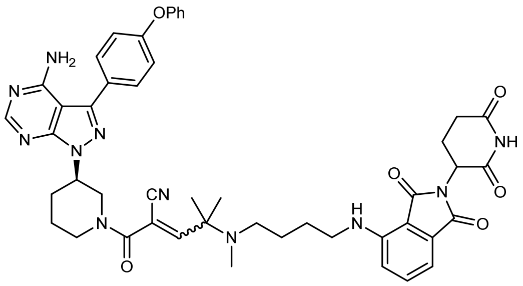

**2-((*R*)-3-(4-amino-3-(4-phenoxyphenyl)-1*H*-pyrazolo[3,4-d]pyrimidin-1-yl)piperidine-1-carbonyl)-4-((4-((2-(2,6-dioxopiperidin-3-yl)-1,3-dioxoisoindolin-4-yl)amino)butyl)(methyl)amino)-4-methylpent-2-enenitrile (RC-7).** $^{1}$H NMR (400 MHz, *D$_6$*-DMSO) δ 11.09 (s, 1H), 8.24 (s, 1H), 7.65 (m, 2H), 7.54 (m, 1H), 7.43 (t, *J* = 6.9 Hz, 2H), 7.18 (m, 5H), 7.00 (d, *J* = 6.4 Hz, 1H), 6.75 (m, 1H), 6.47 (br, 1H), 5.03 (m, 1H), 4.86 (s, 1H), 4.70 (m, 1H), 4.24 (m, 1H), 3.96 (m, 1H), 3.68 (m, 2H), 3.17 (m, 2H), 2.87 (m, 2H), 2.28 (m, 3H), 2.12 (s, 3H), 2.01 (m, 3H), 1.66 (m, 1H), 1.48 (m, 4H), 1.18 (s, 6H). $^{13}$C NMR (100 MHz, *D$_6$*-DMSO) δ 173.2, 170.5, 169.3, 167.7, 162.9, 162.7, 158.6, 157.6, 156.7, 156.1, 154.4, 146.8, 143.8, 136.7, 132.6, 130.6, 130.5, 128.3, 124.2, 119.41, 119.37,

117.6, 115.2, 110.8, 109.4, 108.9, 97.8, 59.9, 59.8, 52.1, 51.3, 49.0, 48.9, 42.2, 35.2, 31.4, 29.4, 26.9, 25.8, 23.0, 22.6. HRMS (m/z): $[M+H]^+$ calcd. for $C_{47}H_{50}N_{11}O_6$, 864.3946; found: 864.3976.

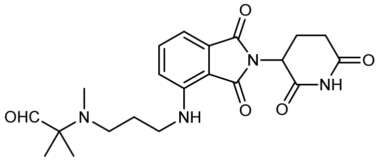


**2-((3-((2-(2,6-dioxopiperidin-3-yl)-1,3-dioxoisoindolin-4-yl)amino)propyl)(methyl)amino)-2-methylpropanal (6f).** $^1$H NMR (400 MHz, $CDCl_3$) δ 9.46 (s, 1H), 7.49 (t, *J* = 7.9 Hz, 1H), 7.08 (d, *J* = 7.2 Hz, 1H), 6.88 (d, *J* = 8.8 Hz, 1H), 6.76 (s, 1H), 4.92 (m, 1H), 3.36 (q, *J* = 5.6 Hz, 2H), 2.81 (m, 3H), 2.46 (t, *J* = 5.8 Hz, 2H), 2.27 (s, 3H), 2.12 (m, 1H), 1.82 (m, 2H), 1.10 (s, 6H).

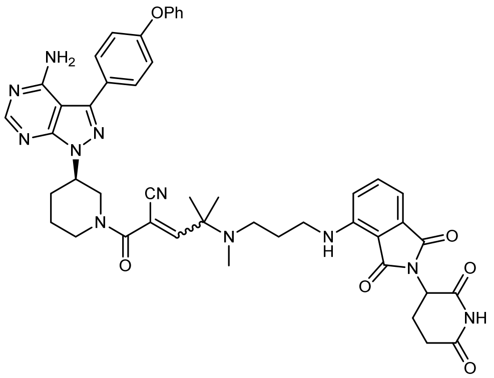


**2-((*R*)-3-(4-amino-3-(4-phenoxyphenyl)-1*H*-pyrazolo[3,4-d]pyrimidin-1-yl)piperidine-1-carbonyl)-4-((3-((2-(2,6-dioxopiperidin-3-yl)-1,3-dioxoisoindolin-4-yl)amino)propyl)(methyl)amino)-4-methylpent-2-enenitrile (RC-8).** $^1$H NMR (400 MHz, $D_6$-DMSO) δ 11.04 (s, 1H), 8.22 (s, 1H), 7.62 (d, *J* = 6.1 Hz, 2H), 7.51 (s, 1H), 7.40 (t, *J* = 7.6 Hz, 2H), 7.13 (m, 5H), 6.97 (d, *J* = 6.3 Hz, 1H), 6.71 (m, 2H), 5.01 (m, 1H), 4.82 (m, 1H), 4.52 (m, 1H), 4.16 (m, 1H), 4.00 (m, 1H), 3.73 (m, 2H), 3.18 (m, 2H), 2.83 (m, 2H), 2.29(m, 3H), 2.12 (s, 3H), 1.97 (m, *J* = 5.8 Hz, 3H), 1.66 (m, 3H), 1.16 (s, 6H). $^{13}$C NMR (100 MHz, $D_6$-DMSO) δ 173.2, 170.5, 169.3, 167.7, 162.9, 162.5, 158.6, 157.6, 156.7, 156.1, 154.4, 146.8, 143.8, 136.7, 132.6, 130.6, 130.5, 128.3, 124.2, 119.42, 119.38, 117.5, 115.3, 110.8, 109.5, 108.9, 97.8, 60.8, 60.1, 52.1, 49.1, 49.0, 40.9, 40.8, 35.4, 31.4, 29.5, 27.3, 23.0, 22.6, 17.5. HRMS (m/z): $[M+H]^+$ calcd. for $C_{46}H_{48}N_{11}O_6$, 850.3789; found: 850.3811.

**General procedure of IRC and RNC PROTACs synthesis (IRC-1 and RNC-1).**

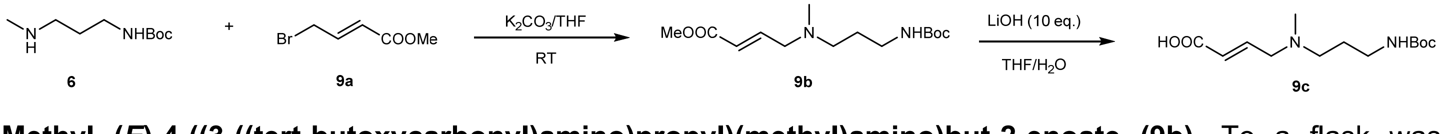


**Methyl (*E*)-4-((3-((tert-butoxycarbonyl)amino)propyl)(methyl)amino)but-2-enoate (9b).** To a flask was added compound **6** (376 mg, 2 mmol), $K_2CO_3$ (552 mg, 4 mmol) in 20 mL THF. The mixture was stirred for 30 min at room temperature. Then compound **9a** (708 mg, 4 mmol) in 2 mL THF was added dropwise with stirring. The mixture was stirred at room temperature for overnight. The solvent was concentrated *in vacuo* and the residue was purified by flash column chromatography (5% MeOH in DCM) to give product **9b** (354 mg, 62%) as a yellow oil. $^{1}$H NMR (400 MHz, $CDCl_3$) δ 6.84 (dt, *J* = 15.7, 6.1 Hz, 1H), 5.88 (dt, *J* = 15.7, 1.4 Hz, 1H), 5.25 (s, 1H), 3.64 (s, 3H), 3.05 (m, 4H), 2.32 (t, *J* = 6.7 Hz, 2H), 2.12 (s, 3H), 1.56 (m, 2H), 1.34 (s, 9H).

**(*E*)-4-((3-((tert-butoxycarbonyl)amino)propyl)(methyl)amino)but-2-enoic acid (9c)** Compound **9b** (143 mg, 0.5 mmol) and LiOH (120 mg, 5 mmol) were dissolved in THF (5 mL) and water (5 mL). The mixture was stirred at 40°C for 6 h. TLC showed **9b** completely disappeared. The mixture was cooled to 0°C and the pH was slowly adjusted to 4–5 with 1N HCl. The solvent was concentrated *in vacuo* and the residue was dissolve in DMF to give a solution **9b** (0.2 mM in DMF), which was used for next step without further purification.

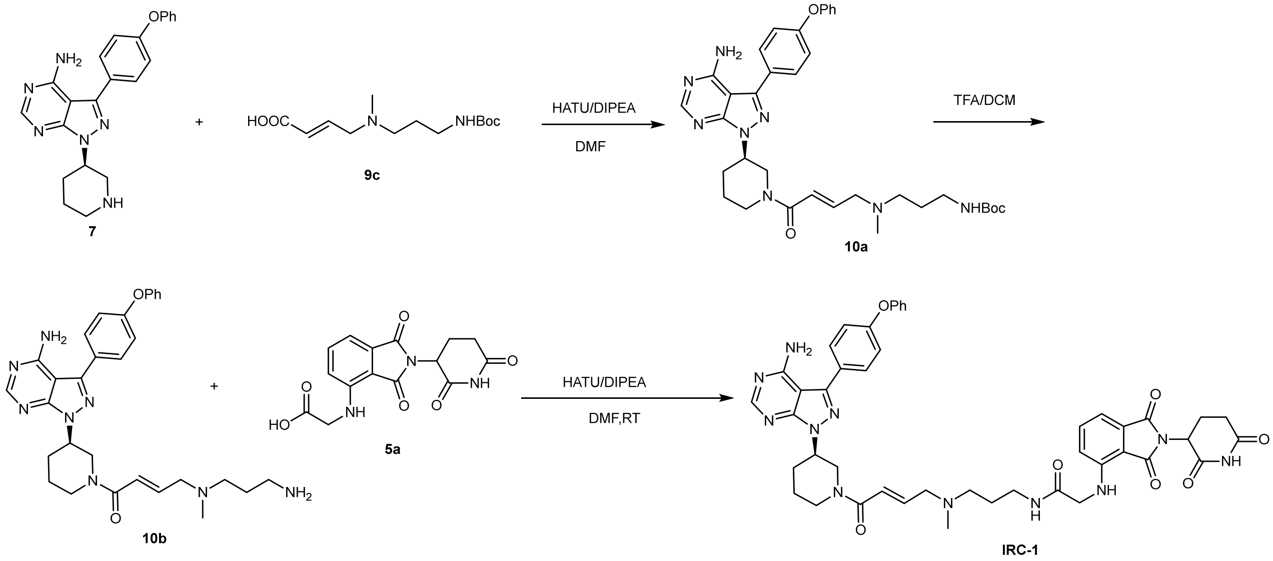


***tert*-Butyl (*R,E*)-(3-((4-(3-(4-amino-3-(4-phenoxyphenyl)-1H-pyrazolo[3,4-d]py-rimidi- n-1-yl)piperidin-1-yl)-4-oxobut-2-en-1-yl)(methyl)amino)propyl)- carbamate (10a).** To a flask was added compound **7** (77 mg, 0.2 mmol), solution **9c** (1.5 mL, 0.3 mmol), HATU (114 mg, 0.3 mmol) and DIPEA (129 mg, 1 mmol) in DMF(2 mL). The mixture was stirred at room temperature for 30 min. Then the reaction mixture was extracted with ethyl acetate, and the organic layer was washed with water, dried over $Na_2SO_4$, and filtered. The filtrate was concentrated *in vacuo* and the residue was purified by flash column chromatography (0~10% MeOH in Ethyl Acetate) to give product **10a** as a white solid (70 mg, 55%).

***N*-(3-(((*E*)-4-((*R*)-3-(4-Amino-3-(4-phenoxyphenyl)-1H-pyrazolo[3,4-d]pyrimi-din-1-yl)piperidin-1-yl)-4-ox-obut-2-en-1-yl)(methyl)amino)propyl)-2-((2-(2,6-dioxopiperidin-3-yl)-1,3-dioxoisoindolin-4-yl)amino)acet-amide (IRC-1).** In a 25 mL flask was added **10a** (54 mg, 0.1 mmol) in TFA/DCM (5 mL, 1/1). The mixture was stirred for 30 min at room temperature. LC-MS showed **10a** converted into **10b** completely. Then remove the solvent *in vacuo* to give product **10b** (52 mg, 95%), which was used for next step without further purification. To above **10b** was added compound **5a** (66 mg, 0.2 mol), HATU (78 mg, 0.2 mmol) and DIPEA (64 mg, 0.5 mmol) in DMF (3 mL). The mixture was stirred at room temperature for 30 min. Then the reaction mixture was concentrated *in vacuo* and the residue was purified by PrepHPLC with a reverse phase C18 column to afford the product as a yellow solid I**RC-1** (30 mg, 35%). $^{1}$H NMR (400 MHz, $D_6$-DMSO) δ 11.11 (s, 1H), 8.25 (s, 1H), 8.10 (d, *J* = 19.6 Hz, 1H), 7.65 (m, 2H), 7.57 (t, *J* = 7.8 Hz, 1H), 7.43 (t, *J* = 7.7 Hz, 2H), 7.16 (m, 5H), 7.06 (d, *J*

= 7.0 Hz, 1H), 6.96 (m, 1H), 6.85 (t, *J* = 8.0 Hz 1H), 6.63 (s, 1H), 6.46 (br, 1H), 5.07 (dd, *J* = 12.8, 5.2 Hz, 1H), 4.69 (m, 1H), 4.53 (d, *J* = 11.3 Hz, 1H), 4.10 (m, 1H), 3.91 (s, 2H), 3.76 (m, 1H), 3.15 (m, 5H), 2.88 (m, 2H), 2.61 (m, 2H), 2.26 (m, 3H), 2.11 (s, 3H), 2.01 (m, 2H), 1.94 (m, 1H), 1.56 (m, 3H). $^{13}C$ NMR (100 MHz, *D6*-DMSO) δ 173.0, 170.2, 169.1, 168.7, 167.7, 165.1, 158.6, 157.7, 156.8, 156.0, 154.5, 146.4, 143.6, 142.2, 136.6, 132.6, 130.49, 130.46, 128.5, 124.2, 122.9, 119.42, 119.39, 117.8, 111.4, 110.5, 98.0, 58.5, 54.8, 53.0, 49.18, 49.14, 45.8, 42.2, 37.6, 31.5, 29.8, 27.2, 25.9, 22.7. HRMS (m/z): $[M+H]^+$ calcd. for $C_{45}H_{48}N_{11}O_7$, 854.3738; found: 854.3772.

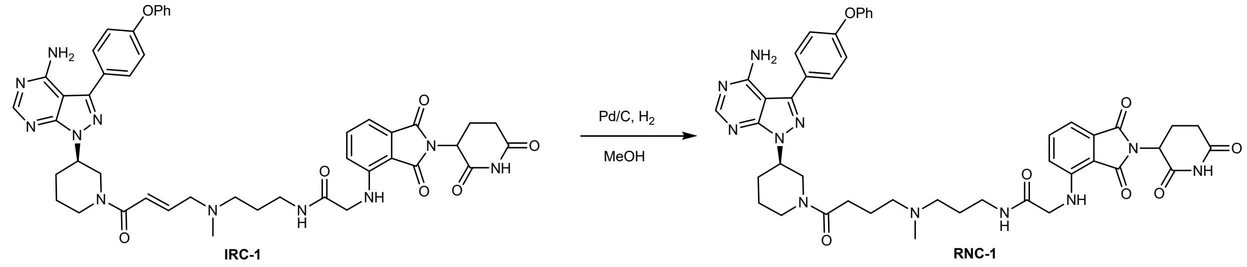


***N*-(3-((4-((*R*)-3-(4-amino-3-(4-phenoxyphenyl)-1*H*-pyrazolo[3,4-d]pyrimidin-1-yl)pi-peridin-1-yl)-4-oxo-butyl)(methyl)amino)propyl)-2-((2-(2,6-dioxopiperidin-3-yl)-1,3-dioxoisoindolin-4-yl)amino)acetamide (RNC-1).** To a flask was added **IRC-1** (17 mg, 0.02 mmol) and Pd/C (1.7 mg, 10%) in MeOH (2 mL). The mixture was stirred under 1atm $H_2$ at room temperature overnigh. LC-MS showed **IRC-1** converted into **RNC-1** completely. Then the reaction mixture was filtered, and the filtrate was concentrated *in vacuo* and the residue was purified by PrepHPLC with a reverse phase C18 column to afford the product as a yellow solid **RNC-1** (9.4 mg, 55%). $^1H$ NMR (400 MHz, *D6*-DMSO) δ 11.08 (s, 1H), 8.30 (s, 1H), 8.21 (s, 1H), 8.10 (m, 1H), 7.63 (d, *J* = 7.9 Hz, 2H), 7.53 (t, *J* = 7.4 Hz, 1H), 7.40 (t, *J* = 7.4 Hz, 2H), 7.20 – 7.06 (m, 5H), 7.03 (d, *J* = 6.3 Hz, 1H), 6.93 (m, 1H), 6.82 (t, *J* = 7.2 Hz, 1H), 5.04 (d, *J* = 7.8 Hz, 1H), 4.65 (m, 1H), 4.49 (d, *J* = 12.5 Hz, 1H), 4.16 (d, *J* = 13.2 Hz, 1H), 3.99 (d, *J* = 12.5 Hz, 1H), 3.87 (d, *J* = 9.3 Hz, 2H), 3.07 (m, 4H), 2.84 (m, 2H), 2.57 (s, 2H), 2.23 (m, 6H), 2.06 (s, 3H), 2.01 (s, 2H), 1.86 (s, 1H), 1.55 (m, 5H). $^{13}C$ NMR (100 MHz, *D6*-DMSO) δ 173.2, 171.1 170.5, 169.1, 168.7, 167.7, 158.6, 157.5, 156.7, 156.1, 154.4, 146.2, 143.6, 136.6, 132.5, 130.6, 130.50, 130.46, 128.4, 124.2, 119.4, 117.8, 111.4, 110.3, 97.8, 56.9, 55.0, 52.5, 49.7, 49.0, 45.7, 42.1, 40.9, 37.6, 31.41, 30.5, 29.7, 27.0, 25.1, 22.8, 22.6. HRMS (m/z): $[M+H]^+$ calcd. for $C_{45}H_{50}N_{11}O_7$, 856.3895; found: 856.3920.

Follow **General Procedure of IRC and RNC PROTACs Synthesis**, other IRC and RNC RPTOACs were synthesized.

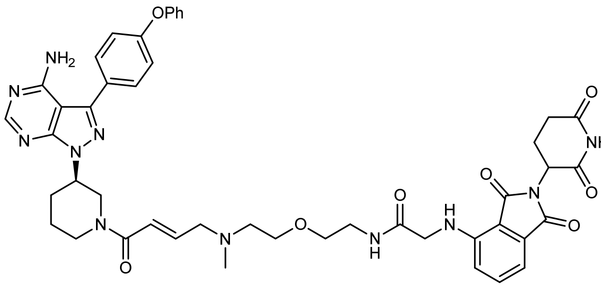

***N*-(2-(2-(((*E*)-4-((*R*)-3-(4-amino-3-(4-phenoxyphenyl)-1*H*-pyrazolo[3,4-d]pyrimidin-1-yl)piperidin-1-yl)-4-oxobut-2-en-1-yl)(methyl)amino)ethoxy)ethyl)-2-((2-(2,6-dioxopiperidin-3-yl)-1,3-dioxoisoindolin-4-yl)amino)acetamide (IRC-3)** $^{1}$H NMR (400 MHz, $D_6$-DMSO) δ 8.25 (s, 1H), 8.01 (s, 1H), 7.66 (d, *J* = 8.1 Hz, 2H), 7.57 (t, *J* = 7.7 Hz, 1H), 7.43 (t, *J* = 7.7 Hz, 2H), 7.14 (m, 5H), 7.06 (d, *J* = 7.0 Hz, 1H), 6.88 (d, *J* = 8.7 Hz, 2H), 6.73 (s, 1H), 6.56 (s, 2H), 5.11 – 4.95 (m, 1H), 4.71 (s, 1H), 4.04 (s, 1H), 3.93 (d, *J* = 5.0 Hz, 2H), 3.43 (d, *J* = 16.6 Hz, 5H), 3.18 – 3.02 (m, 6H), 2.87 (dd, *J* = 21.8, 9.3 Hz, 1H), 2.60 (d, *J* = 18.5 Hz, 2H), 2.35 – 2.20 (m, 2H), 2.16 (s, 4H), 2.04 (d, *J* = 11.5 Hz, 1H), 1.94 (s, 1H), 1.59 (d, *J* = 11.3 Hz, 1H). $^{13}$C NMR (100 MHz, $D_6$-DMSO) δ 172.9, 170.2, 169.1, 169.0, 167.7, 165.1, 158.7, 157.7, 156.8, 156.04, 156.00, 154.6, 146.4, 143.6, 142.2, 136.6, 136.5, 132.6, 130.5, 130.4, 128.5, 124.2, 122.9, 119.44, 119.35, 117.9, 111.5, 111.4, 110.6, 98.1, 69.3, 69.0, 58.8, 56.5, 53.0, 49.3, 49.1, 45.8, 42.9, 42.8, 41.2, 41.1, 39.2, 31.5, 29.8, 22.7. HRMS (m/z): $[M+H]^+$ calcd. for $C_{46}H_{50}N_{11}O_8$, 884.3833; found: 884.3868.

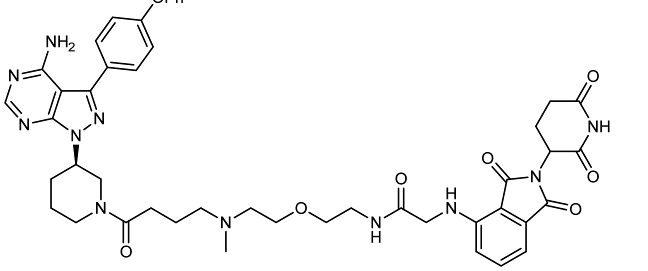

***N*-(2-(2-((4-((*R*)-3-(4-amino-3-(4-phenoxyphenyl)-1*H*-pyrazolo[3,4-d]pyrimidin-1-yl)piperidin-1-yl)-4-oxobutyl)(methyl)amino)ethoxy)ethyl)-2-((2-(2,6-dioxopiperidin-3-yl)-1,3-dioxoisoindolin-4-yl)amino)acetamide (RNC-3).** $^{1}$H NMR (400 MHz, $D_6$-DMSO) δ 8.22 (s, 1H), 7.94 (s, 1H), 7.64 (d, *J* = 8.4 Hz, 2H), 7.54 (t, *J* = 7.8 Hz, 1H), 7.40 (t, *J* = 7.8 Hz, 2H), 7.13 (m, 5H), 7.03 (d, *J* = 7.1 Hz, 1H), 6.86 (m, 2H), 6.68 (br, 2H), 5.01 (dd, *J* = 12.7, 5.4 Hz, 1H), 4.65 (m, 1H), 3.91 (d, *J* = 5.3 Hz, 2H), 3.39 (m, 5H), 2.86 (m, 1H), 2.58 (d, *J* = 19.8 Hz, 2H), 2.43 (m, 3H), 2.26 (m, 4H), 2.12 (s, 4H), 2.01 (m, 1H), 1.88 (m, 1H), 1.60 (m, 3H). $^{13}$C NMR (100 MHz, $D_6$-DMSO) δ 172.9, 171.3, 170.2, 169.1 ,169.0, 167.7, 158.7, 157.7, 156.8, 156.1, 156.0, 154.6, 146.4, 143.6, 136.6, 136.5, 132.6, 130.6, 130.4, 128.5, 124.2, 119.5, 119.4, 117.9, 111.5, 111.4, 110.6, 98.1, 69.3, 69.1, 57.3, 56.8, 49.3, 49.2, 45.9, 42.8, 41.2, 41.1, 39.3, 31.4, 30.5, 29.8, 23.1, 22.7. HRMS (m/z): $[M+H]^+$ calcd. for $C_{46}H_{52}N_{11}O_8$, 886.4000; found: 886.4027.

**Procedure of RNC-CN-DiMe and IRC-DiMe synthesis**

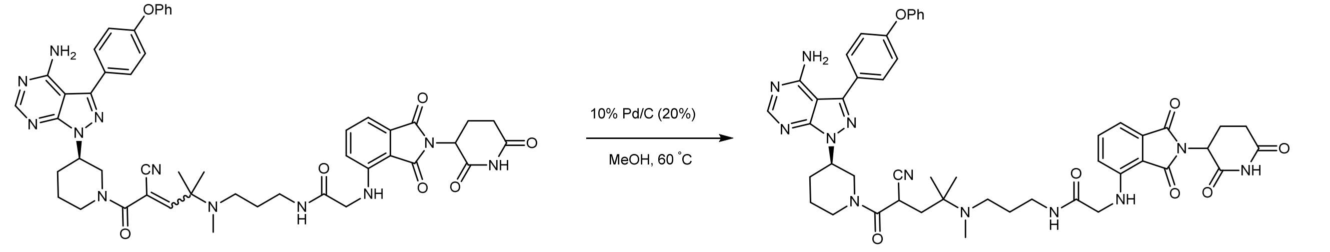


RC-1 RNC-1-CN-DiMe

***N*-(3-((5-((*R*)-3-(4-amino-3-(4-phenoxyphenyl)-1*H*-pyrazolo[3,4-*d*]pyrimidin-1-yl)piperidin-1-yl)-4-cyano-2-methyl-5-oxopentan-2-yl)(methyl)amino)propyl)-2-((2-(2,6-dioxopiperidin-3-yl)-1,3-dioxoisoindolin-4-yl)amino)acetamide (RNC-CN-DiMe)**. To a flask was added **RC-1** (18 mg, 0.02 mmol) and Pd/C (3.6 mg, 20%)

in MeOH (2 mL). The mixture was stirred under 1 atm $H_2$ at 60 ˚C overnigh. LC-MS showed **RC-1** converted into **RNC-CN-DiMe** completely. Then the reaction mixture was filtered, and the filtrate was concentrated *in vacuo* and the residue was purified by PrepHPLC with a reverse phase C18 column to afford the product as a yellow solid **RNC-CN-Dime** (8.5 mg, 47%). $^1$H NMR (400 MHz, *D$_6$*-DMSO) δ 11.09 (s, 1H), 8.22 (m, 2H), 8.00 (m, 1H), 7.62 (d, *J* = 8.3 Hz, 2H), 7.52 (t, *J* = 7.7 Hz, 1H), 7.39 (d, *J* = 7.8 Hz, 2H), 7.20 – 7.06 (m, 5H), 7.02 (d, *J* = 7.1 Hz, 1H), 6.93 (m, 1H), 6.80 (m, 1H), 5.03 (dd, *J* = 12.9, 5.0 Hz, 1H), 4.67 (m, 1H), 4.34 (m, 1H), 4.18 (m, 2H), 3.89 (m, 6H), 3.32 (m, 1H), 3.02 (m, 4H), 2.54 (m, 2H), 2.24 (m, 2H), 2.11 (m, 2H), 2.05 – 1.85 (m, 6H), 1.48 (m, 2H), 1.08 – 0.80 (m, 6H). $^{13}$C NMR (100 MHz, *D$_6$*-DMSO) δ 173.2, 170.5, 169.1, 168.7, 168.6, 167.7, 164.9, 164.6, 158.6, 157.5, 156.7, 156.1, 154.4, 146.2, 143.8, 136.6, 132.5, 130.6, 130.5, 128.3, 124.2, 120.0, 119.7, 119.4, 117.8, 111.4, 110.3, 97.8, 56.2, 52.2, 49.0, 47.5, 46.7, 46.2, 45.7, 40.8, 37.2, 34.7, 31.4, 29.9, 29.4, 28.5, 24.5, 23.6, 23.0, 22.6. HRMS (m/z): $[M+H]^+$ calcd. for $C_{48}H_{53}N_{12}O_7$, 909.4160; found: 909.4178.

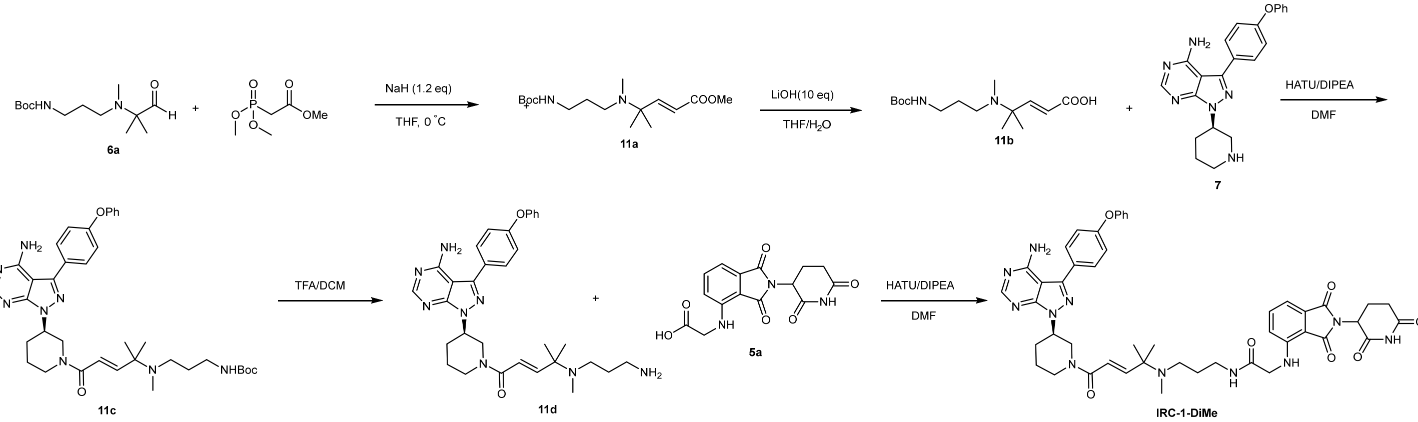


**Methyl (*E*)-4-((3-((tert-butoxycarbonyl)amino)propyl)(methyl)amino)-4-methylpent-2-enoate (11a).** To a flask was added NaH (18 mg, 0.43 mmol) in 5 mL THF at 0 ˚C. Then trimethyl phosphonoacetate (98 mg, 0.54 mmol) in 2 mL THF was added dropwise with stirring. The mixture was stirred at 0 ˚C for 15 min. Then **6a** (93 mg, 0.36) in 2 mL THF was added dropwise and the mixture was stirred for overnight at room temperature. Remove the solvent *in vacuo* and the residue was purified by flash column chromatography (5% MeOH in DCM) to give product **11a** (85 mg, 75%) as a light yellow oil. LC/MS: m/z 315 $[M+H]^+$.

**(*E*)-4-((3-((tert-butoxycarbonyl)amino)propyl)(methyl)amino)-4-methylpent-2-enoic acid(11b).** Compound **11a** (85 mg, 0.27 mmol) and LiOH (65 mg, 2.7 mmol) were dissolved in THF (5 mL) and water (5 mL). The mixture was stirred at 40°C for 6 h. TLC showed **11a** completely disappeared. The mixture was cooled to 0°C and the pH was slowly adjusted to 4–5 with 1N HCl. The solvent was concentrated *in vacuo* and the residue was dissolve in DMF to give a solution **11b** (0.2 mM in DMF), which was used for next step without further purification. LC/MS: m/z 301 $[M+H]^+$.

***tert*-Butyl(*R*,*E*)-(3-((5-(3-(4-amino-3-(4-phenoxyphenyl)-1*H*-pyrazolo[3,4-*d*]pyrimidin-1-yl)piperidin-1-yl)-2-methyl-5-oxopent-3-en-2-yl)(methyl)amino)propyl)carbamate(11C).** To a flask was added compound **11c** (77 mg, 0.2 mmol), solution **11b** (1.5 mL, 0.3 mmol), HATU (114 mg, 0.3 mmol) and DIPEA (129 mg, 1 mmol) in DMF(3 mL). The mixture was stirred at room temperature for 30 min. Then the reaction mixture was extracted with ethyl acetate, and the organic layer was washed with water, dried over $Na_2SO_4$, and filtered. The filtrate was concentrated *in vacuo* and the residue was purified by flash column chromatography (0~10% MeOH in Ethyl Acetate) to give product **11c** as a white solid (96 mg, 72%). LC/MS: m/z 669 $[M+H]^+$.

***N*-(3-(((*E*)-5-((*R*)-3-(4-amino-3-(4-phenoxyphenyl)-1*H*-pyrazolo[3,4-*d*]pyrimidin-1-yl)piperidin-1-yl)-2-methyl-5-oxopent-3-en-2-yl)(methyl)amino)propyl)-2-((2-(2,6-dioxopiperidin-3-yl)-1,3-dioxoisoindolin-4-yl)amino)acetamide (IRC-DiMe).** In a 25 mL flask was added **11c** (28 mg, 0.04 mmol) in TFA/DCM (5 mL, 1/1). The mixture was stirred for 30 min at room temperature. LC-MS showed **11c** converted into **11d** completely. Then remove the solvent *in vacuo* to give product **11d** (22 mg, 95%), which was used for next step without further purification. To above **11d** was added compound **5a** (20 mg, 0.06 mol), HATU (23 mg, 0.06 mmol) and DIPEA (26 mg, 0.2 mmol) in DMF (3 mL). The mixture was stirred at room temperature for 30 min. Then the reaction mixture was concentrated *in vacuo* and the residue was purified by PrepHPLC with a reverse phase C18 column to afford the product as a yellow solid I**RC-DiMe** (10 mg, 29%). $^1$H NMR (400 MHz, *D$_6$*-DMSO) δ 11.11 (s, 1H), 8.24 (s, 1H), 8.17 (s, 1H), 8.08 (m, 1H), 7.64 (s, 2H), 7.54 (m, 1H), 7.43 (t, *J* = 7.5 Hz, 2H), 7.22 – 7.09 (m, 5H), 7.04 (m, 1H), 6.94 (m, 1H), 6.83 (d, *J* = 8.5 Hz, 1H), 6.59 (m, 1H), 6.32 (m Hz, 1H), 5.07 (dd, *J* = 12.8, 5.2 Hz, 1H), 4.71 (m, 1H), 4.06 (m, 3H), 3.90 (m, 2H), 3.10 (m, 3H), 2.94 – 2.82 (m, 2H), 2.58 (m, 3H), 2.24 (m, 3H), 2.10 (m, 2H), 2.01 (m, 2H), 1.48 (m, 3H), 1.11 (m, 3H), 0.96 (m, 3H). $^{13}$C NMR (100 MHz, *D$_6$*-DMSO) δ 173.2, 170.5, 169.1, 168.7, 167.7, 165.4, 163.8, 158.6, 157.5, 156.7, 156.0, 154.4, 152.4, 146.2, 136.6, 132.5, 130.6, 130.5, 130.4, 128.4, 119.4, 118.9, 117.8, 110.4, 110.3, 97.9, 58.8, 52.5, 49.8, 49.0, 46.2, 45.7, 37.4, 34.9, 31.4, 30.1, 29.2, 28.2, 25.3, 23.0, 22.6. HRMS (m/z): $[M+H]^+$ calcd. for $C_{47}H_{52}N_{11}O_7$, 882.4051; found: 882.4065.
LC/MS: m/z 882 $[M+H]^+$.

**Procedure of DD-03-171 synthesis**

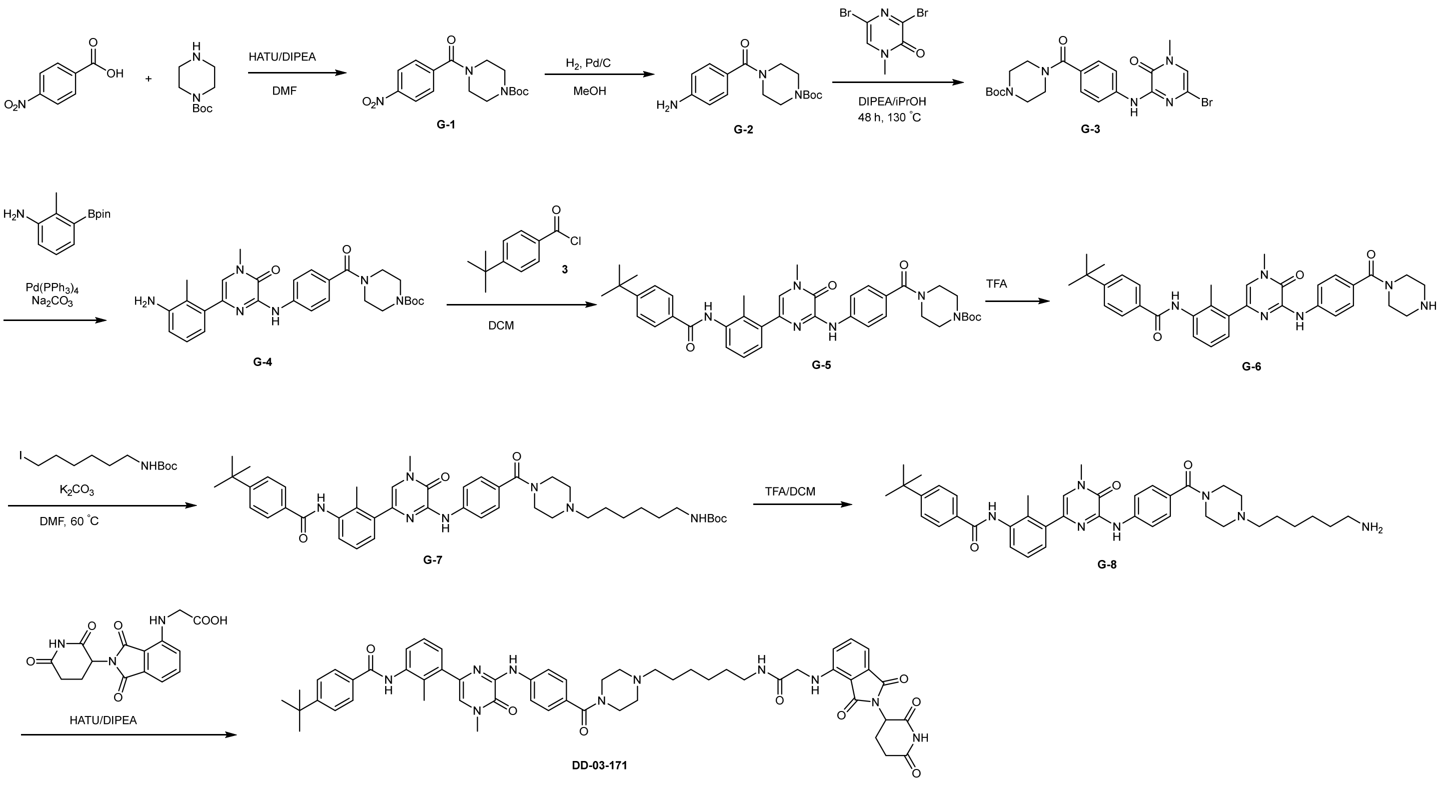

***tert*-Butyl 4-(4-nitrobenzoyl)piperazine-1-carboxylate(G-1).** To a flask was added 4-nitrobenzoic acid (835 mg, 5 mmol), *tert*-butyl piperazine-1-carboxylate (930 mg, 5 mmol), HATU (2850 mg, 7.5 mmol) and DIPEA (1935 mg, 15 mmol) in DMF(20 mL). The mixture was stirred at room temperature for 30 min. Then the reaction mixture was extracted with ethyl acetate, and the organic layer was washed with water, dried over $Na_2SO_4$, and filtered. The filtrate was concentrated *in vacuo* and the residue was purified by flash column chromatography (0~20% Ethyl Acetate in Hexane) to give product **G-1** as a white solid (1.26 g, 75%).$^1$H NMR (400 MHz, $CDCl_3$) δ 8.29 (d, *J* = 8.7 Hz, 2H), 7.57 (d, *J* = 8.7 Hz, 2H), 3.76 (s, 2H), 3.54 (s, 2H), 3.37 (m, 4H), 1.47 (s, 9H). LC/MS: m/z 336 $[M+H]^+$.

***tert*-Butyl 4-(4-aminobenzoyl)piperazine-1-carboxylate (G-2).** To a flask was added **G-1** (670 mg, 2 mmol) and 10%Pd/C (67 mg, 10%) in MeOH (20 mL). The mixture was stirred under 1 atm $H_2$ overnigh. LC-MS showed **G-1** converted into **G-2** completely. Then the reaction mixture was filtered, and the filtrate was concentrated *in vacuo* and the residue was purified by flash column to give product **G-2** as a white solid (550 mg, 90%). $^1$H NMR (400 MHz, $CDCl_3$) δ 7.26 – 7.22 (m, 2H), 6.65 (d, *J* = 8.4 Hz, 2H), 3.89 (s, 2H), 3.58 (s, 4H), 3.44 (s, 4H), 1.46 (s, 9H). LC/MS: m/z 306 $[M+H]^+$.

***tert*-Butyl4-(4-((6-bromo-4-methyl-3-oxo-3,4-dihydropyrazin-2-yl)amino)benzoyl)piperazine-1-carboxylate (G-3).** A solution of 3,5-dibromo-1-methylpyrazin-2(1*H*)-one (265 mg, 1 mmol), G-2 (427 mg, 1.4 mmol), and DIPEA (387 mg, 3 mmol) in isopropanol (5 mL) was stirred in a sealed Schlenk tube at 130 ˚C for 48 hours. Then the reaction mixture was cooled to room temperature and extracted with $CH_2Cl_2$. The organic layer was washed with brine, dried over $Na_2SO_4$, and filtered. The filtrate was concentrated *in vacuo* and the residue was purified by flash column chromatography to afford **G-3** (344 mg, 70%) as a white solid. $^1$H NMR (400 MHz, $CDCl_3$) δ 8.38 (s, 1H), 7.80 (d, *J* = 8.3 Hz, 2H), 7.43 (d, *J* = 8.2 Hz, 2H), 6.81 (s, 1H), 3.71 (m, 2H), 3.53 (s, 3H), 3.46 (m, 5H), 3.13 (m, 1H), 1.46 (s, 9H). LC/MS: m/z 492 $[M+H]^+$.

***tert*-Butyl 4-(4-((6-(3-amino-2-methylphenyl)-4-methyl-3-oxo-3,4-dihydropyrazin-2-yl)amino)benzoyl)piperazine-1-carboxylate(G-4).** To a 25 mL of Schlenk tube equipped with a magnetic stir bar were added 2-methyl-3-(4,4,5,5-tetramethyl-1,3,2-dioxaborolan-2-yl)aniline (82 mg, 0.35 mmol), **G-3** (172 mg, 0.59 mmol), $Na_2CO_3$ (74 mg, 0.7 mmol), $Pd(PPh_3)_4$ (40 mg, 10 mol%). Then dioxane/$H_2O$ (2.4 mL, v/v=5/1) was added under $N_2$. The Schlenk tube was screw capped and heated to 105 ˚C for 12 hours. Then the reaction mixture was cooled to room temperature, and sat. $NH_4Cl$ aq. was poured into the reaction mixture and extracted with EtOAc. The organic layer was washed with brine, dried over $Na_2SO_4$, and filtered. The filtrate was concentrated *in vacuo* and the residue was purified by flash column chromatography to afford **G-4** (70 mg, 39%) as a light yellow solid. LC/MS: m/z 519 $[M+H]^+$.

***tert*-Butyl 4-(4-((6-(3-(4-(tert-butyl)benzamido)-2-methylphenyl)-4-methyl-3-oxo-3,4-dihydropyrazin-2-yl)amino)benzoyl)piperazine-1-carboxylate(G-5).** To a solution of **G-4** (70 mg, 0.135 mmol) in dry DCM (5 mL) was added pyridine (16 μL, 0.2 mmol) and 4-*tert*--butyl benzoyl chloride (32 mg, 0.162 mmol). After stirring at room temperature for 2 h, remove the solvent *in vacuo* and the residue was purified by flash column chromatography to afford **G-5** (55 mg, 60%) as a white solid. LC/MS: m/z 679 $[M+H]^+$.

***tert*-Butyl (6-(4-(4-((6-(3-(4-(tert-butyl)benzamido)-2-methylphenyl)-4-methyl-3-oxo-3,4-dihydropyrazin-2-yl)amino)benzoyl)piperazin-1-yl)hexyl)carbamate(G-7).** In a 25 mL flask was added **G-5** (27 mg, 0.04 mmol) in TFA/DCM (3 mL, v/v=1/1). The mixture was stirred for 30 min at room temperature. LC-MS showed **G-5** converted into **C-6** completely. Then remove the solvent *in vacuo* to give product **G-6** (21 mg, 90%), which was used for next step without further purification. To above solution of **G-6** in DMF (2 mL), add $K_2CO_3$ (25 mg, 0.18 mmol) and stirred for 15 min, then *tert*-butyl (6-iodohexyl)carbamate (23 mg, 0.07 mmol) was added to the mixture. The mixture was stirred at 60 ˚C for 2 hour. Then the reaction mixture was concentrated *in vacuo* and the residue was purified by flash column chromatography to afford **G-7** (14 mg, 50%) as a colorless oil. $^1$H NMR (400 MHz, $CDCl_3$) δ 8.43 (s, 1H), 7.95 (d, *J* = 8.0 Hz, 1H), 7.88 (d, *J* = 8.3 Hz, 2H), 7.85 (d, *J* = 8.0 Hz, 2H), 7.54 (d, *J* = 8.3 Hz, 2H), 7.39 (d, *J* = 8.4 Hz, 2H), 7.31 (t, *J* = 7.7 Hz, 1H), 7.22 (d, *J* = 7.6 Hz, 1H), 6.77 (s, 1H), 4.52 (m, 1H), 3.64 (m, 2H), 3.63 (s, 3H), 3.48 (m, 1H), 3.09 (m, 2H), 2.49 (m, 4H), 2.40 (m, 2H), 2.38 (s, 3H), 1.43 (s, 9H), 1.36 (s, 9H), 1.52-1.21(m, 6H). LC/MS: m/z 778 $[M+H]^+$.

**4-(*tert*-Butyl)-*N*-(3-(6-((4-(4-(6-(2-((2-(2,6-dioxopiperidin-3-yl)-1,3-dioxoisoindolin-4-yl)amino)acetamido) hexyl)piperazine-1-carbonyl)phenyl)amino)-4-methyl-5-oxo-4,5-dihydropyrazin-2-yl)-2-methylphenyl) benzamide(DD-03-171).** In a 25 mL flask was added **G-7** (14 mg, 0.018 mmol) in TFA/DCM (5 mL, 1/1). The mixture was stirred for 30 min at room temperature. LC-MS showed **G-7** converted into **G-8** completely. Then remove the solvent *in vacuo* to give product **G-8** (12 mg, 95%), which was used for next step without further purification. To above **G-8** was added compound **5a** (10 mg, 0.03 mol), HATU (12 mg, 0.03 mmol) and DIPEA (13 mg, 0.1 mmol) in DMF (3 mL). The mixture was stirred at room temperature for 30 min. Then the reaction mixture was concentrated *in vacuo* and the residue was purified by PrepHPLC with a reverse phase C18 column to afford the product as a yellow solid **DD-03-171** (7 mg, 42%). $^1$H NMR (400 MHz, *D*$_6$-DMSO) δ 11.12 (s, 1H), 9.92 (s, 1H), 9.45 (s, 1H), 8.11 (s, 1H), 8.07 (d, *J* = 8.3 Hz, 2H), 7.95 (d, *J* = 8.0 Hz, 2H), 7.57 (m, 3H), 7.36 (m, 1H), 7.29 (m, 5H), 7.06 (d, *J* = 6.9 Hz, 1H), 6.94 (t, *J* = 5.8 Hz, 1H), 6.85 (d, *J* = 8.4 Hz, 1H), 5.07 (dd, *J* = 12.7, 5.2 Hz, 1H), 3.91 (d, *J* = 4.6 Hz, 2H), 3.56 (s, 3H), 3.07 (m, 6H), 2.88 (m, 1H), 2.57 (m, 2H), 2.32 (m, 3H), 2.28 (m, 3H), 2.24 (m, 3H), 2.01 (m, 1H), 1.38 (m, 2H), 1.32 (m, 9H), 1.23 (s, 6H). $^{13}$C NMR (100 MHz, *D*$_6$-DMSO) δ 172.9, 170.1, 168.9, 168.7, 168.3, 167.4, 165.3, 154.4, 150.5, 146.3, 145.9, 138.3, 137.2, 136.2, 132.6, 132.1, 131.8, 131.2, 129.0, 127.9, 127.6, 127.2, 126.6, 125.5, 125.2, 120.5, 118.6, 117.5, 111.0, 109.9, 57.7, 52.8, 48.7, 48.6, 45.2, 38.6, 36.7, 34.7, 31.01, 31.00, 29.1, 26.7, 26.3, 26.2, 22.2, 15.6. LC/MS: m/z 991 $[M+H]^+$.

**Procedure of MT-802 synthesis**

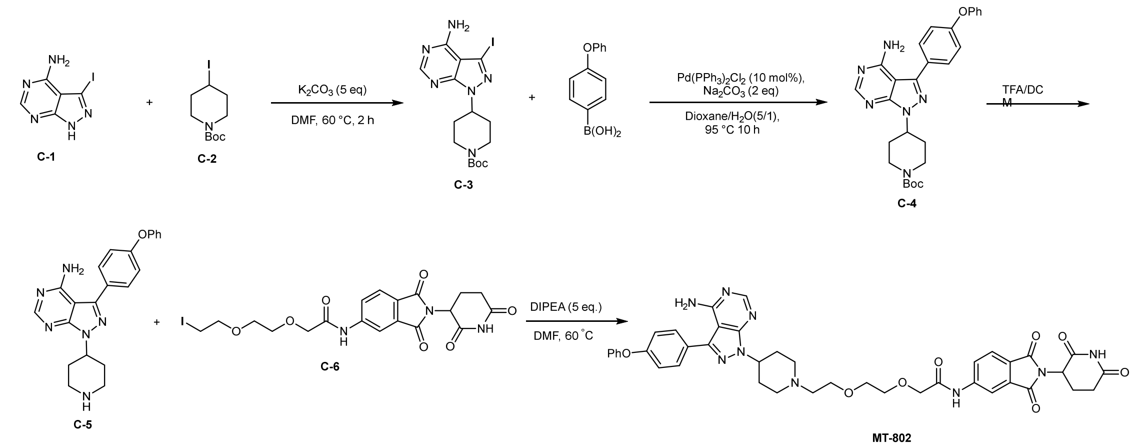

***tert*-Butyl 4-(4-amino-3-iodo-1*H*-pyrazolo[3,4-*d*]pyrimidin-1-yl)piperidine-1-carboxylate (C-3).** To a solution of **C-1** (260 mg, 1 mmol) in 5 mL DMF, add $K_2CO_3$ (690 mg, 5 mmol) and stirred for 15 min, then **C-2** (622 mg, 2 mmol) was added to the mixtrue. The solution was stirred for 2 hours at 60 ˚C. Then the reaction mixture was concentrated *in vacuo* and the residue was purified by flash column chromatography (5% MeOH in DCM) to give product **C-3** (260 mg, 59%) as a white solid. LC/MS: m/z 445 $[M+H]^+$.

***tert*-Butyl 4-(4-amino-3-(4-phenoxyphenyl)-1*H*-pyrazolo[3,4-*d*]pyrimidin-1-yl)piperidine-1-carboxylate (C-4)**. To a 25 mL of Schlenk tube equipped with a magnetic stir bar were added (4-phenoxyphenyl)boronic acid (152 mg, 0.71 mmol), **C-3** (260 mg, 0.59 mmol), $K_3PO_4$ (250 mg, 1.18 mmol), Pd(dppf)$Cl_2$ (24 mg, 5 mol%). Then DMF/$H_2O$ (5 mL, v/v=3/2) was added under $N_2$. The Schlenk tube was screw capped and heated to 130 ˚C for 12 hours. Then the reaction mixture was cooled to room temperature, and sat. $NH_4Cl$ aq. was poured into the reaction mixture and extracted with $CH_2Cl_2$. The organic layer was washed with brine, dried over $Na_2SO_4$, and filtered. The filtrate was concentrated *in vacuo* and the residue was purified by flash column chromatography to afford **C-4** (88 mg, 80%) as a light yellow solid. $^1$H NMR (400 MHz, *$D_6$*-DMSO) δ 8.24 (s, 1H), 7.65 (d, *J* = 8.5 Hz, 2H), 7.43 (t, *J* = 7.6 Hz, 2H), 7.21 – 7.08 (m, 5H), 4.88 (m, 1H), 4.07 (br, 2H), 2.98 (br, 2H), 2.06 – 1.86 (m, 4H), 1.42 (s, 9H).

**2-(2-(2-(4-(4-Amino-3-(4-phenoxyphenyl)-1*H*-pyrazolo[3,4-*d*]pyrimidin-1-yl)piperidin-1-yl)ethoxy)ethoxy)-N-(2-(2,6-dioxopiperidin-3-yl)-1,3-dioxoisoindolin-5-yl)acetamide (MT-802).** In a 25 mL flask was added **C-4** (8 mg, 0.02 mmol) in TFA/DCM (3 mL, v/v=1/1). The mixture was stirred for 30 min at room temperature. LC-MS showed **C-4** converted into **C-5** completely. Then remove the solvent *in vacuo* to give product **C-5** (7.3 mg, 95%), which was used for next step without further purification. To above solution of **C-5** in DMF, add DIPEA (13 mg, 0.1 mmol) and stirred for 15 min, then **C-6** (16 mg, 0.03 mmol) was added to the mixture. The mixture was stirred at 60 ˚C for 1 hour. Then the reaction mixture was concentrated *in vacuo* and the residue was purified by PrepHPLC with a reverse phase C18 column to afford the product as a white solid MT-802 (8 mg, 51%). $^1$H NMR (400 MHz, $CDCl_3$) δ 9.82 (br, 1H), 9.48 (s, 1H), 8.39 (s, 1H), 8.27 (d, *J* = 8.2 Hz, 1H), 7.99 (s, 1H), 7.82 (d, *J* = 8.2 Hz, 1H), 7.61 (d, *J* = 8.5 Hz, 2H), 7.38 (t, *J* = 7.8 Hz, 2H), 7.15 (m, 3H), 7.07 (d, *J* = 7.8 Hz, 2H), 4.95 (dd, *J* = 12.1, 5.3 Hz, 1H), 4.86 (m, 1H), 4.16 (m, 2H), 3.85 – 3.70 (m, 6H), 3.32 – 3.17 (m, 2H), 2.93 – 2.72 (m, 5H), 2.49 (m, 4H), 2.20 – 2.12 (m, 1H), 2.05 (d, *J* = 8.6 Hz, 2H). $^{13}$C NMR (100 MHz, $CDCl_3$) δ 171.4, 169.0, 168.7, 167.0, 166.8, 158.5, 157.8, 156.3, 155.0, 153.5, 143.8, 143.5, 133.0, 130.0, 129.9, 127.7, 126.5, 125.1, 124.6, 124.0, 119.5, 119.1, 114.8, 98.5, 71.4, 70.5, 70.0, 68.3, 57.1, 53.4, 52.8, 50.9, 49.4, 31.5, 30.5, 30.4, 22.8. LC/MS: m/z 789.3 $[M+H]^+$.

**Supplementary Figures and Tables**

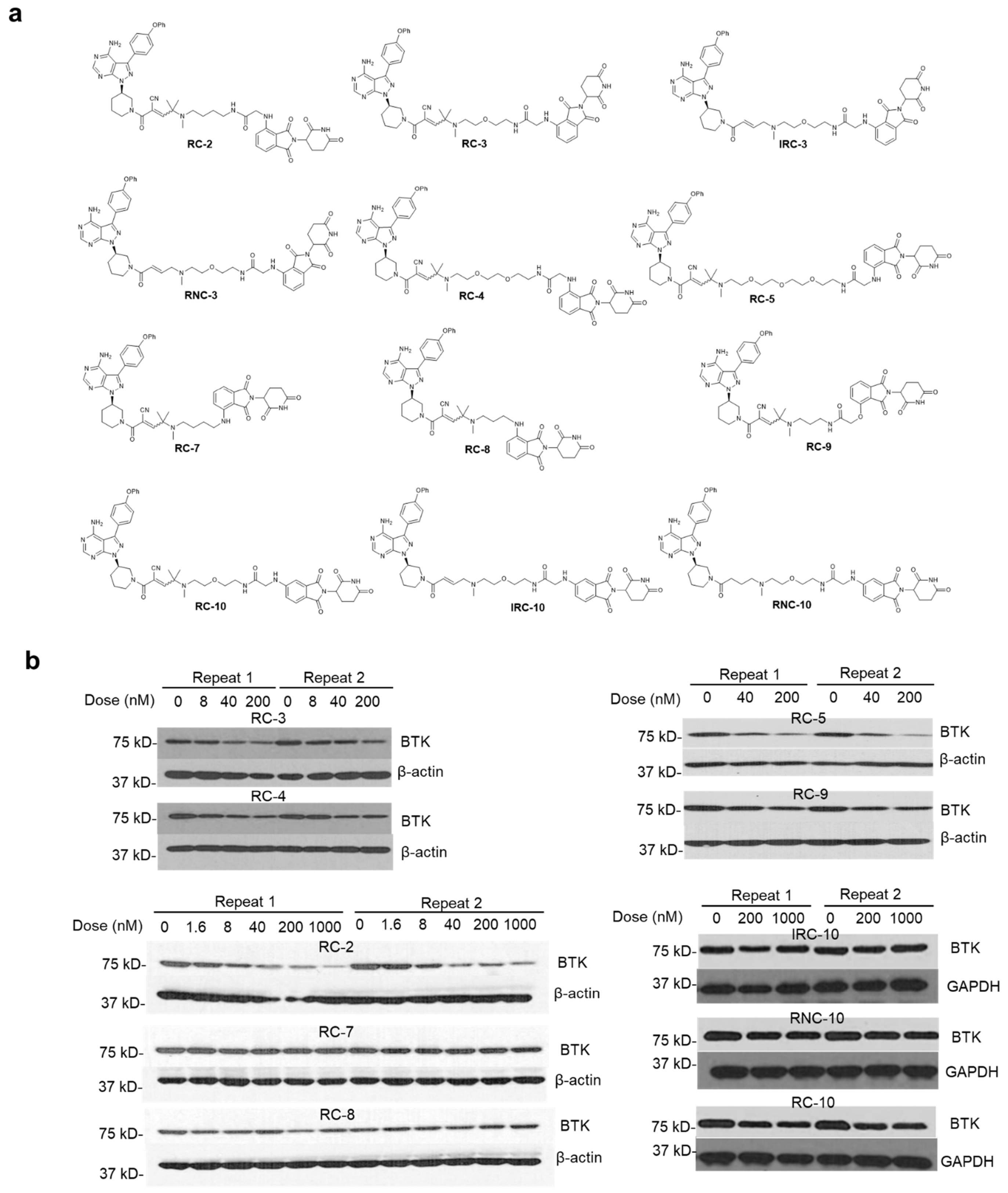


**Supplementary Figure 1. BTK degradation induced by PROTACs.** (a) Chemical structures of BTK degraders. (b) Western blot from MOLM-14 cells treated with the indicated doses of PROTACs for 24 h. Duplicates were performed. Source data are provided as a Source Data file.

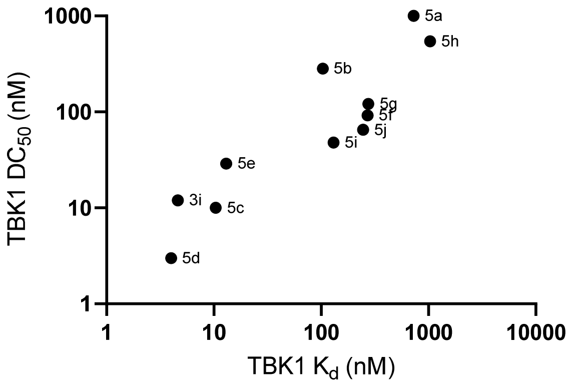


**Supplementary Figure 2. The correlation between $K_d$ towards TBK-1 and $DC_{50}$.**

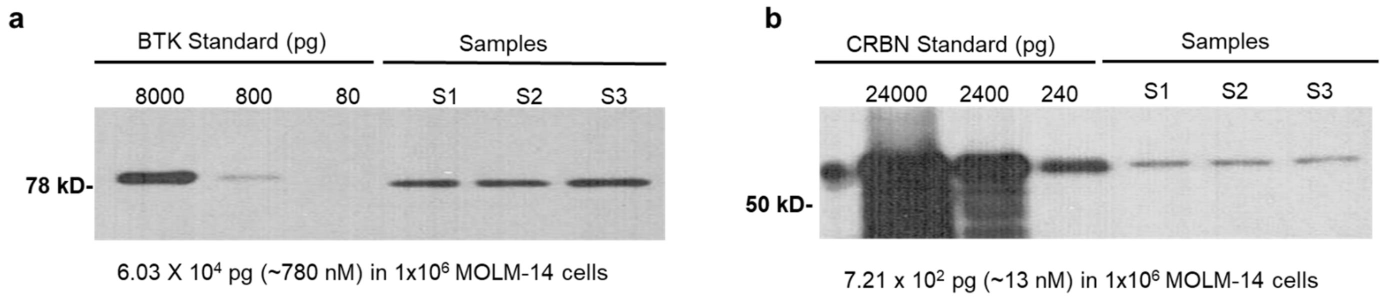


**Supplementary Figure 3. Quantification of BTK and CRBN levels in MOLM-14 cells.** MOLM-14 cells were maintained in RPMI-1640 complete culture medium. $1x10^6$ cells were collected and processed for Western blot analysis of BTK and CRBN levels using their corresponding primary and secondary antibodies. (a) BTK and (b) CRBN were quantified by normalizing the sample BTK and CRBN to their standard recombinant BTK (SignalChem. Cat. No.  B10-10H-10) and CRBN (LifeSpan BioSciences, LS-G55983), respectively. The data shown are average of 3 repeat samples. Source data are provided as a Source Data file.

**a**

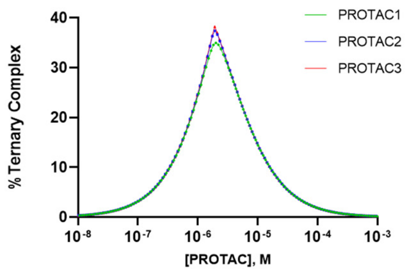


**b**

| | Input Parameters | | | Output Exact Values | | | |
|---|---|---|---|---|---|---|---|
| | PROTAC1 | PROTAC2 | PROTAC3 | | PROTAC1 | PROTAC2 | PROTAC3 |
| $[A]_t$/M | 1.90E-06 | 1.90E-06 | 1.90E-06 | $[B]_{t,max}$/M | 2.00E-06 | 1.93E-06 | 1.91E-06 |
| $k_{ab}$/M | 2.00E-08 | 2.00E-09 | 2.00E-10 | $[ABC]_{max,noncoop}$ | 1.12E-08 | 1.20E-08 | 1.22E-08 |
| $[C]_t$/M | 3.20E-08 | 3.20E-08 | 3.20E-08 | $[ABC]_{max,noncoop}/[L]_t$ | 34.99% | 37.42% | 38.24% |
| $k_{bc}$/M | 3.00E-06 | 3.00E-06 | 3.00E-06 | $TF_{50}$ (exact)/M | 6.54E-07 | 6.98E-07 | 7.15E-07 |
| α | 1.0 | 1.0 | 1.0 | $TI_{50}$ (exact)/M | 7.82E-06 | 7.14E-06 | 6.93E-06 |

**Supplementary Figure 4.** (a) Theoretical ternary complex formation curves of PROTACs for a non-cooperative system (α=1). (b) The input parameters and output exact values of PROTACs based on the ternary complex formation modeling.

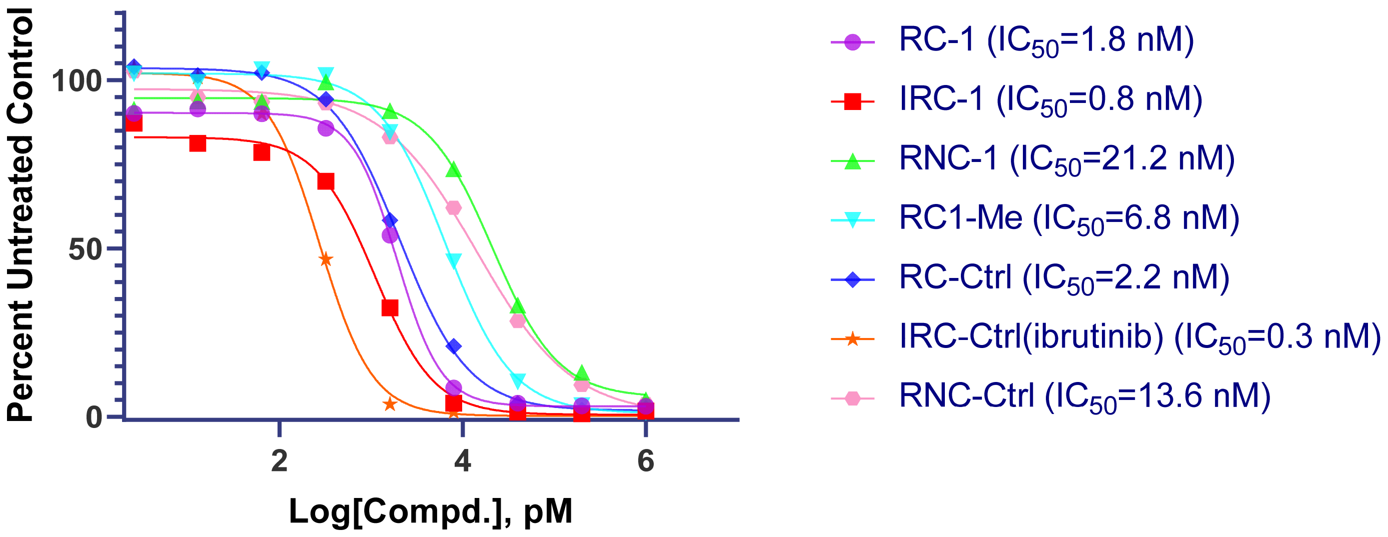


**Supplementary Figure 5. BTK kinase inhibition $IC_{50}$.** 9-point dose response curves were performed using PhosphoSens® Kinase Assay Kit. Each concentration point was performed in duplicate. Source data are provided as a Source Data file.

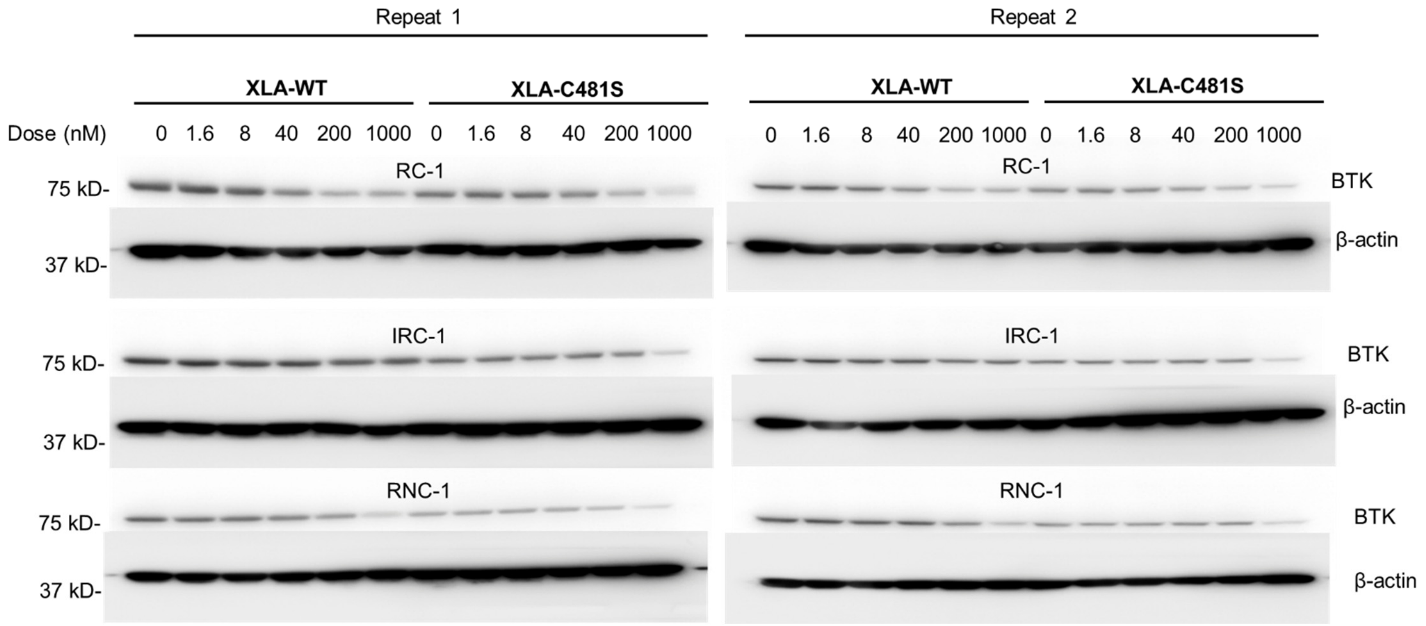


**Supplementary Figure 6. BTK degradation by three categories of PROTACs in** XLA cells overexpressing wild type BTK (XLA-WT) or mutant C481S BTK (XLA-C481S). Duplicates were performed. Source data are provided as a Source Data file.

a

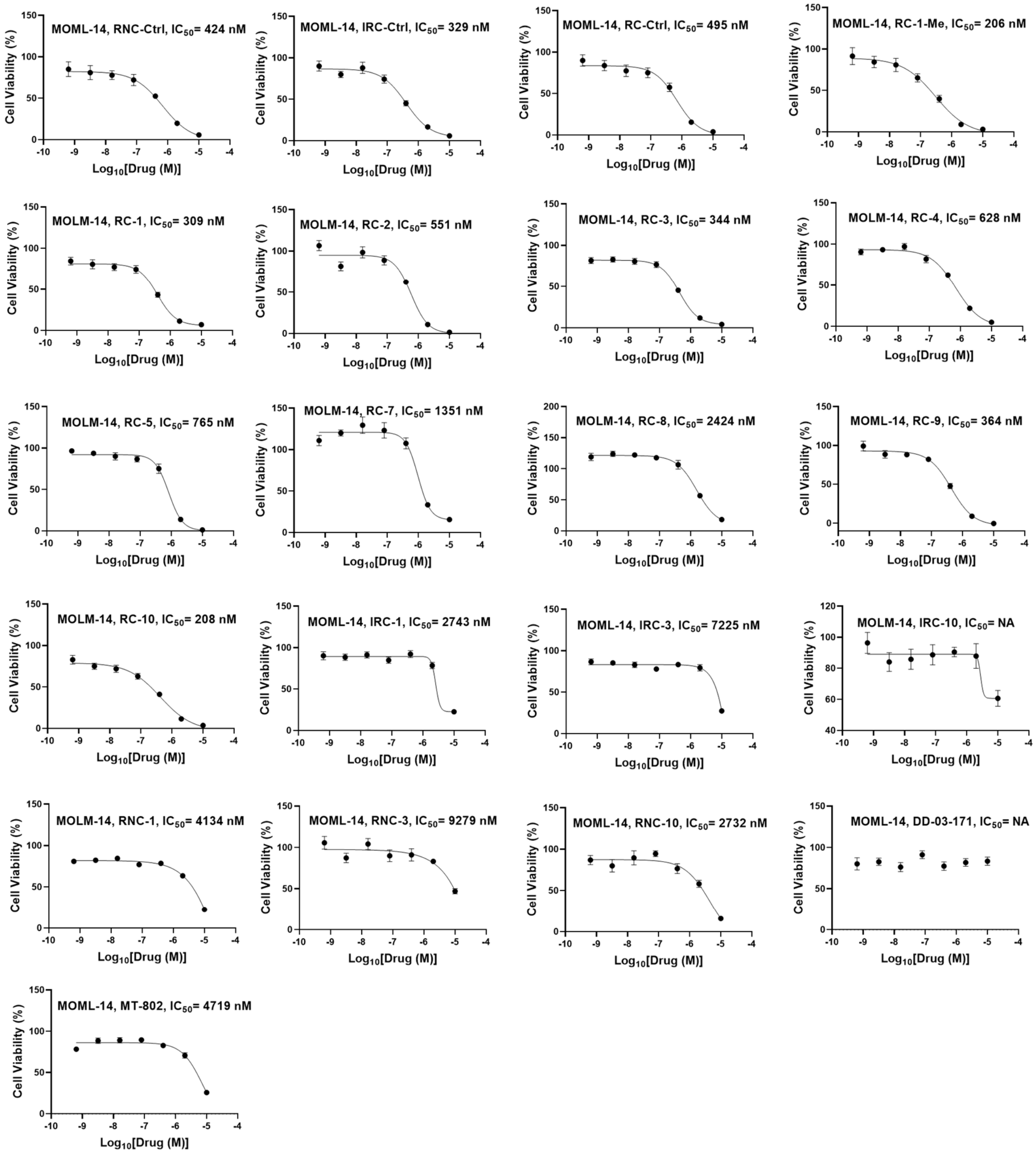


b

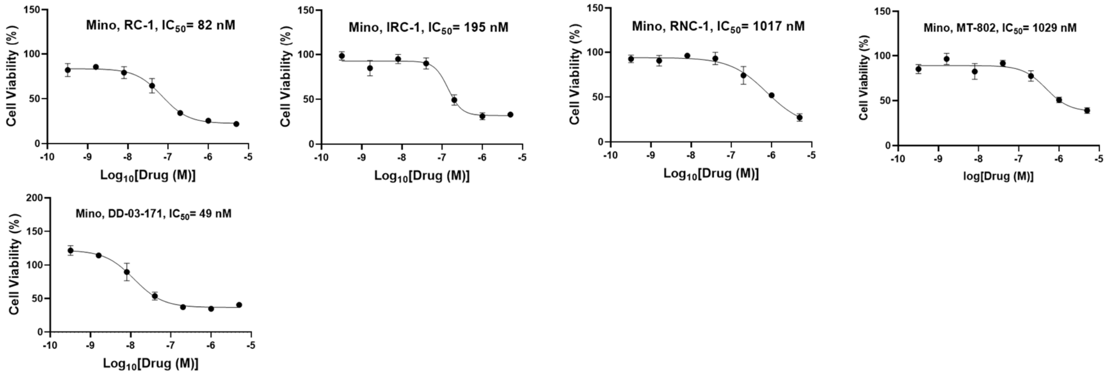


**Supplementary Figure 7. Cell viability assay following treatment with BTK degraders and their corresponding warhead controls.** (a) Cell viability in MOLM-14 cells. (b) Cell viability in Mino cells. Data are presented as mean values ± SEM (n = 5 biologically independent samples). Source data are provided as a Source Data file.

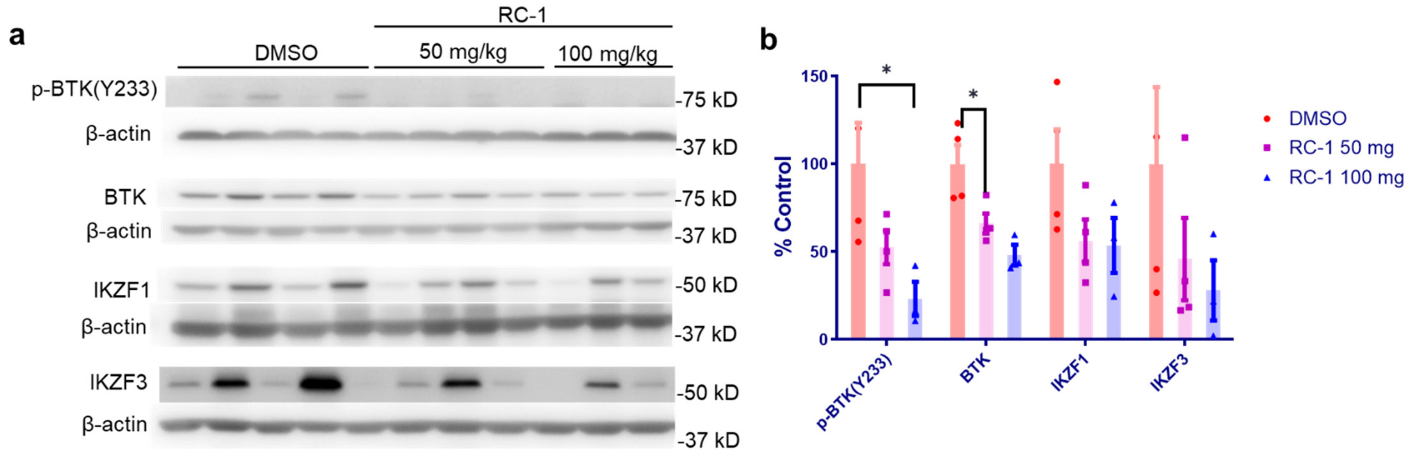


**Supplementary Figure 8. Protein degradation by BTK PROTAC RC-1 in mouse spleen.** ICR mice were subjected to single IP injection of RC-1 at the dose of 50 mg/kg (n = 4) or 100 mg/kg (n = 3) and spleens were harvested 24 h after injection. (a) The splenic p-BTK(Y233), BTK, IKZF1 and IKZF3 levels were measured with Western blot. (b). Quantification of protein levels shown in (a). Data are presented as mean values ± SEM. Asterisks indicate that the differences between samples are statistically significant, using two-tailed, unpaired t-test (* $p < 0.05$; significant). Source data are provided as a Source Data file.

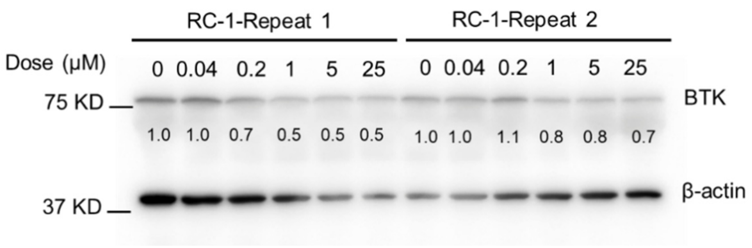


**Supplementary Figure 9. BTK degradation induced by RC-1 in mouse cell line.** E mu-myc transgenic mouse cells were incubated with RC-1 for 24 h. The BTK levels were quantified by Western blotting. Duplicates were performed. Source data are provided as a Source Data file.

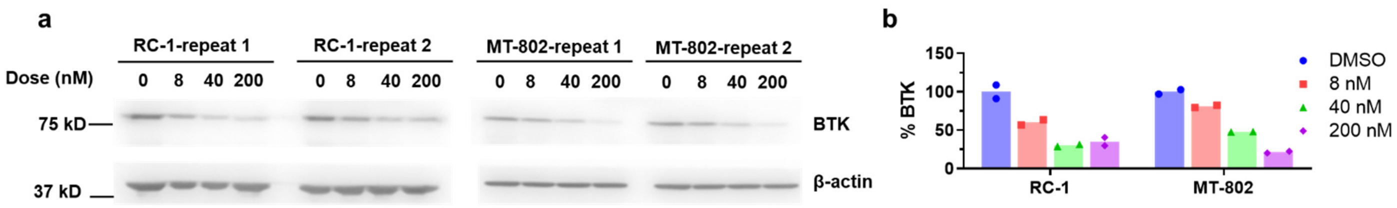


**Supplementary Figure 10. BTK degradation induced by RC-1 and MT-802.** (a) MOLM-14 cells were incubated with RC-1 and MT-802 for 24 h. The BTK levels were quantified by Western blotting. (b) Quantification of BTK levels shown in (a). Duplicates were performed. Source data are provided as a Source Data file.

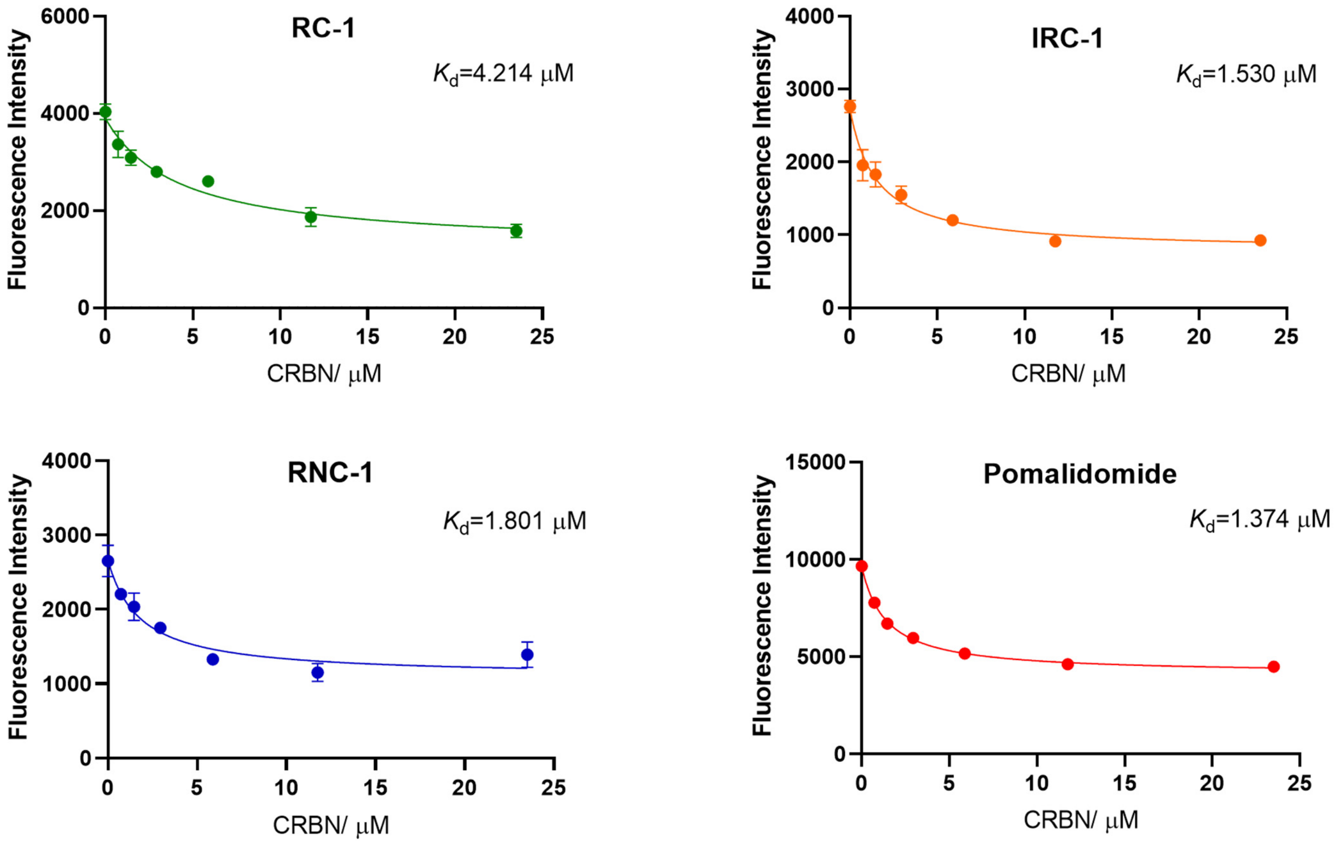


**Supplementary Figure 11. CRBN binding affinity.** The dissociation equilibrium constant $K_d$ was measured based on fluorescence quenching after compounds binding to CRBN. Data are presented as mean values ± SD (n = 3 biologically independent samples). Source data are provided as a Source Data file.

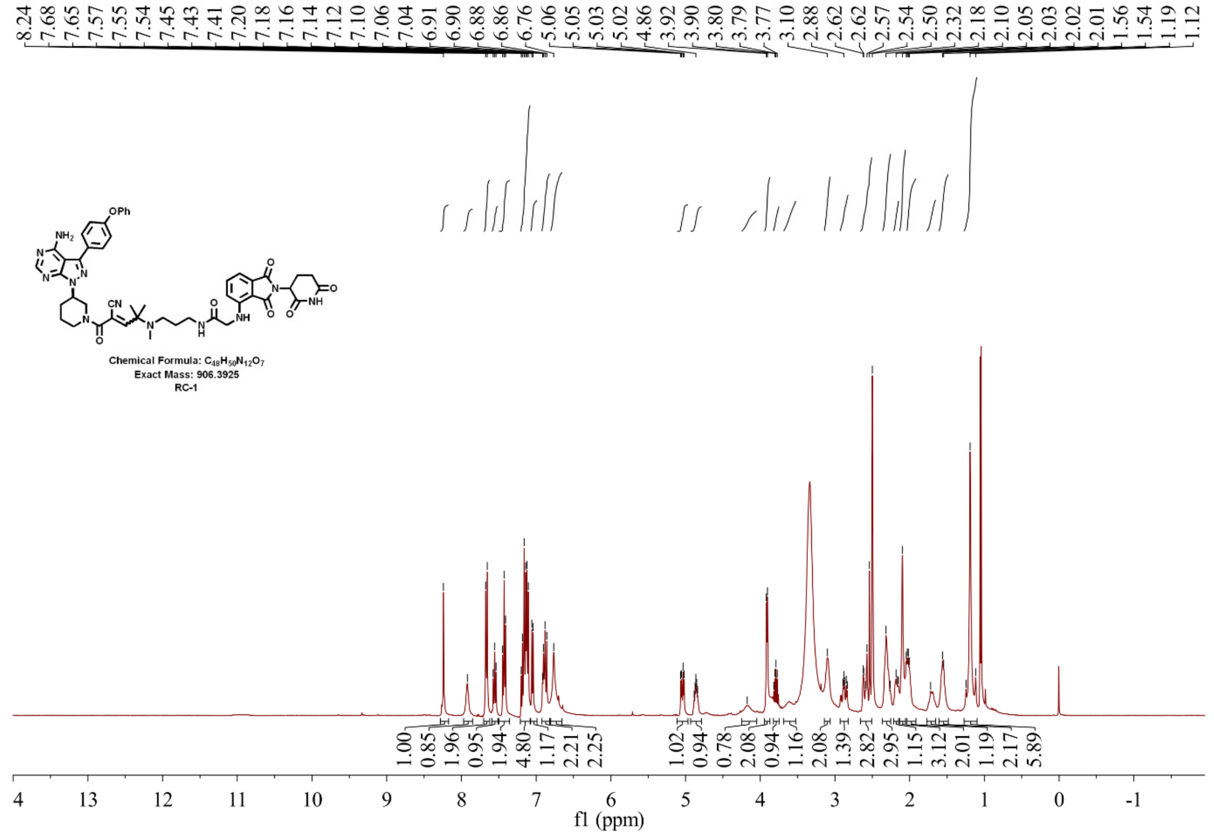


**Supplementary Figure 12. $^1$HNMR spectrum of RC-1**

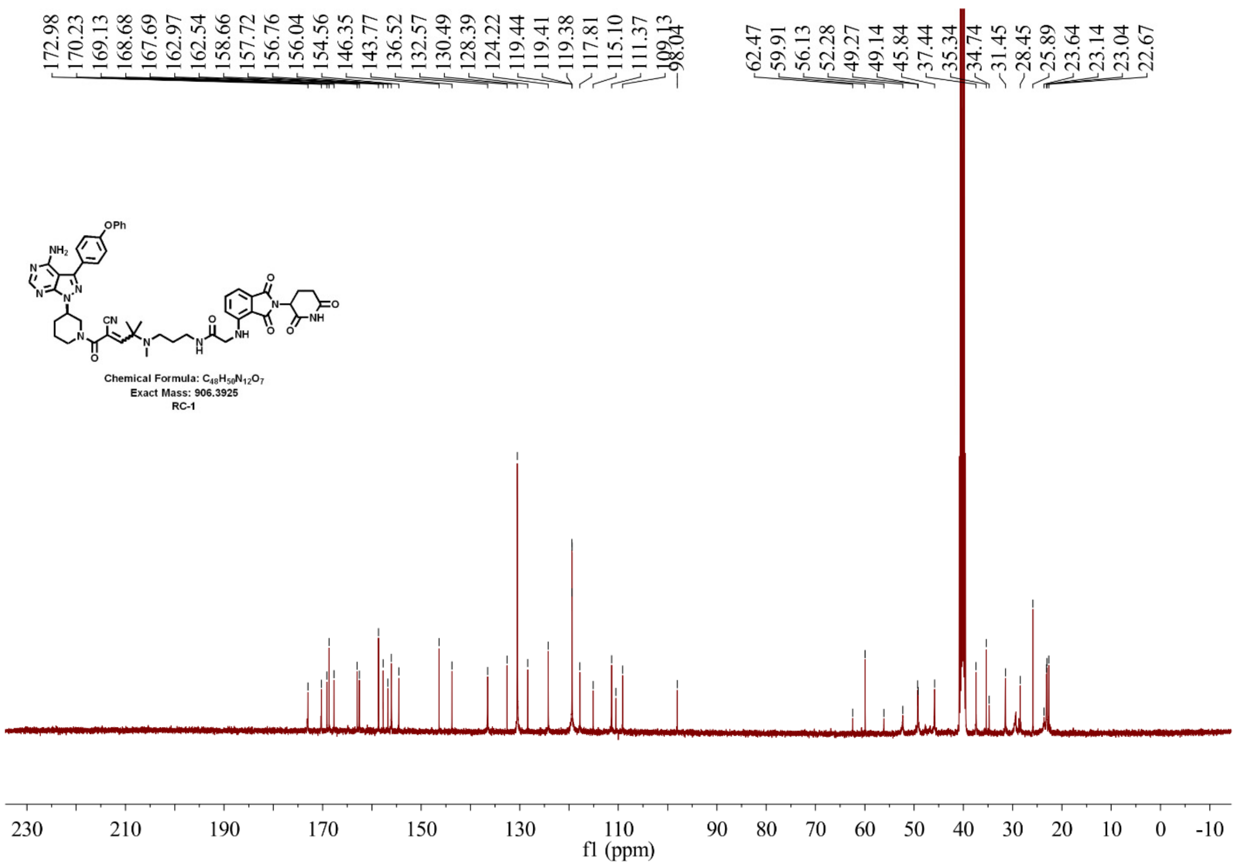


**Supplementary Figure 13. $^{13}$CNMR spectrum of RC-1**

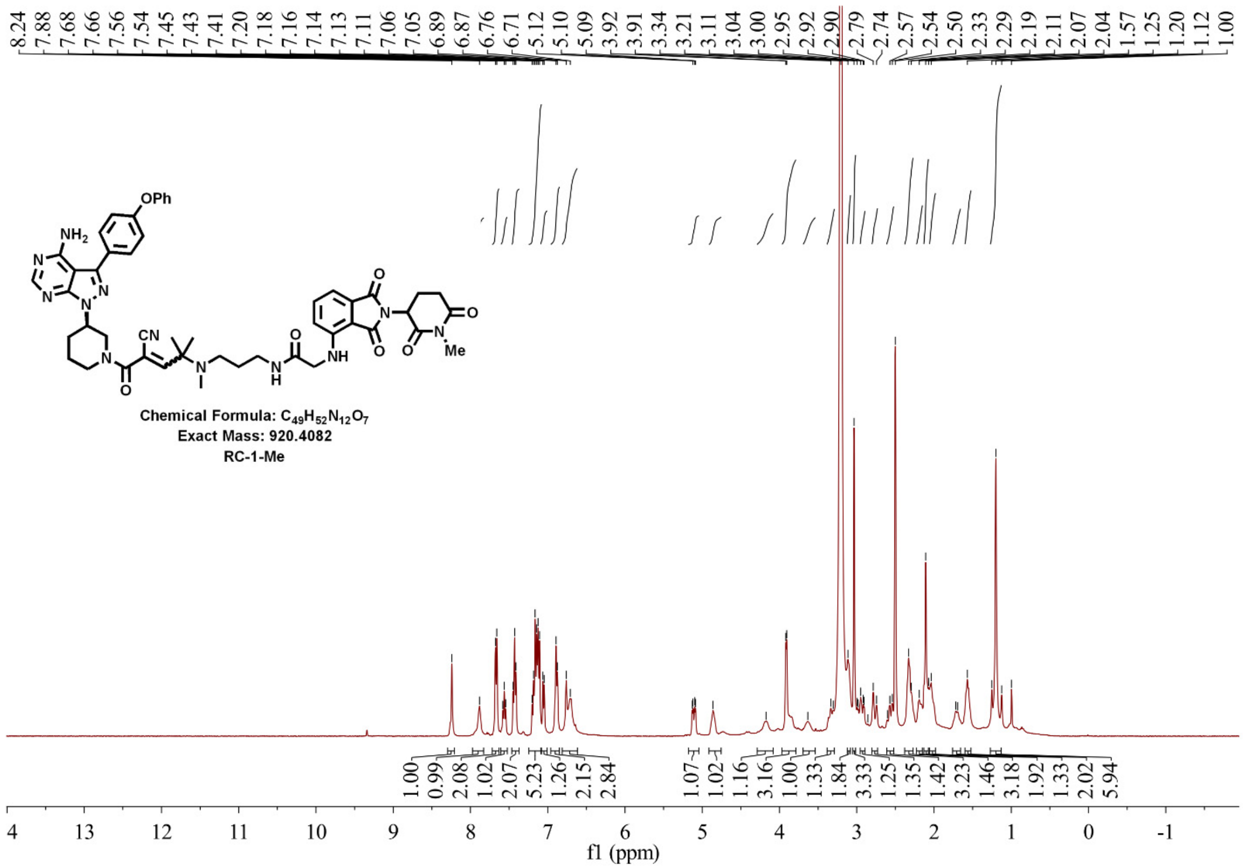


**Supplementary Figure 14. $^{1}$HNMR spectrum of RC-1-Me**

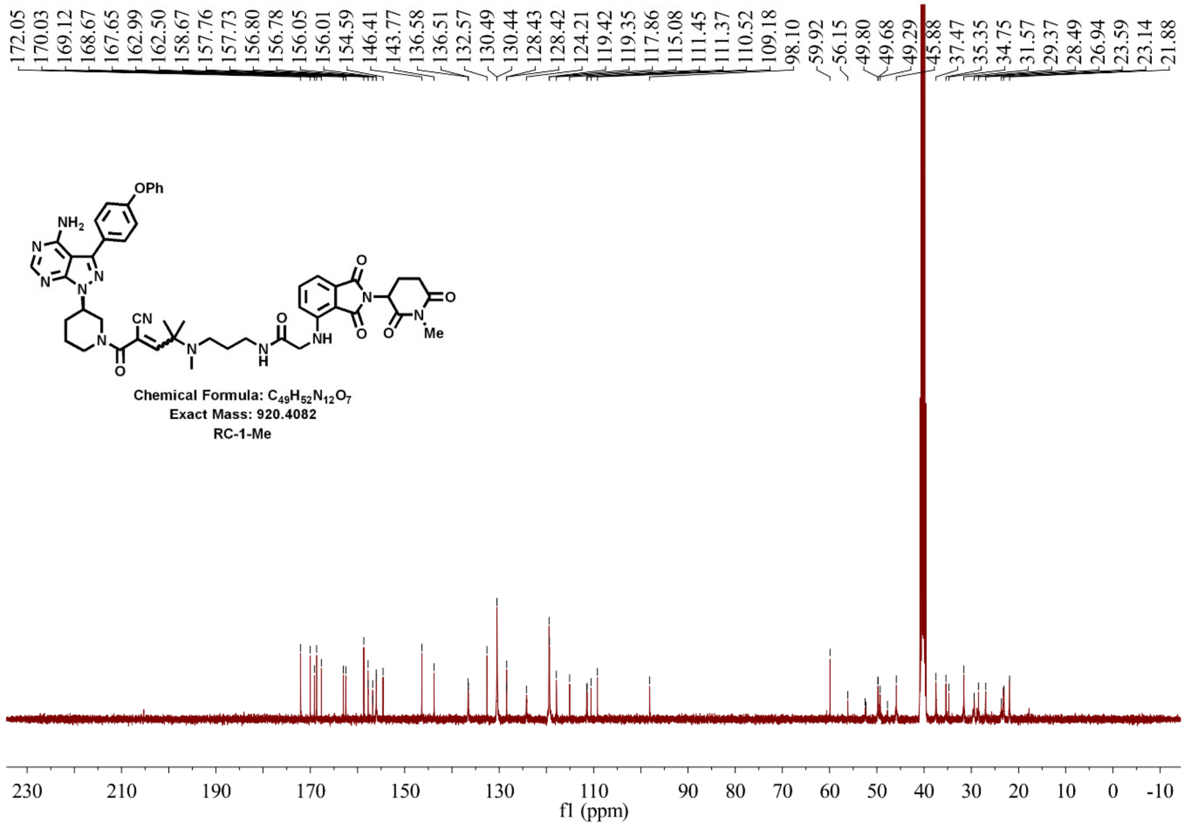


**Supplementary Figure 15. $^{13}$CNMR spectrum of RC-1-Me**

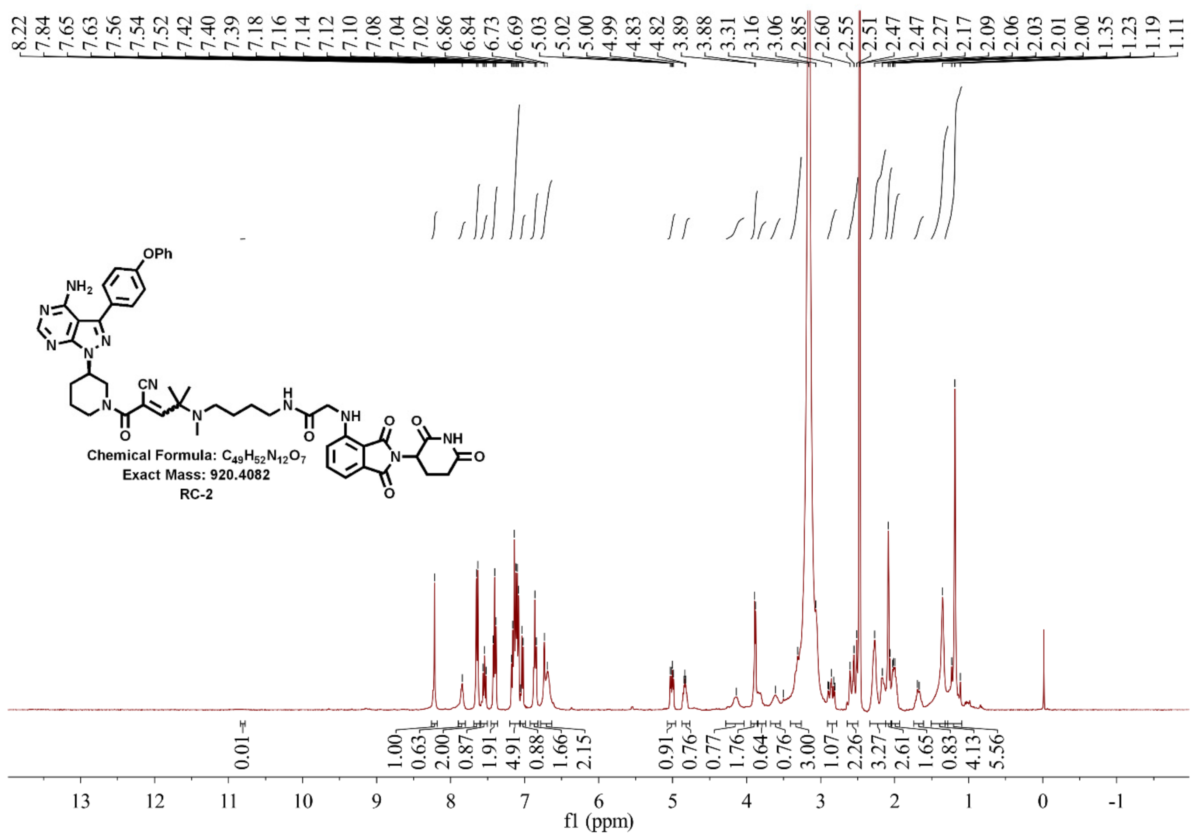

**Supplementary Figure 16. $^{1}$HNMR spectrum of RC-2**

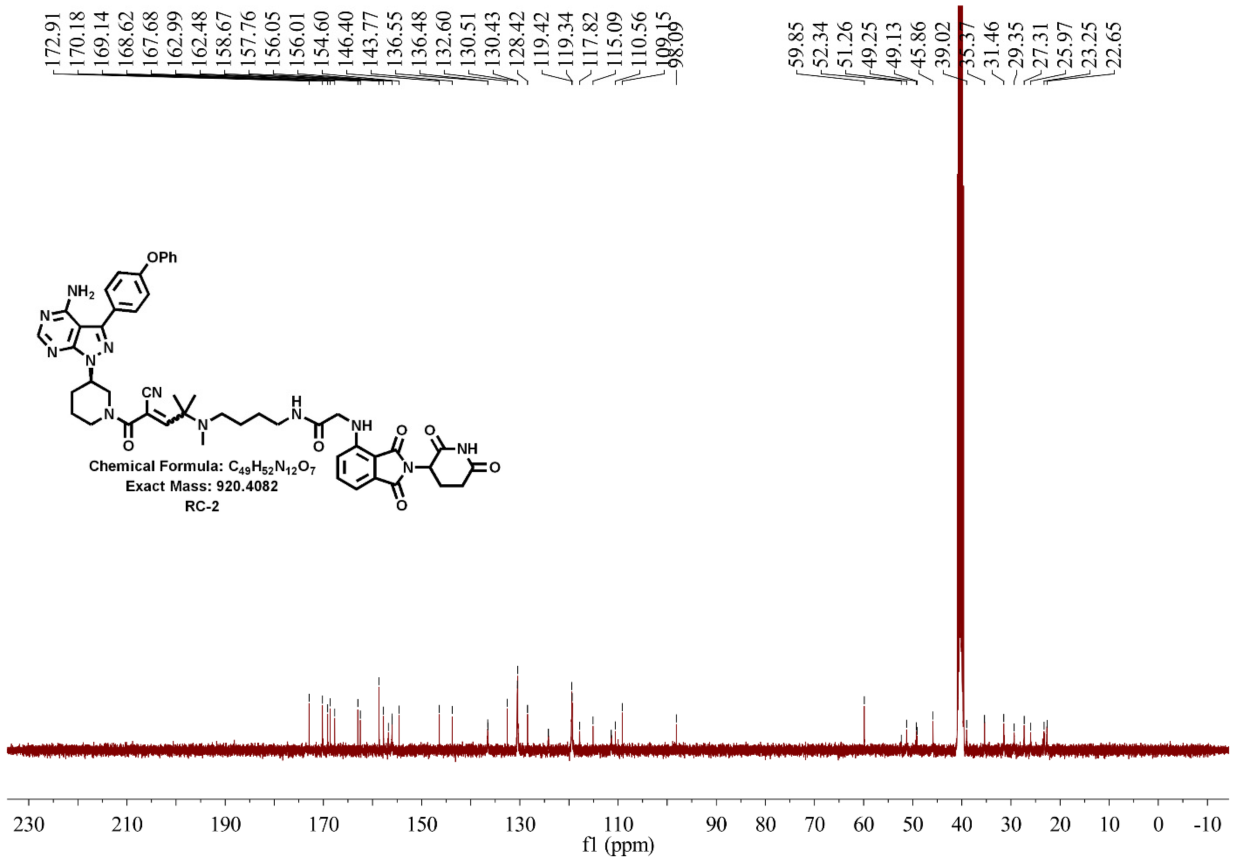

**Supplementary Figure 17. $^{13}$CNMR spectrum of RC-2**

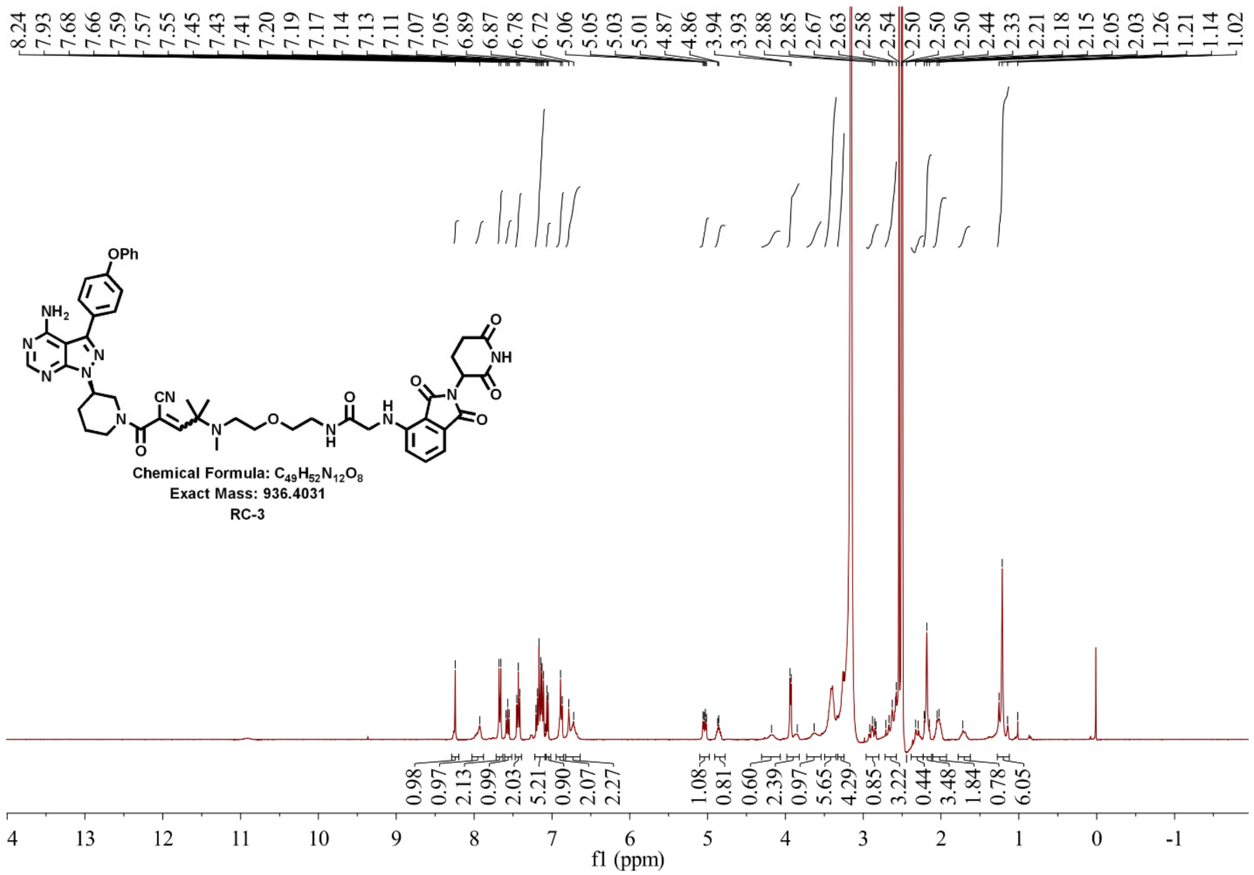


**Supplementary Figure 18. [1]HNMR spectrum of RC-3**

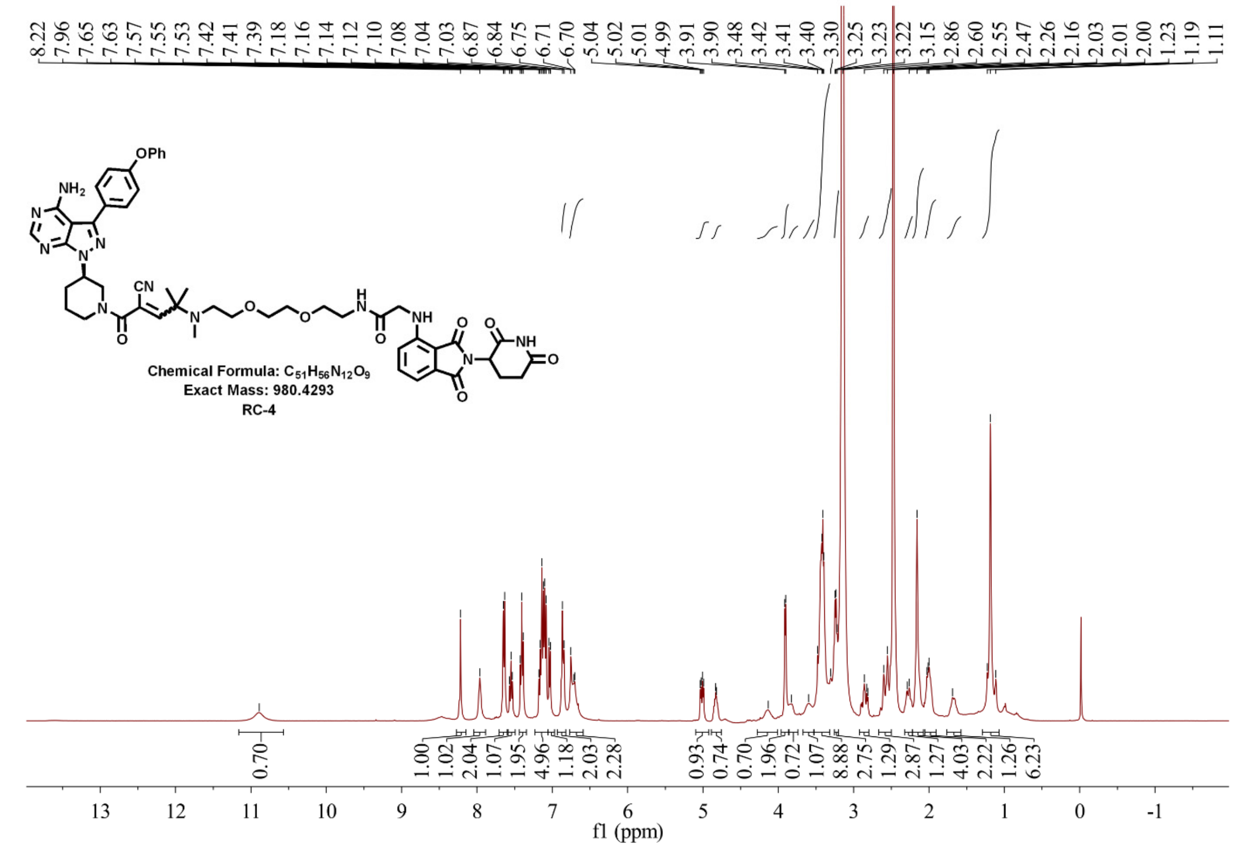


**Supplementary Figure 19. [1]HNMR spectrum of RC-4**

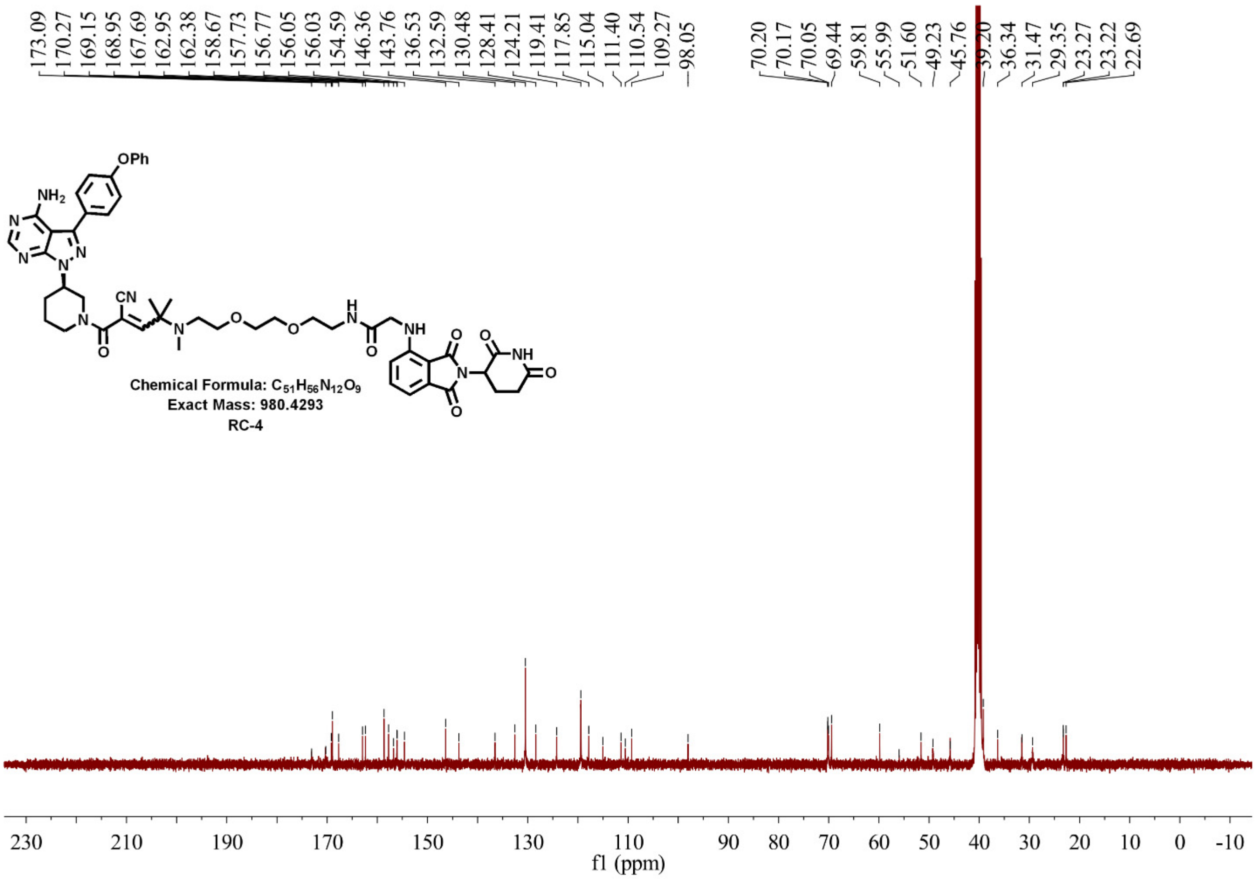


**Supplementary Figure 20. $^{13}$CNMR spectrum of RC-4**

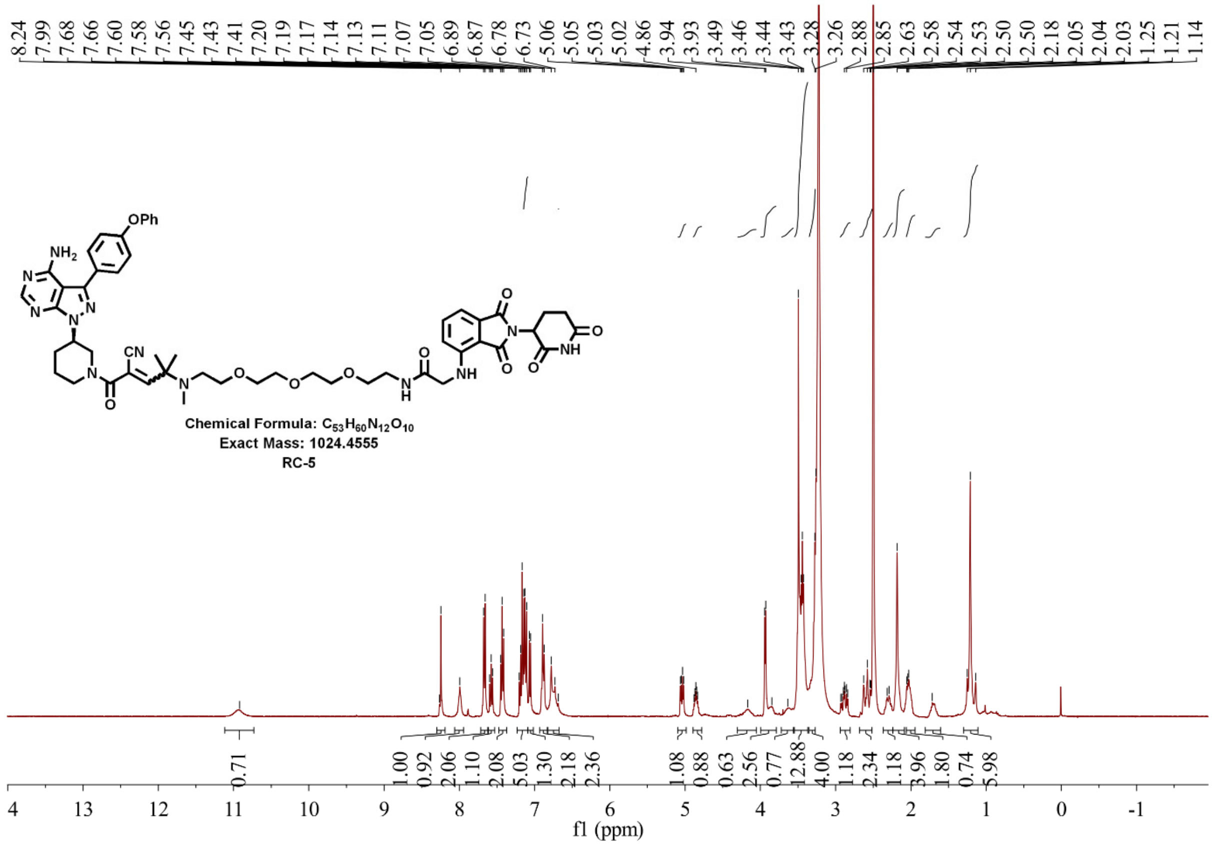


**Supplementary Figure 21. $^{1}$HNMR spectrum of RC-5**

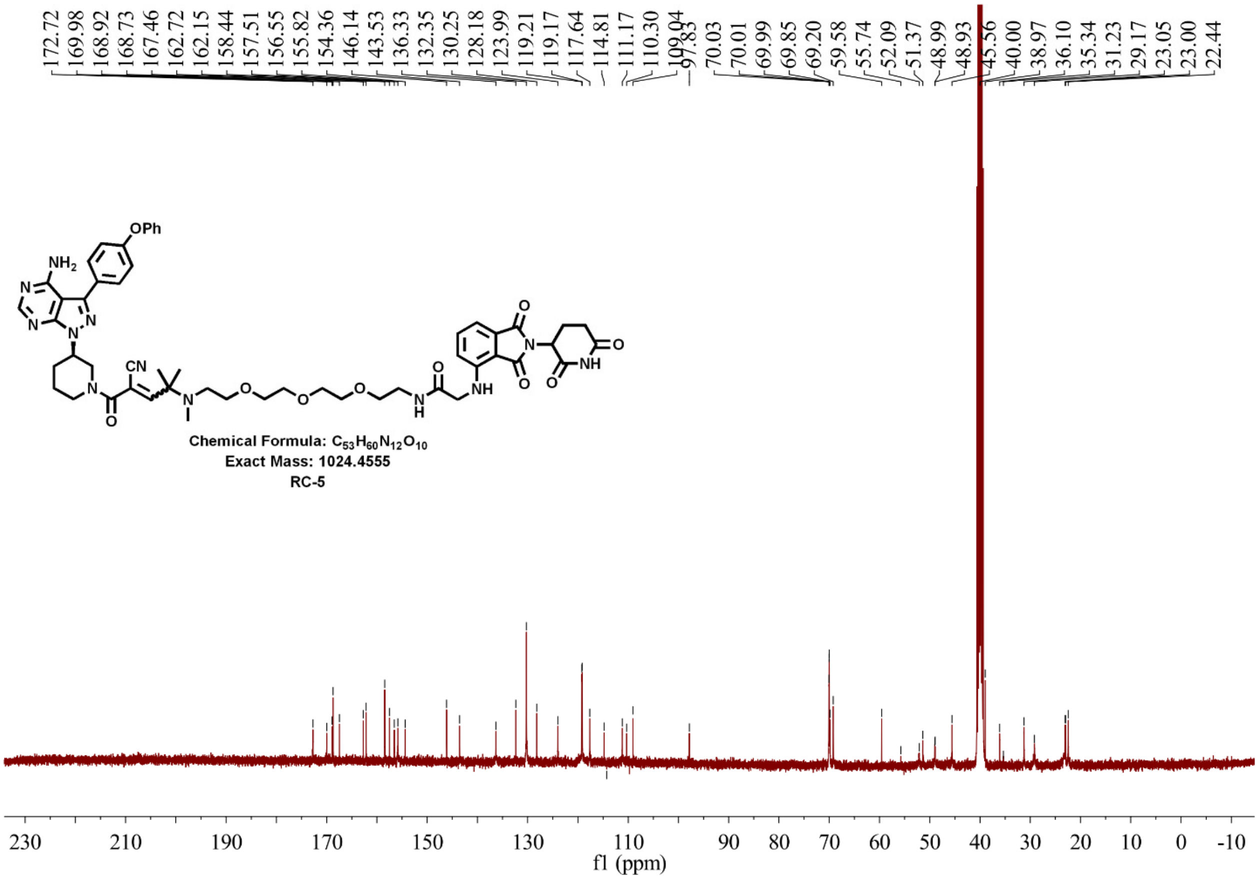


**Supplementary Figure 22. $^{13}$CNMR spectrum of RC-5**

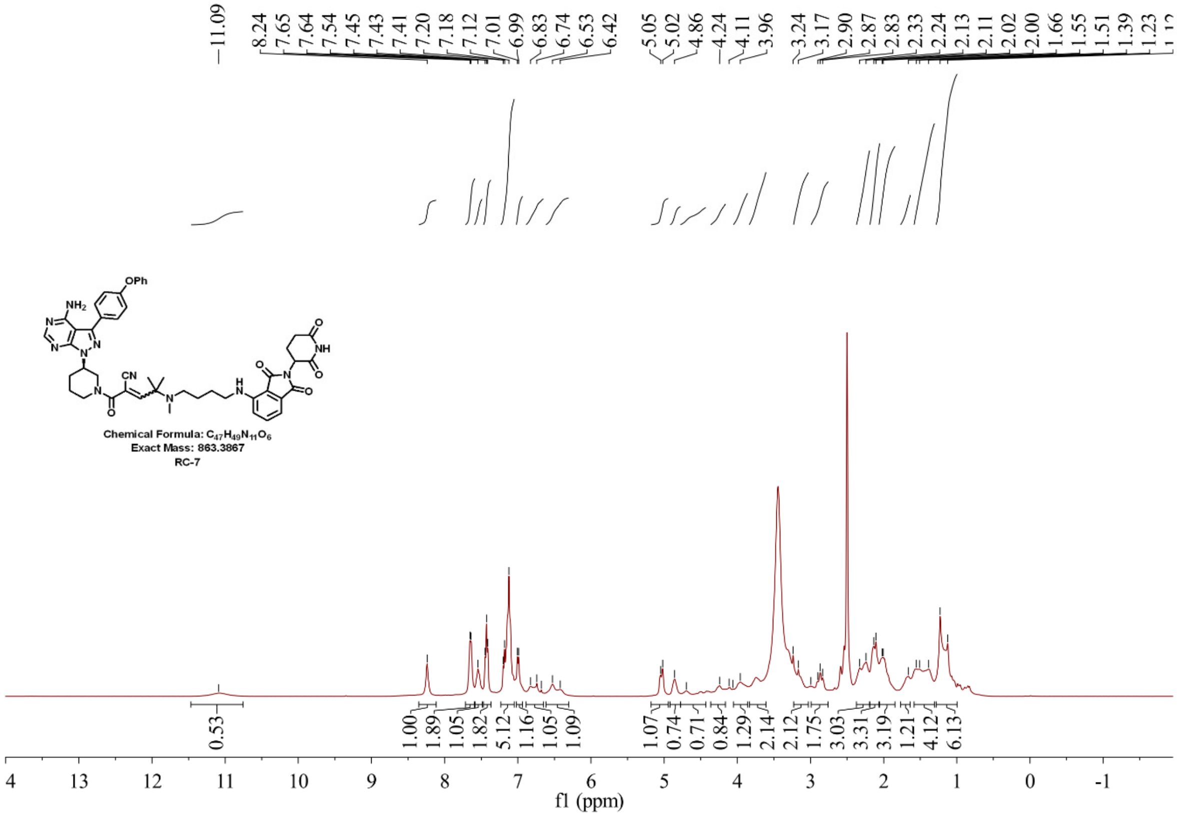


**Supplementary Figure 23. $^{1}$HNMR spectrum of RC-7**

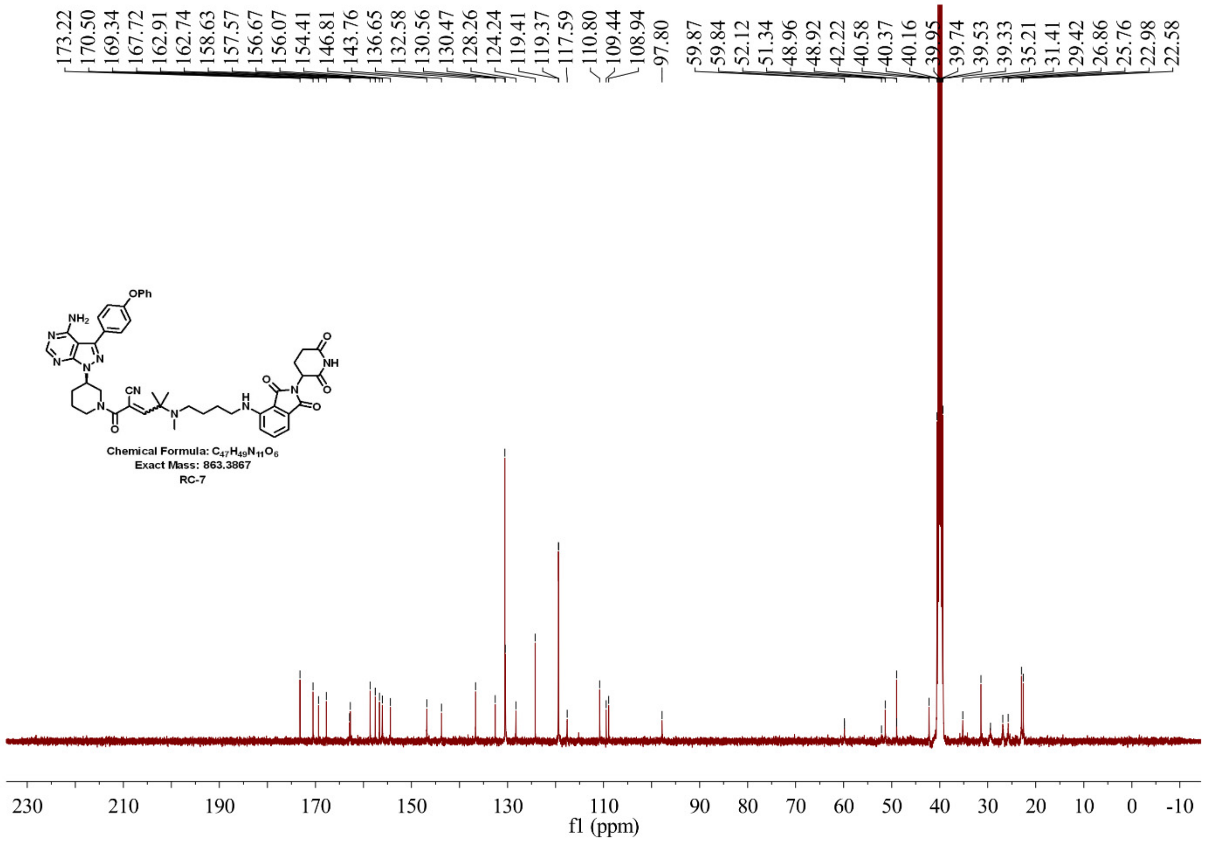


**Supplementary Figure 24. $^{13}$CNMR spectrum of RC-7**

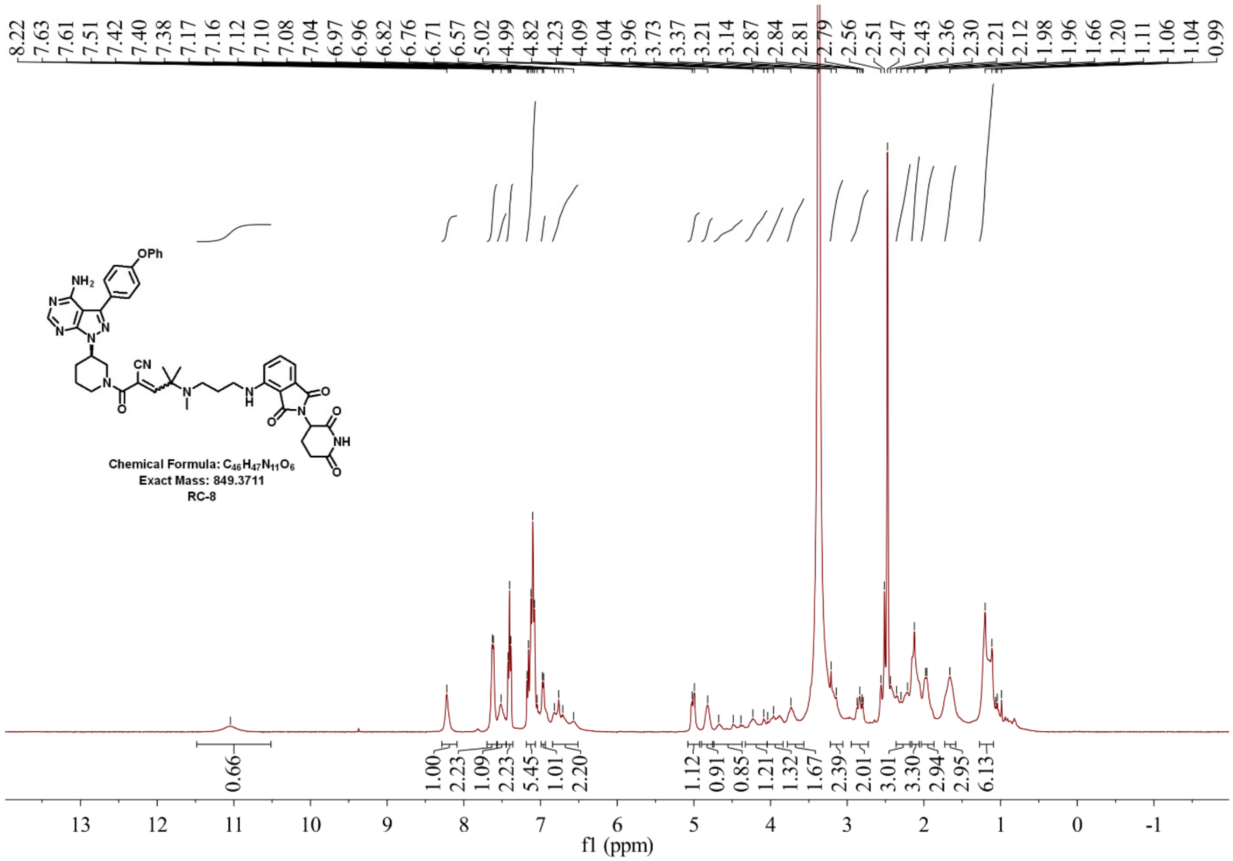


**Supplementary Figure 25. $^{1}$HNMR spectrum of RC-8**

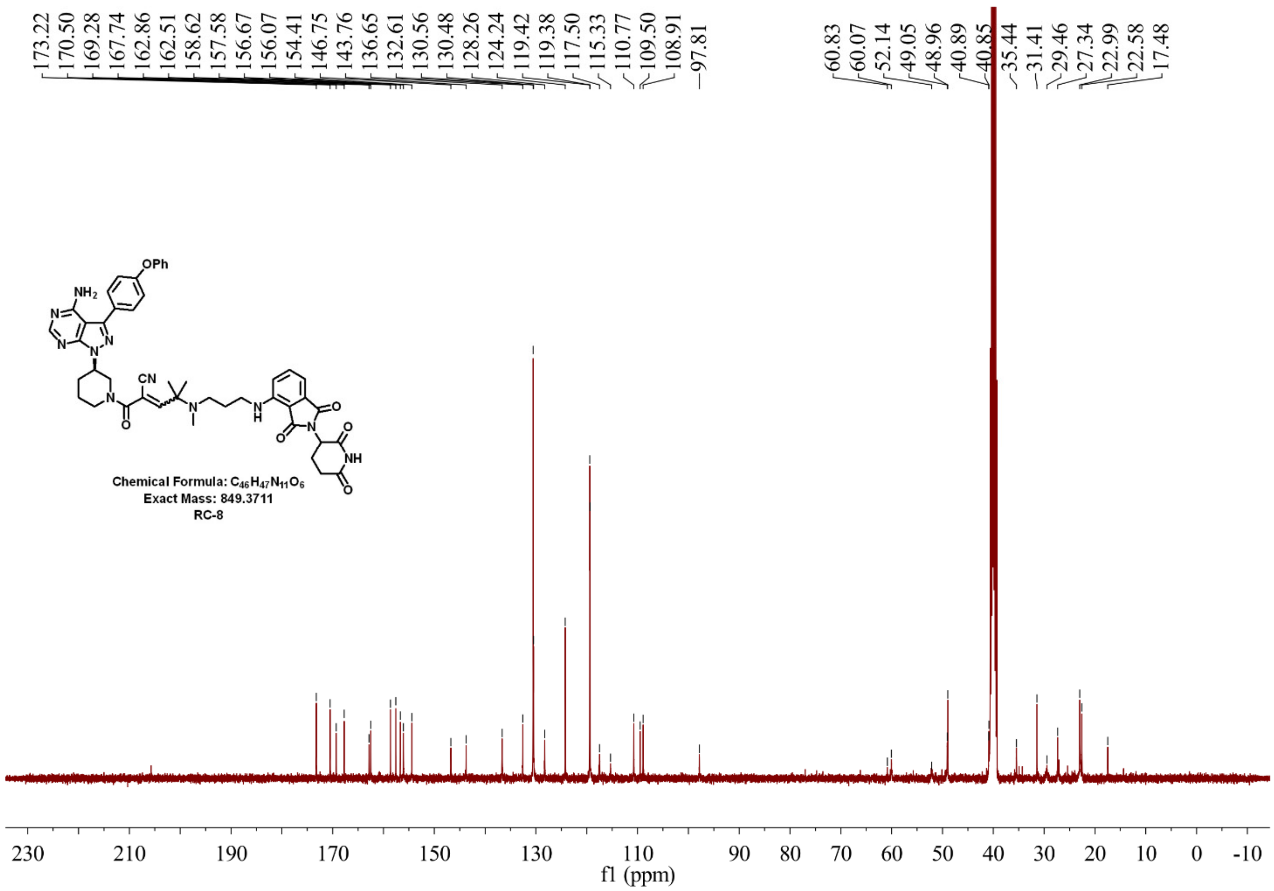


**Supplementary Figure 26. [13]CNMR spectrum of RC-8**

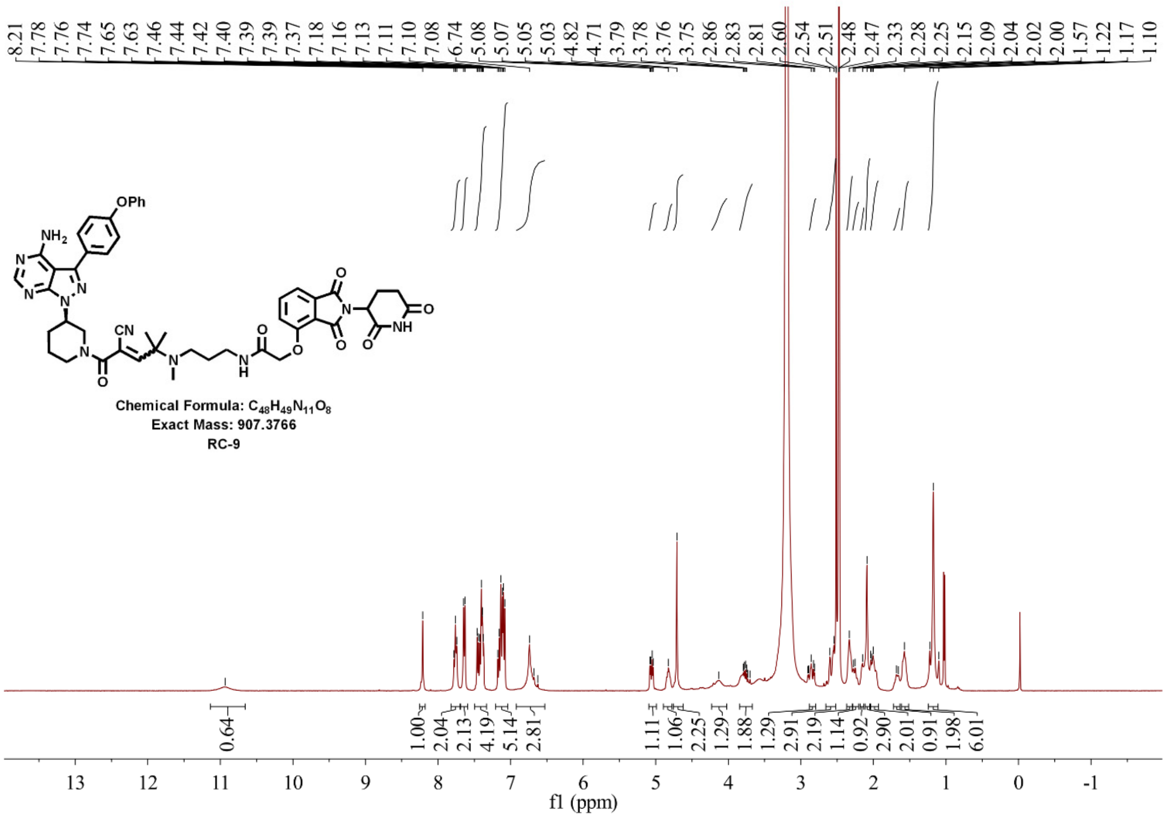


**Supplementary Figure 27. [1]HNMR spectrum of RC-9**

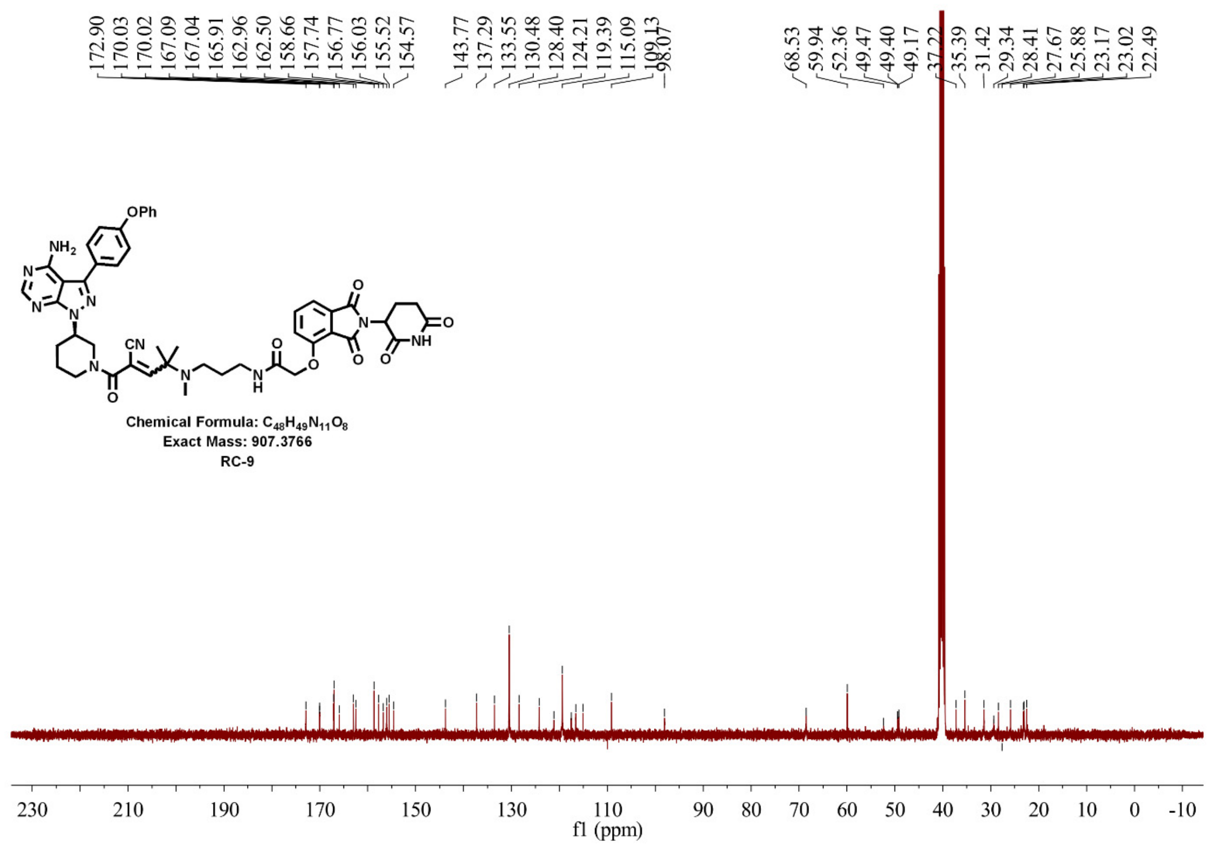


**Supplementary Figure 28. [13]CNMR spectrum of RC-9**

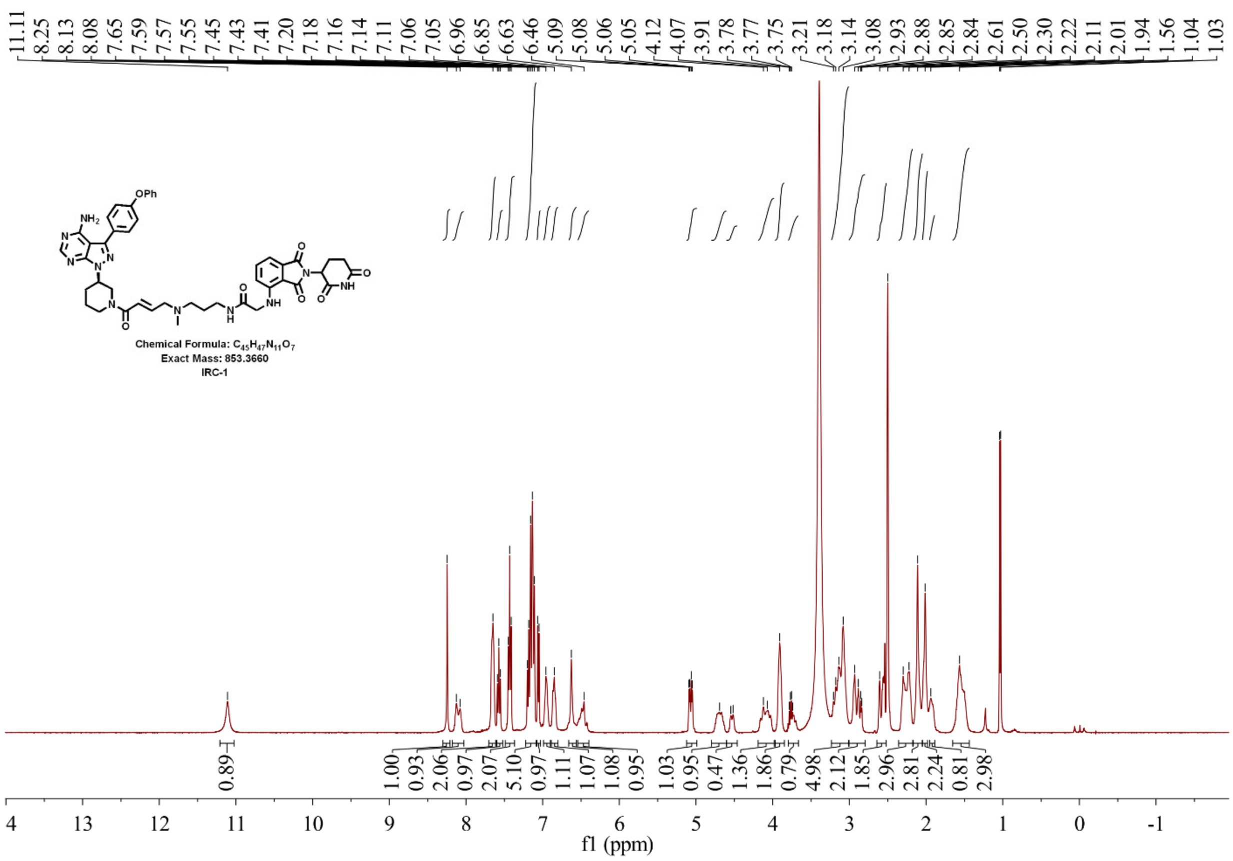


**Supplementary Figure 29. [1]HNMR spectrum of IRC-1**

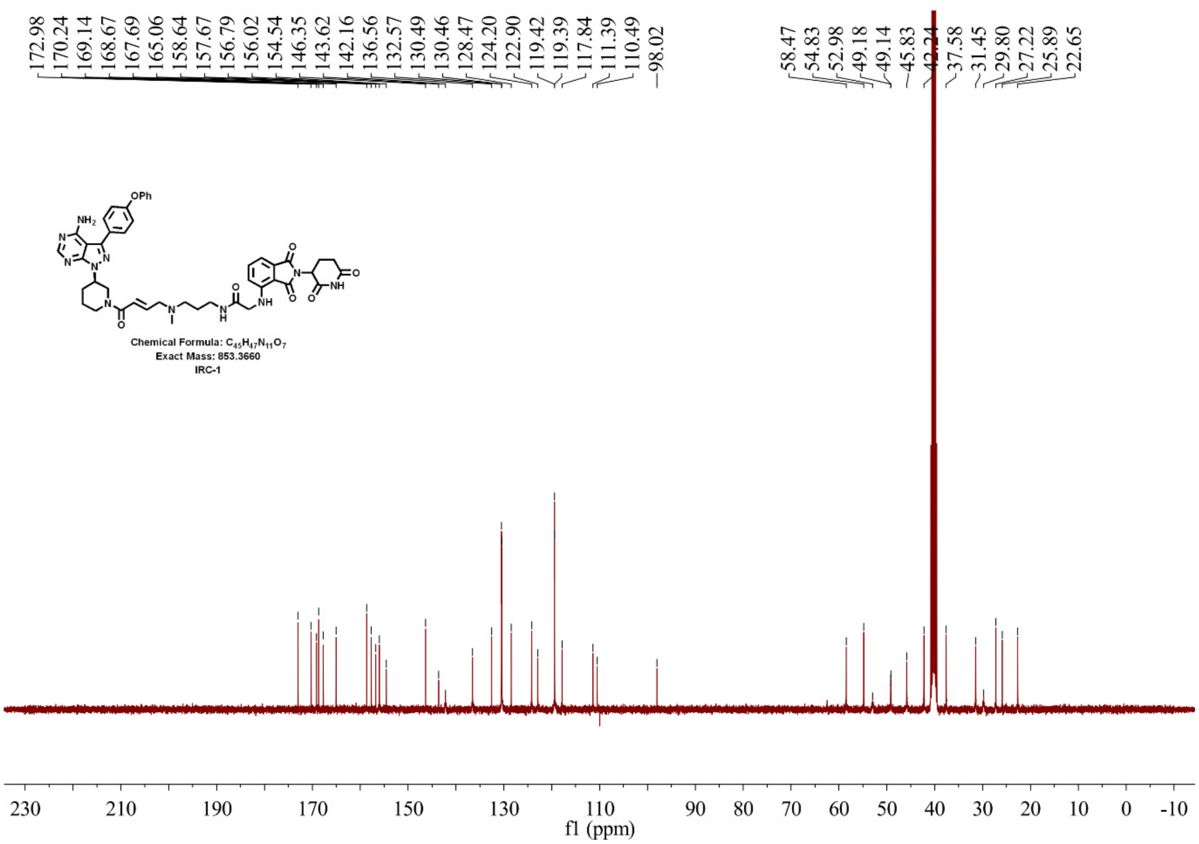


**Supplementary Figure 30. [13]CNMR spectrum of IRC-1**

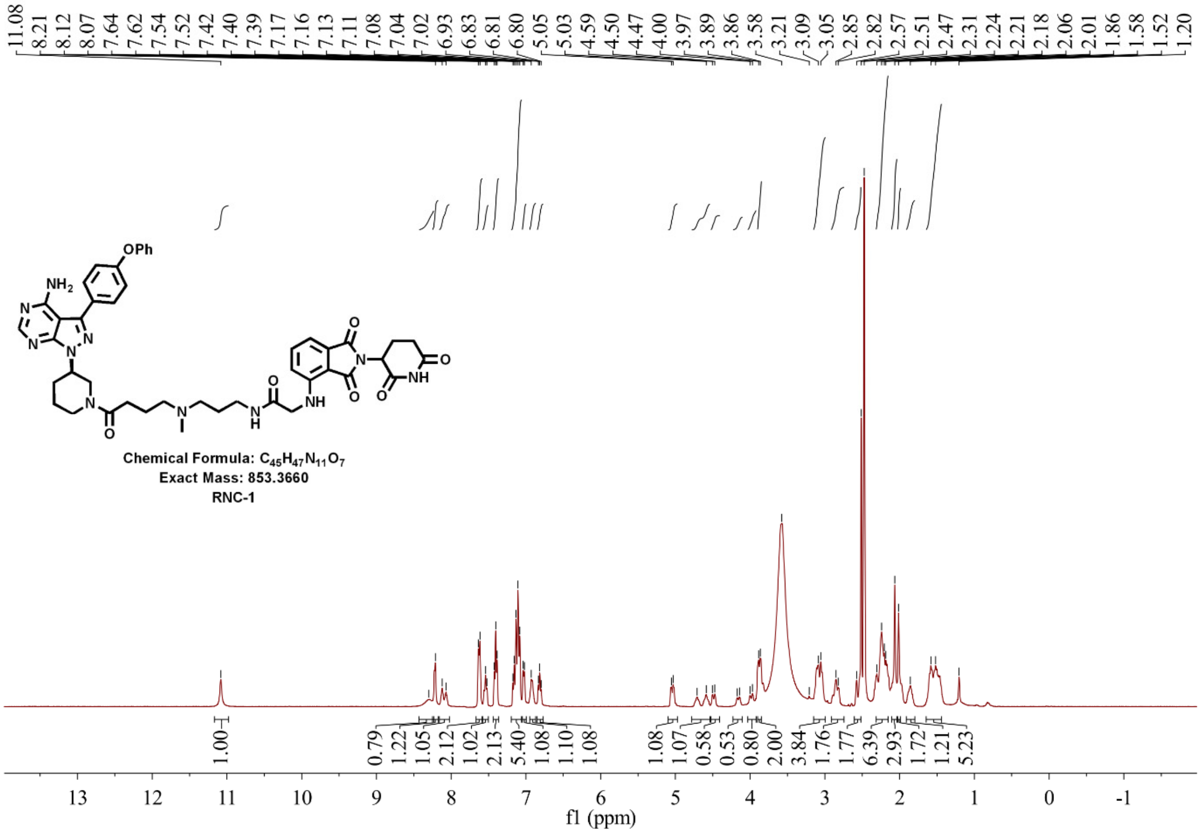


**Supplementary Figure 31. [1]HNMR spectrum of RNC-1**

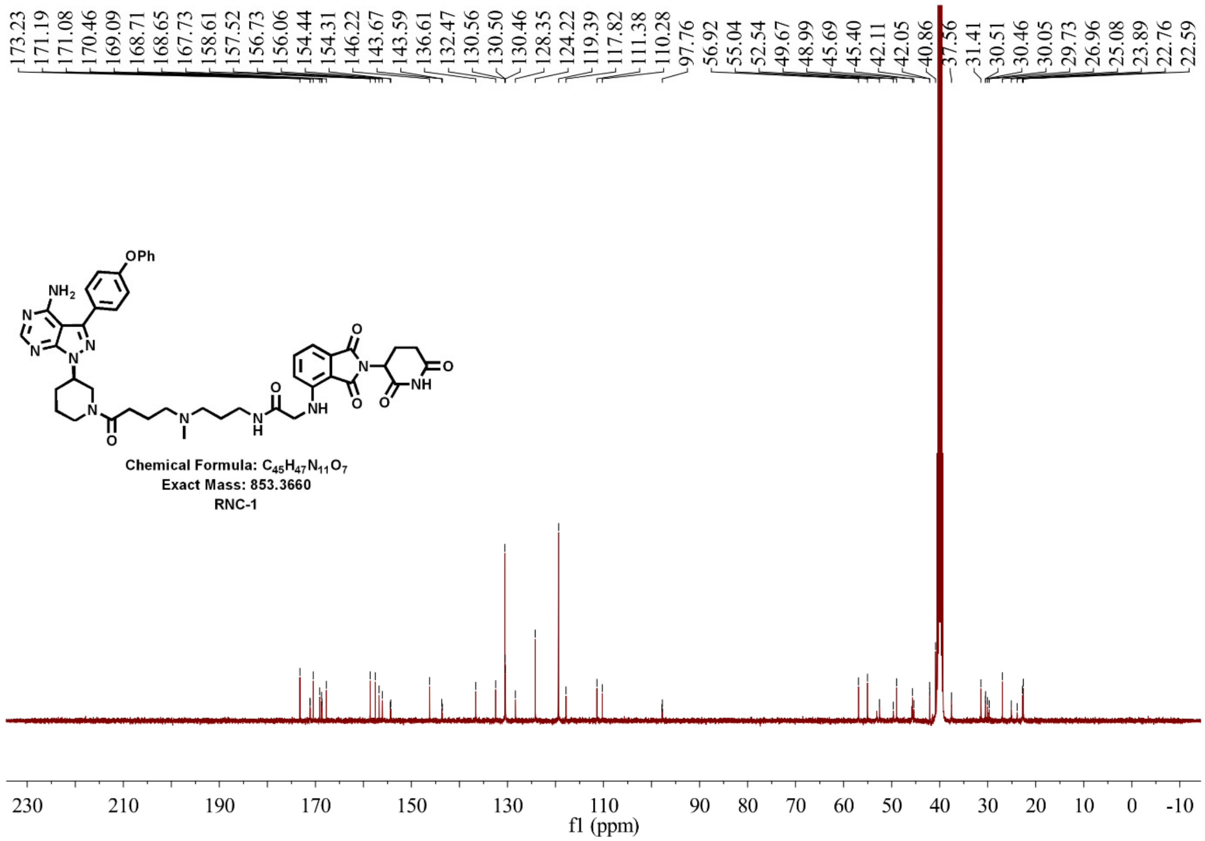


**Supplementary Figure 32. $^{13}$CNMR spectrum of RNC-1**

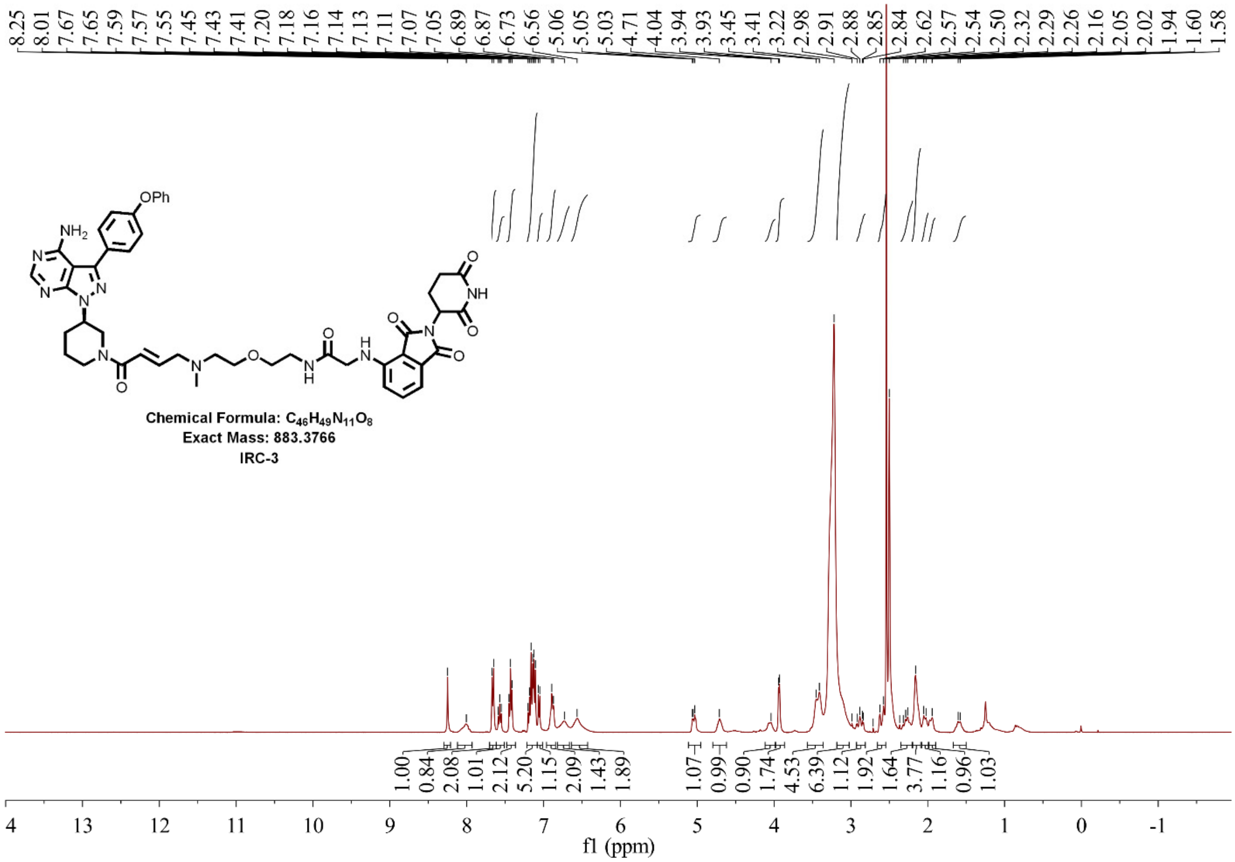


**Supplementary Figure 33. $^{1}$HNMR spectrum of IRC-3**

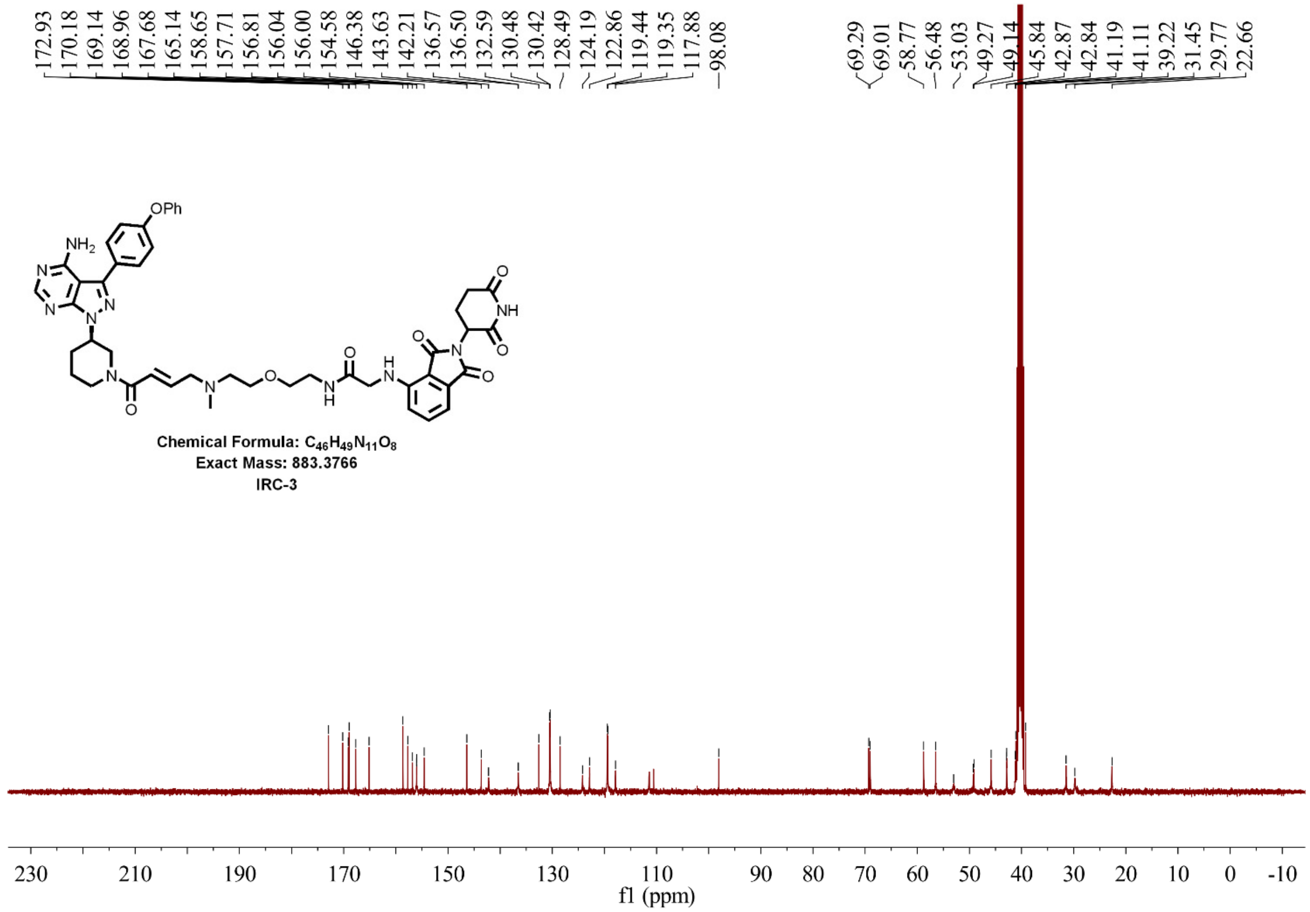


**Supplementary Figure 34. [13]CNMR spectrum of IRC-3**

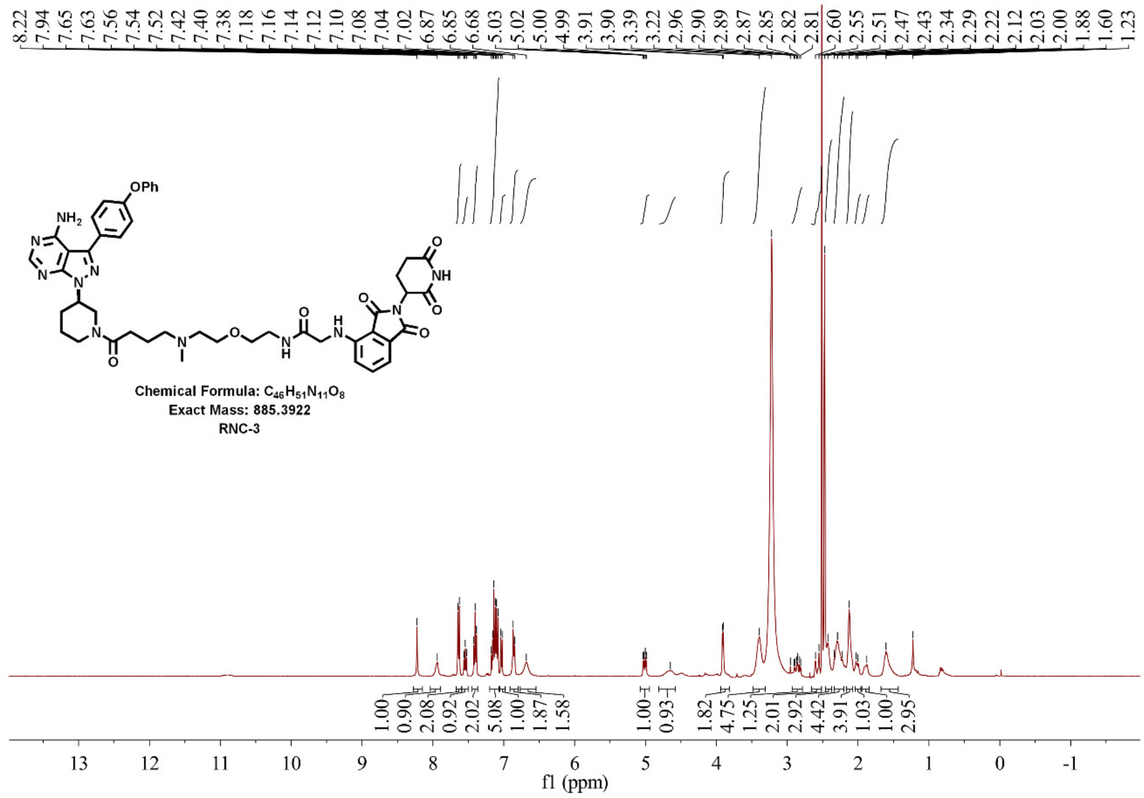


**Supplementary Figure 35. [1]HNMR spectrum of RNC-3**

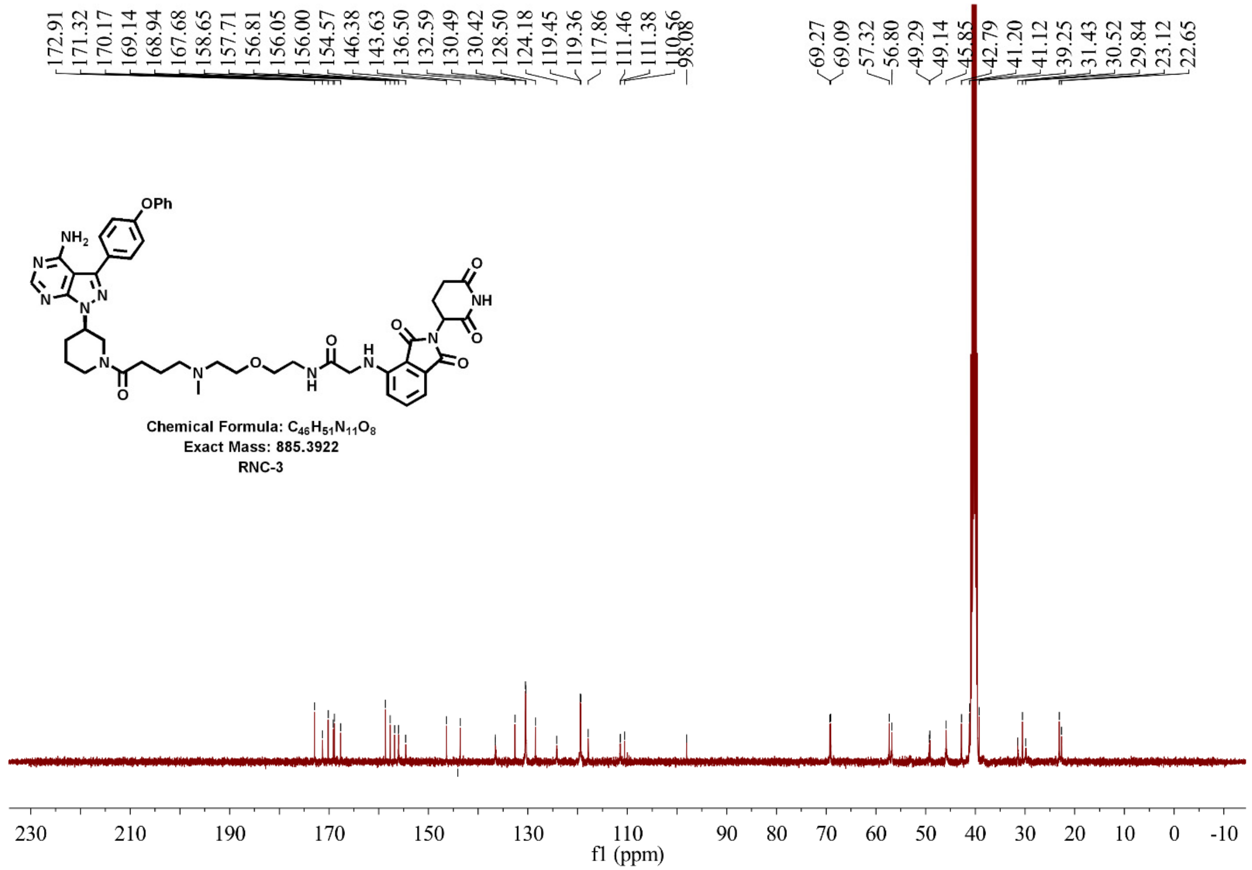

**Supplementary Figure 36. $^{13}$CNMR spectrum of RNC-3**

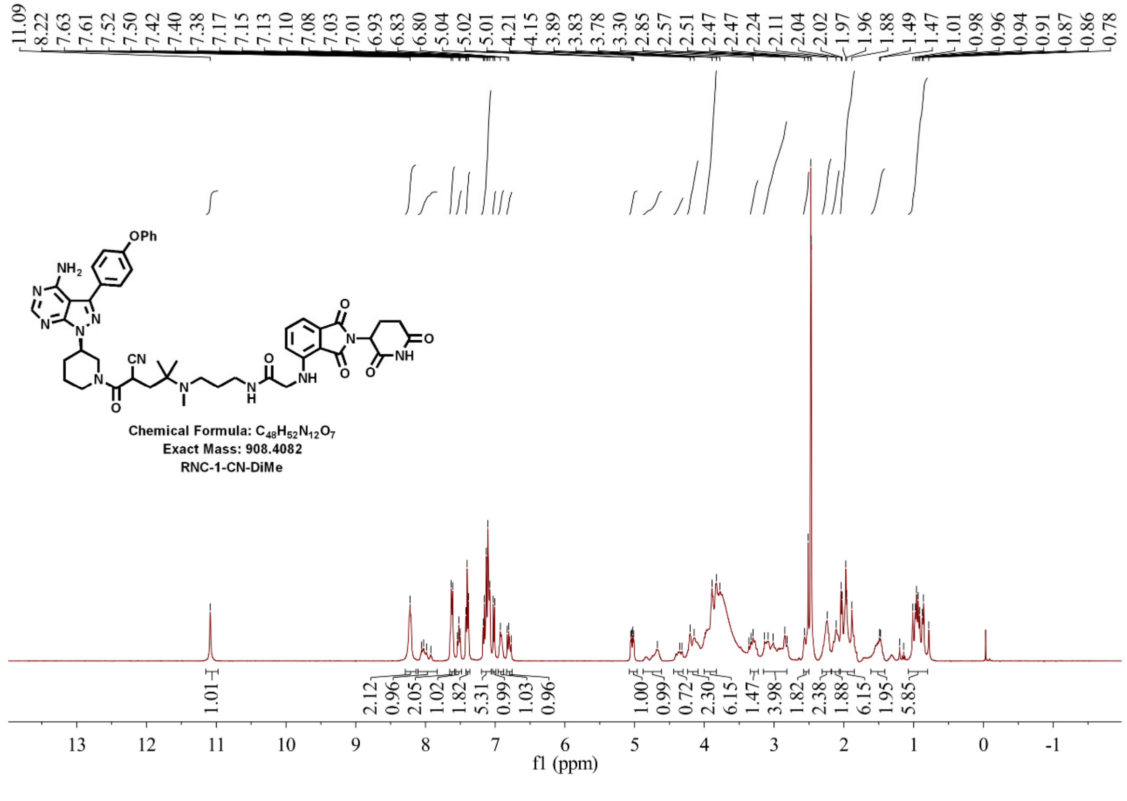

**Supplementary Figure 37. $^{1}$HNMR spectrum of RNC-1-CN-DiMe**

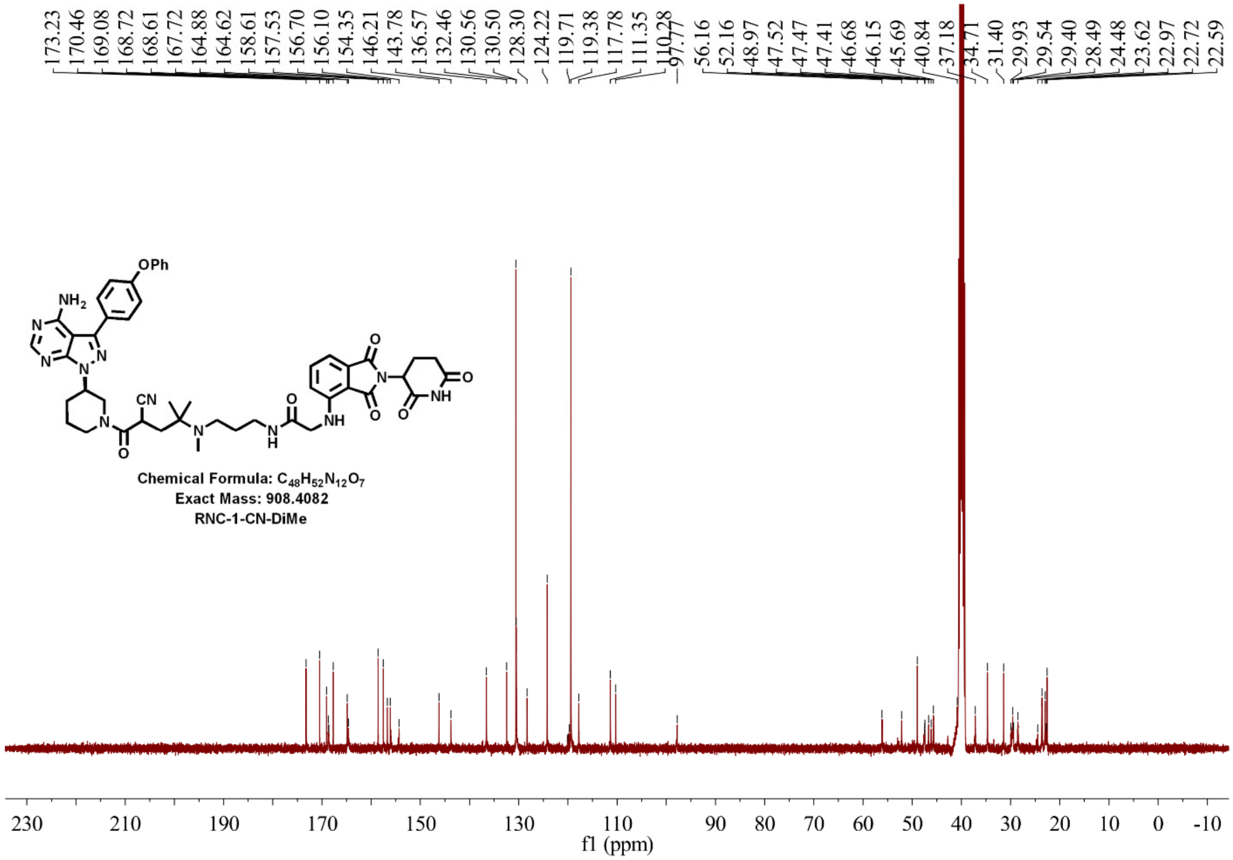


**Supplementary Figure 38. [1]CNMR spectrum of RNC-1-CN-DiMe**

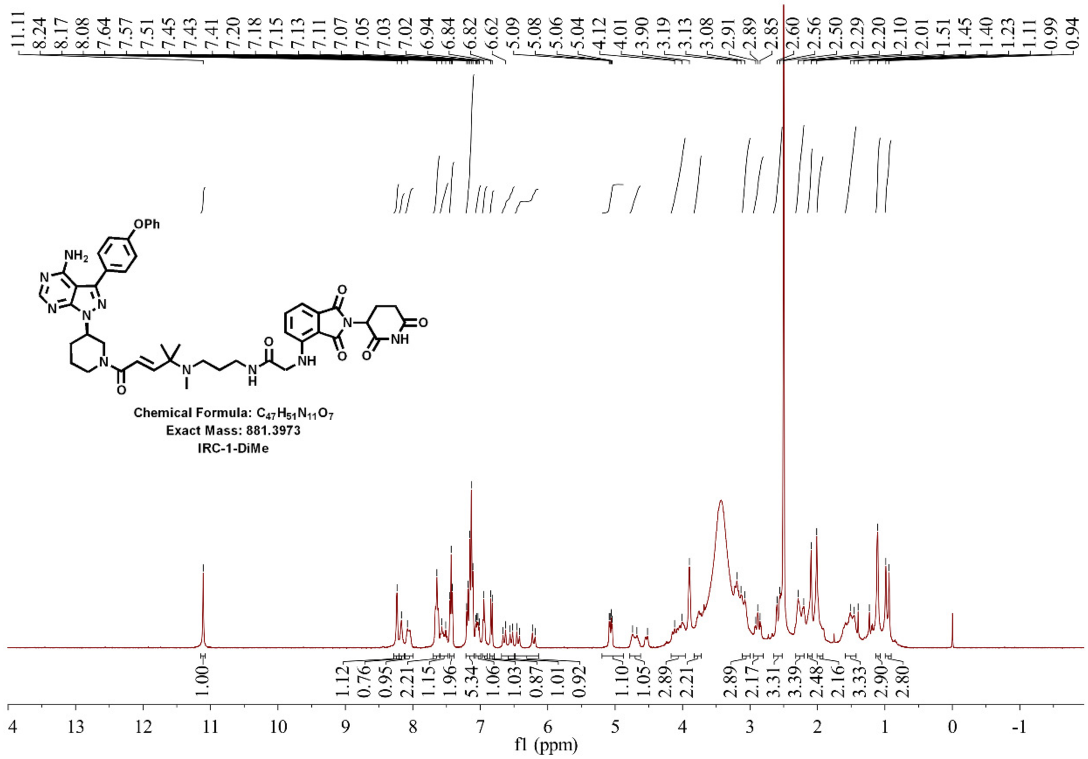


**Supplementary Figure 39. [1]HNMR spectrum of IRC-1-DiMe**

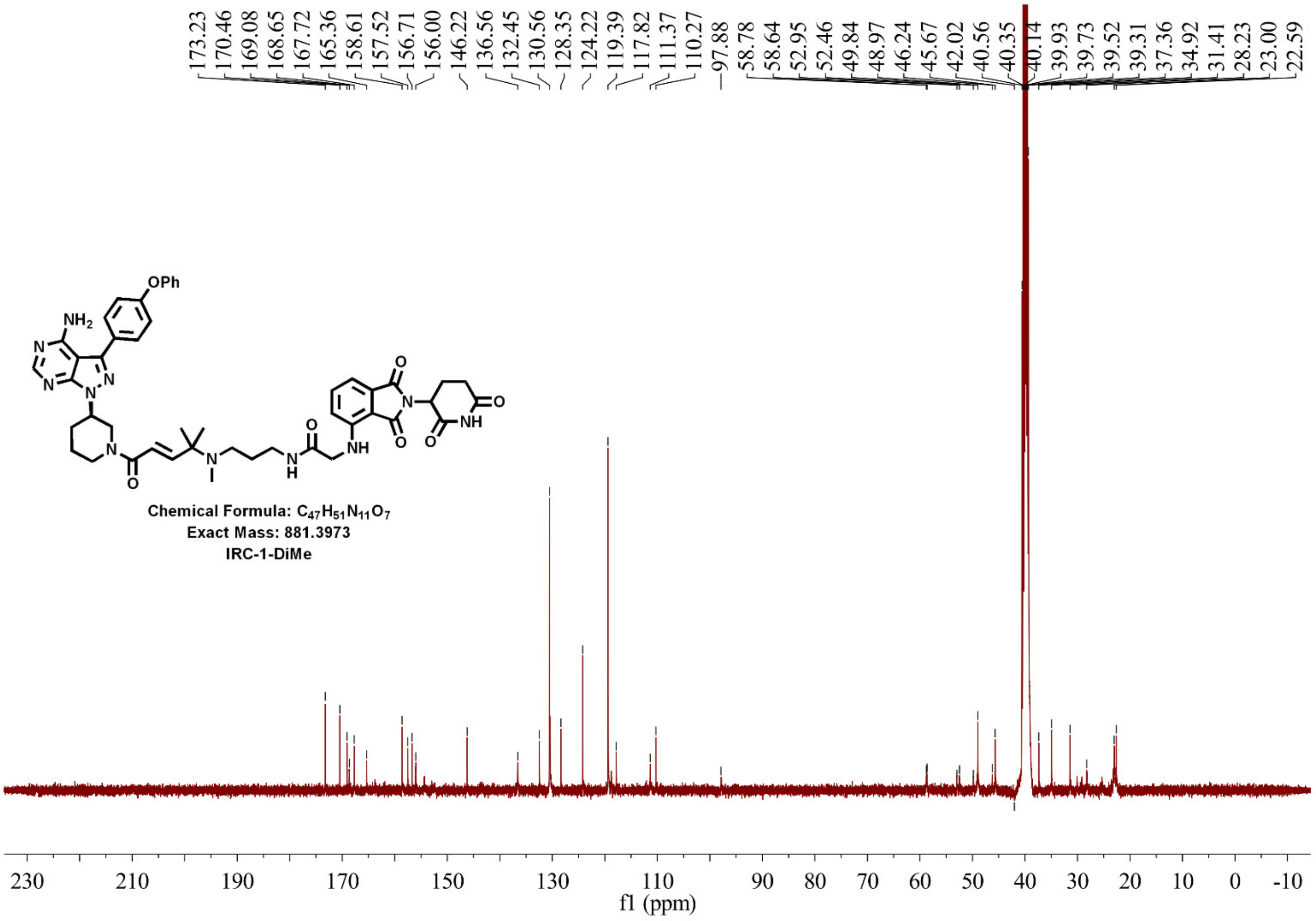


**Supplementary Figure 40. $^{13}$CNMR spectrum of IRC-1-DiMe**

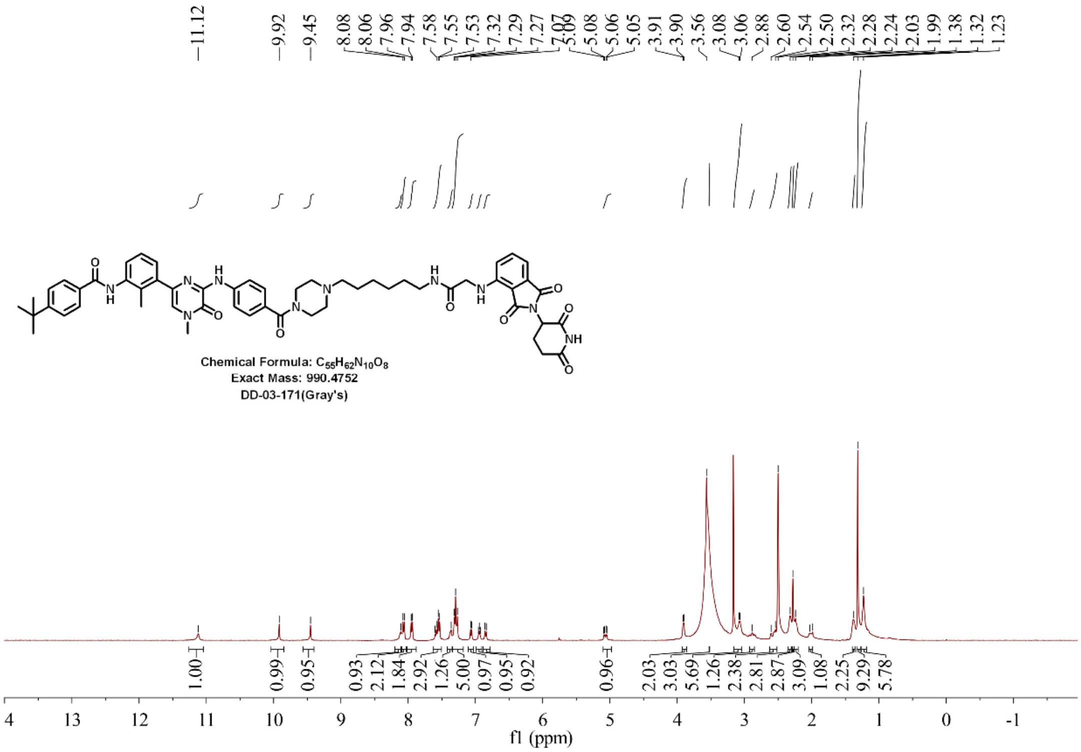


**Supplementary Figure 41. $^{1}$HNMR spectrum of DD-03-171**

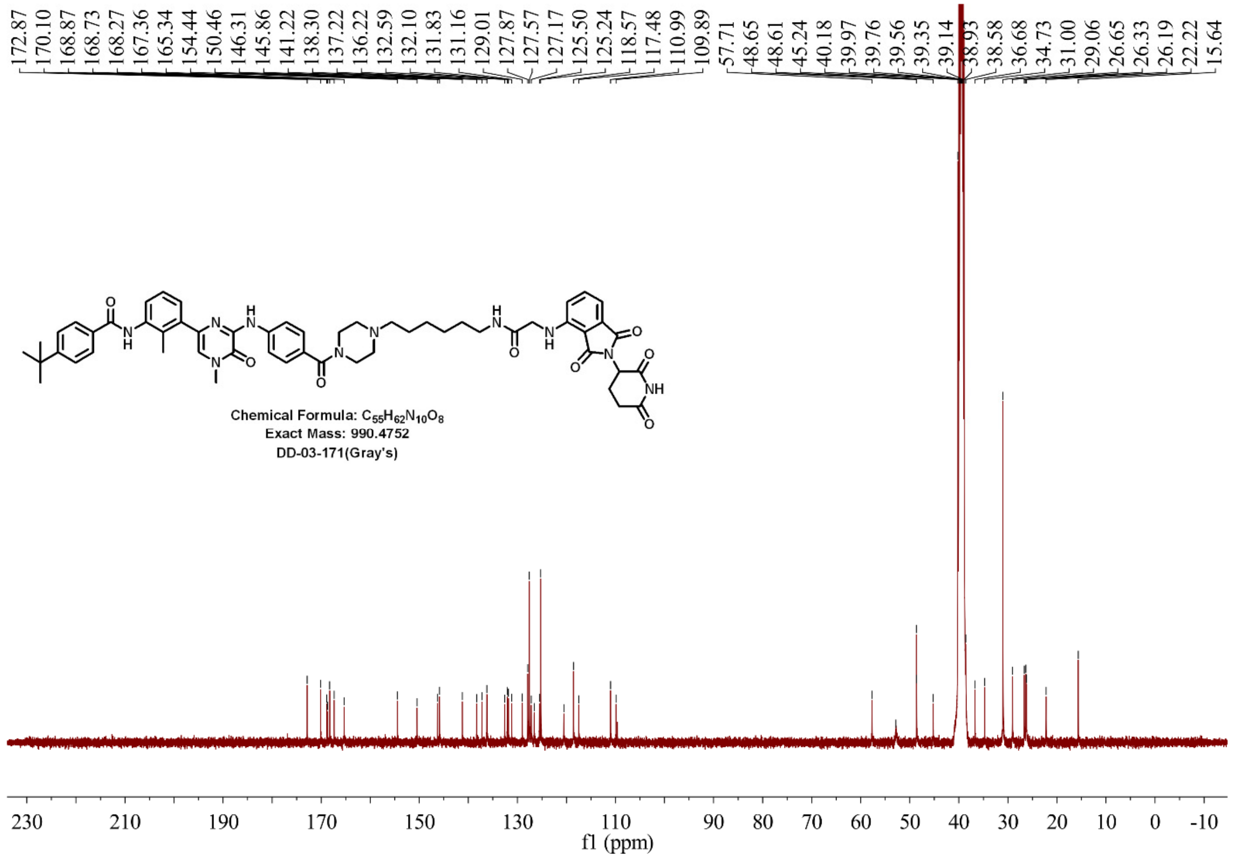


**Supplementary Figure 42. $^{13}$CNMR spectrum of DD-03-171**

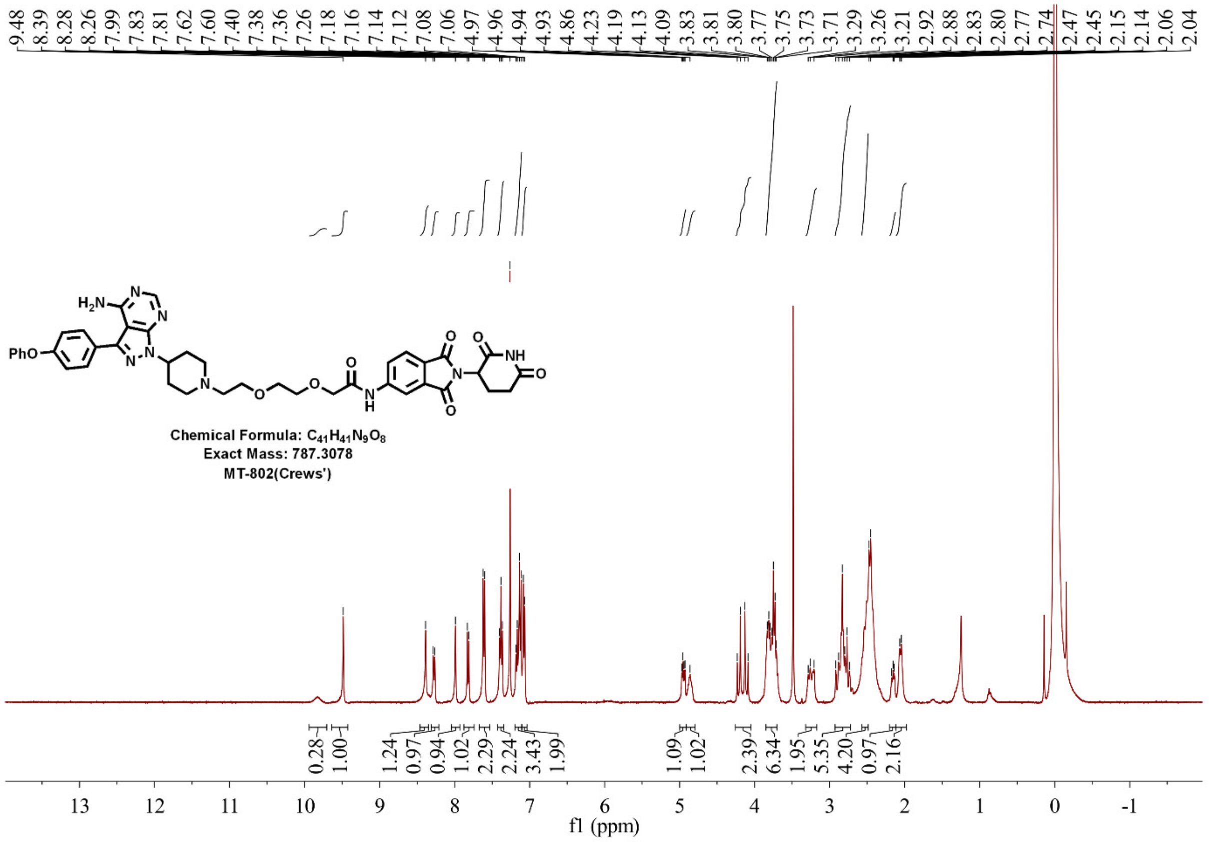


**Supplementary Figure 43. $^{1}$HNMR spectrum of MT-802**

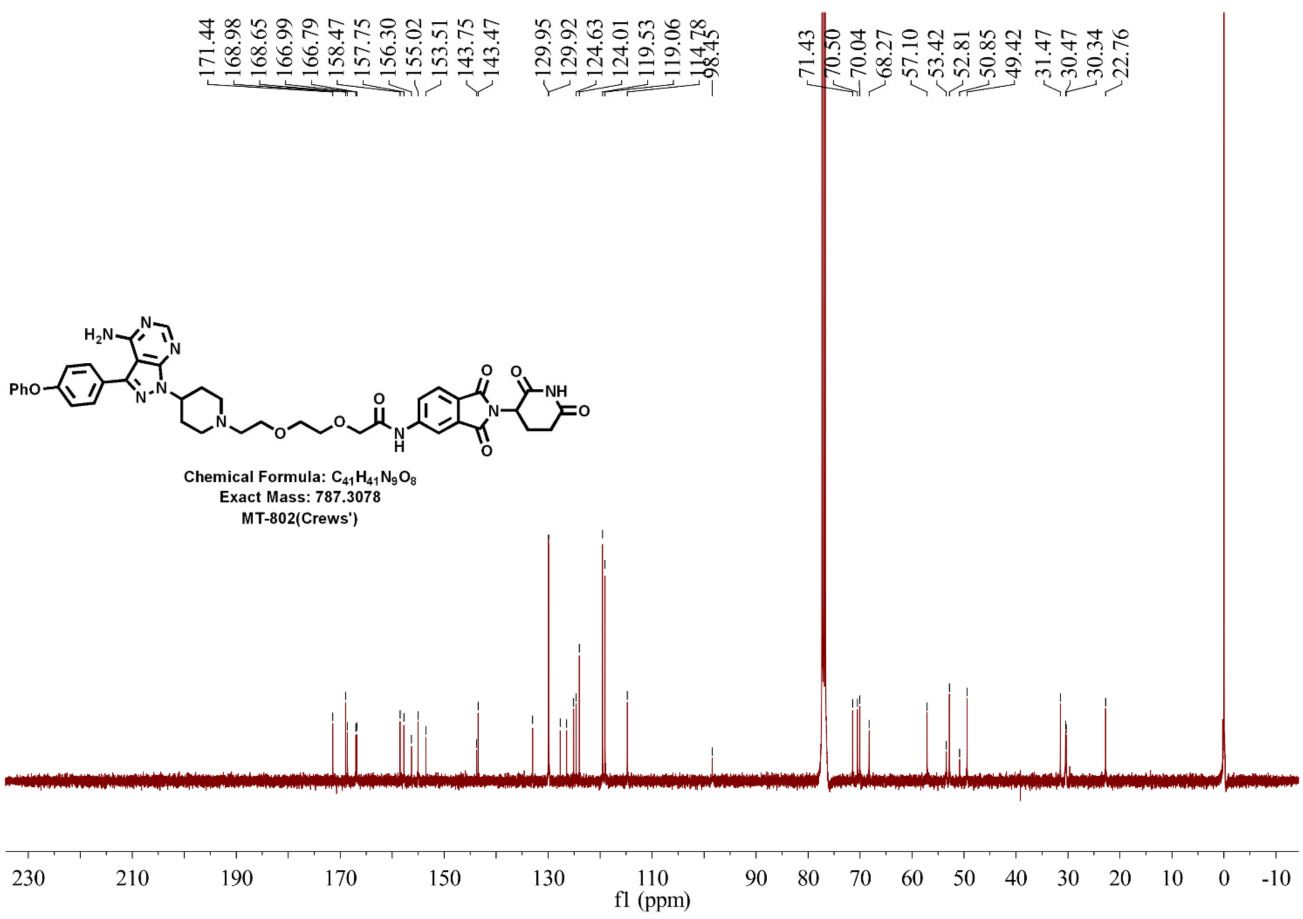


**Supplementary Figure 44. 13CNMR spectrum of MT-802**

**Supplementary Table 1. The calculated chemical properties for RC-1, IRC-1 and RNC-1 by ChemDraw**

| Compound | CLogP | tPSA |
|---|---|---|
| RC-1 | 4.26 | 247.6 |
| IRC-1 | 4.71 | 223.8 |
| RNC-1 | 4.55 | 223.8 |
| RNC-1-CN-DiMe | 3.81 | 247.6 |
| IRC-1-DiMe | 5.42 | 223.8 |

**Supplementary Table 2. $IC_{50}$ values of ibrutinib, RC-1, RC-Ctrl, and RC-1-Me in MCL cell lines**

| Compound | Mino | Jeko-1 | Rec-1 | Maver-1 |
|---|---|---|---|---|
| Ibrutinib | 3.8 | 0.3 | 6.2 | 5.4 |
| RC-Ctrl | >5 | 0.6 | >5 | >5 |
| RC-1-Me | 3.3 | 0.6 | >5 | 4.2 |
| RC-1 | 0.4 | 0.2 | 4.1 | 3.3 |

Triplicates were performed. Source data are provided as a Source Data file.

**Supplementary Table 3. The effective permeability coefficients ($P_e$) of control compounds and PROTACs**

| Compound ID | Replicate | -Log $P_e$ | Recovery% |
|---|---|---|---|
| Methotrexate | Replicate 1 | >8.99* | 104.72 |
| | Replicate 2 | >9.02* | 103.56 |
| | Replicate 3 | >9.00* | 99.70 |
| | Mean | >9.00* | 102.66 |
| Testosterone | Replicate 1 | 4.30 | 95.84 |
| | Replicate 2 | 4.51 | 88.71 |
| | Replicate 3 | 4.29 | 88.88 |
| | Mean | 4.37 | 91.14 |
| RC-1 | Replicate 1 | >8.19* | 25.10 |
| | Replicate 2 | >8.15* | 23.52 |
| | Replicate 3 | >8.12* | 23.40 |
| | Mean | >8.16* | 24.01 |
| RNC-1 | Replicate 1 | >8.70* | 30.03 |
| | Replicate 2 | >8.75* | 33.51 |
| | Replicate 3 | >8.59* | 24.91 |
| | Mean | >8.68* | 29.48 |
| IRC-1 | Replicate 1 | >7.61* | 17.38 |
| | Replicate 2 | >7.72* | 21.75 |
| | Replicate 3 | >7.51* | 15.06 |
| | Mean | >7.61* | 18.06 |

*Compound could not be detected in the receptor side, cut off values were calculated by LLOD. For more information see Supplementary Data 3.

**Supplementary Table 4. Quantification of PROTACs cellular concentration by LC-MS**

| Compound ID | Replicate | Cellular concentration (nmol/million cells) | | | | | | |
|---|---|---|---|---|---|---|---|---|
| | | 5 min | 15 min | 30 min | 60 min | 90 min | 120 min | 240 min |
| RC-1 | Replicate 1 | 0.0607 | 0.0487 | 0.0474 | 0.0471 | 0.0436 | 0.0460 | 0.0426 |
| | Replicate 2 | 0.0507 | 0.0414 | 0.0471 | 0.0390 | 0.0454 | 0.0460 | 0.0421 |
| | Replicate 3 | 0.0451 | 0.0451 | 0.0447 | 0.0480 | 0.0405 | 0.0513 | 0.0424 |
| | Mean | 0.0522 | 0.0451 | 0.0464 | 0.0447 | 0.0432 | 0.0478 | 0.0424 |
| RNC-1 | Replicate 1 | 0.0283 | 0.0286 | 0.0255 | 0.0183 | 0.0195 | 0.0166 | 0.0161 |
| | Replicate 2 | 0.0378 | 0.0292 | 0.0275 | 0.0188 | 0.0181 | 0.0169 | 0.0163 |
| | Replicate 3 | 0.0313 | 0.0308 | 0.0287 | 0.0180 | 0.0187 | 0.0199 | 0.0155 |
| | Mean | 0.0325 | 0.0295 | 0.0272 | 0.0183 | 0.0188 | 0.0178 | 0.0160 |
| IRC-1 | Replicate 1 | 0.0285 | 0.0307 | 0.0259 | 0.0274 | 0.0275 | 0.0239 | 0.0228 |
| | Replicate 2 | 0.0325 | 0.0322 | 0.0296 | 0.0266 | 0.0290 | 0.0281 | 0.0239 |
| | Replicate 3 | 0.0332 | 0.0305 | 0.0288 | 0.0240 | 0.0295 | 0.0261 | 0.0246 |
| | Mean | 0.0314 | 0.0311 | 0.0281 | 0.0260 | 0.0287 | 0.0261 | 0.0238 |

For more information see Supplementary Data 4.
